# Discovering Cognitive Strategies with Tiny Recurrent Neural Networks

**DOI:** 10.1101/2023.04.12.536629

**Authors:** Li Ji-An, Marcus K. Benna, Marcelo G. Mattar

## Abstract

Normative modeling frameworks such as Bayesian inference and reinforcement learning provide valuable insights into the fundamental principles governing adaptive behavior. While these frameworks are valued for their simplicity and interpretability, their reliance on few parameters often limits their ability to capture realistic biological behavior, leading to cycles of handcrafted adjustments that are prone to research subjectivity. Here, we present a novel modeling approach leveraging recurrent neural networks to discover the cognitive algorithms governing biological decision-making. We show that neural networks with just 1-4 units often outperform classical cognitive models and match larger neural networks in predicting the choices of individual animals and humans across six well-studied reward learning tasks. Critically, we then interpret the trained networks using dynamical systems concepts, enabling a unified comparison of cognitive models and revealing detailed mechanisms underlying choice behavior. Our approach also estimates the dimensionality of behavior and offers insights into algorithms implemented by AI agents trained in a meta-reinforcement learning setting. Overall, we present a systematic approach for discovering interpretable cognitive strategies in decision-making, offering insights into neural mechanisms and a foundation for studying both healthy and dysfunctional cognition.

## Introduction

Understanding the neural basis of adaptive behavior is a fundamental goal of neuroscience. Researchers have long strived to develop computational models that encapsulate the complexities of learning and decision-making, from early symbolic models^1^ to connectionist approaches^2^. Normative frameworks such as Bayesian inference^3,4^ and reinforcement learning^5–8^ have been particularly influential for their ability to elucidate the fundamental principles governing adaptive behavior. These cognitive models formalize how agents accumulate and apply knowledge from environmental interactions to make decisions, a process thought to be carried out by neural circuits in the prefrontal cortex and striatum^9–13^. A key advantage of these models is their simplicity, as they typically have few parameters and can be easily augmented with additional assumptions such as forgetting, choice biases, perseveration, exploration, and capacity limitations^12,14^. While the simplicity and extensibility of these models are advantageous in many ways, this approach is prone to bias and researcher subjectivity, potentially leading to incorrect or incomplete characterizations of behavior^15^.

An alternative modeling framework employs artificial neural networks, a class of computational models consisting of interconnected neuron-like units that can express a wide range of functions. Compared to classical cognitive models, neural networks impose fewer structural assumptions, require less handcrafting, and provide a more flexible framework for modeling behavior and neural activity^16,17^. A common approach involves adjusting the network parameters to produce optimal behavior in a given task. This approach has been used in neuroscience to explain the neural activity associated with vision,^18,19^, spatial navigation^20–22^, learning, decision-making, and planning^23–27^. An alternative approach involves adjusting the network parameters to predict some patterns of biological behavior. Due to the large number of parameters used in most neural networks, this approach often results in highly accurate predictions of future behavior^28–31^. However, this increased flexibility leads to difficulties in interpreting the fitted models, hindering the identification of cognitive and neural mechanisms underlying observable behavior.

Here, we present a novel modeling framework that combines the flexibility of neural networks with the interpretability of classical cognitive models. Our framework involves fitting recurrent neural networks (RNNs) to the behavior of individual subjects in reward learning tasks. The fitted RNNs describe how individuals accumulate knowledge by interacting with their environment and how this knowledge is applied to decision-making. In contrast to previous approaches, however, our framework uses very small RNNs, often composed of just 1-4 units, which greatly facilitates their interpretation. In contrast to previous approaches, however, our framework uses very small RNNs, often composed of just 1-4 units, which greatly facilitates their interpretation. We show that these tiny RNNs outperform classical cognitive models of equal dimensionality in predicting the choices of humans and animals, across eight datasets. We then interpret the fitted networks as discrete dynamical systems, leveraging concepts and visualizations from dynamical systems theory to investigate their dynamics and to provide a direct comparison with alternative models. We show that this framework reveals several novel behavioral patterns overlooked by classical models, including variable learning rates, a new type of “reward-induced indifference” effect, previously unrecognized forms of value updating, state-dependent choice perseveration, and choice biases. Overall, our results show that tiny RNNs can not only predict behavior better than previous models, but also provide deeper insights into cognitive mechanisms, addressing both the interpretability challenges of larger neural networks and the subjectivity of classical models.

## Results

### Task description and model overview

We studied biological behavior in six reward learning tasks widely used in neuroscience and psychology, three performed by animals and three by humans (Fig. 1, 2). These tasks capture fundamental processes by which animals and humans learn to make decisions through environmental interactions, which our modeling framework aims to describe. We first present our results for the animal tasks: a reversal learning task^11^, a two-stage task^13^, and a transition-reversal two-stage task^12^ (Fig. 1d). In all of these tasks, each trial consists of a choice between actions *A*_1_ and *A*_2_, resulting in either state *S*_1_ or *S*_2_. Each state is associated with a probability of receiving a binary reward, and these probabilities switch unpredictably in moments called “reversals”. The subject’s goal is to choose the action most likely to yield a reward. In the reversal learning task, each action leads deterministically to one state (e.g., *A*_1_ leads to *S*_1_ and *A*_2_ to *S*_2_). The two-stage task introduces probabilistic transitions, where each action can lead to either states with different probabilities (e.g., *A*_1_ leads to *S*_1_ with probability 0.8 and to *S*_2_ with probability 0.2). The transition-reversal two-stage task adds a stochastic reversal to the action-state transition probabilities (e.g., A1 leads to *S*_1_ and *S*_2_ with high and low probabilities, respectively, with these probabilities switching during a reversal). We analyzed data from two monkeys (Bartolo dataset^11^) and ten mice (Akam dataset^12^) performing the reversal learning task; from four rats (Miller dataset^13^) and ten mice (Akam dataset^12^) performing the two-stage task; and from seventeen mice (Akam dataset^12^) performing the transition-reversal two-stage task.

**Fig. 1.**
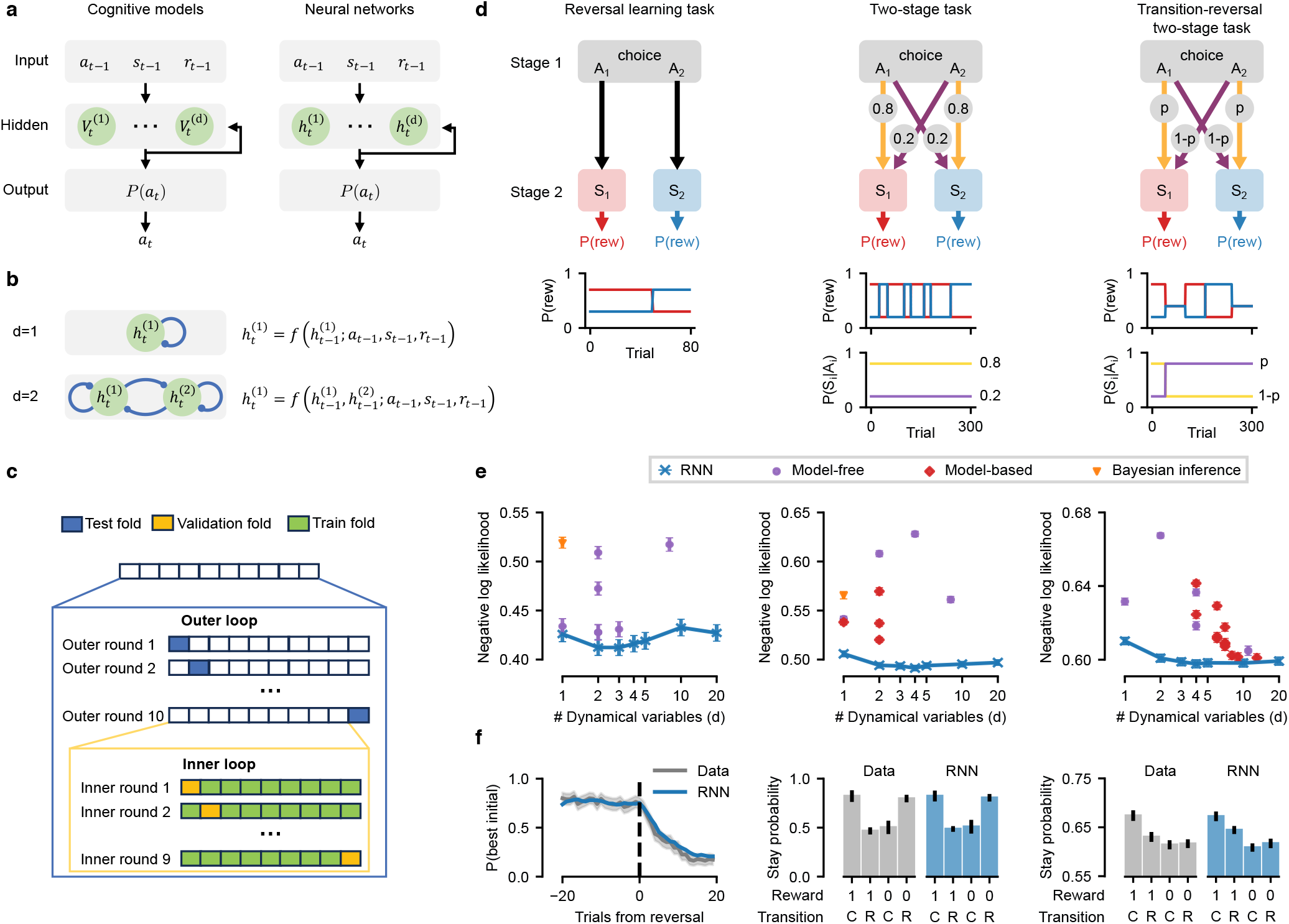
Model overview and model performance on animal tasks. **(a)** Cognitive models and neural networks have similar architectures. Inputs (previous action *a*_*t*−1_, state *s*_*t*−1_, reward *r*_*t*−1_) update *d* dynamical variables, which in turn determine output (action probability *P* (*a*_*t*_)) via softmax. Models are optimized to predict observed actions. **(b)** Recurrent layer examples: *d* = 1 (top) and *d* = 2 (bottom) units. **(c)** Nested cross-validation. In the outer loop, the whole dataset is split into ten folds of consecutive trials, with each round using one fold for testing and the other nine for the inner loop. Each inner loop round uses one fold for validation (e.g., hyperparameter tuning) and the other eight for training. **(d)** Task structures: Subjects choose action *A*_1_ or *A*_2_ at the choice state, transitioning into one of two second-stage states, *S*_1_ or *S*_2_, which probabilistically yield a reward. Reward probabilities change over time. In the transition-reversal two-stage task, the action-state transition probabilities also change over time. **(e)** Model performance (cross-validated trial-averaged negative log-likelihood; lower is better) vs. number of dynamical variables *d*. Tiny RNNs outperform classical models in all tasks. Identical markers within a plot represent different variants of a model class. Error bars: SEM across rounds, averaged over individuals. *Left:* In the reversal learning task with monkeys, the best performing model is a two-unit RNN. *Center:* In the two-stage task with rats, the best-performing model is a two-unit RNN. *Right:* In the transition-reversal two-stage task with mice, the best performing model is a four-unit RNN. **(f)** RNN reproduction of behavioral metrics: *Left:* Probability of choosing high-reward action pre-reversal (reversal learning, *d* = 2 RNN). Shaded region (left): 95% CI across blocks. *Center:* Probability of taking the same action (stay probability) following each trial type (two-stage task, *d* = 2 RNN). Transition C: common; R: rare; Error bars: cross-subject SEM. *Right:* Stay probabilities (transition-reversal task, *d* = *d*_*∗*_ GRU). Transition C: common; R: rare; Error bars: cross-subject SEM.

**Fig. 2.**
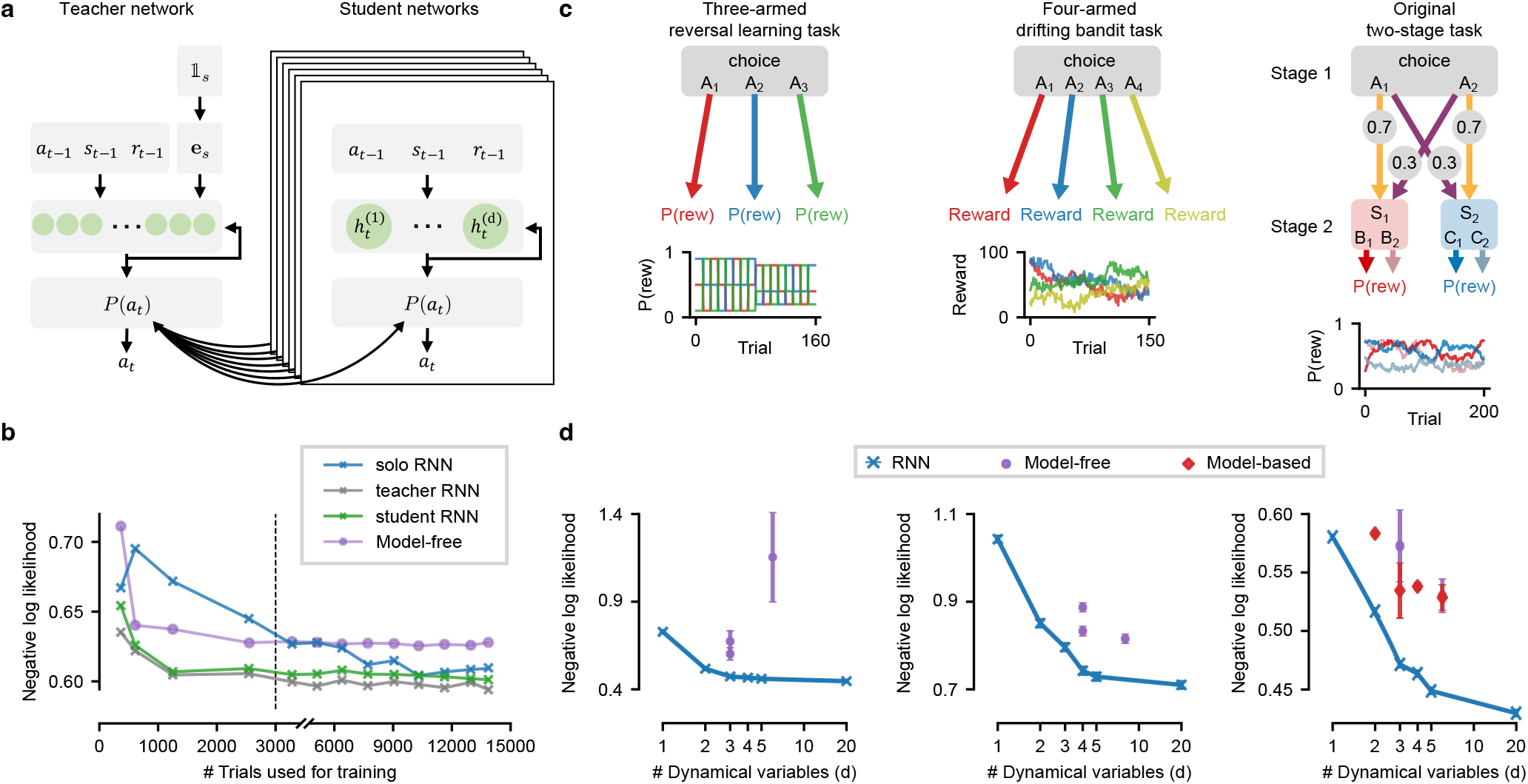
Model performance on human tasks using knowledge distillation. **(a)** Knowledge distillation framework: A large teacher network (TN) is trained on data from multiple subjects; the subject ID corresponding to each datapoint, provided as input, enables subject-specific embeddings **e**_*s*_ to be learned. A tiny student network (SN) is then trained on single-subject data to match TN’s output probabilities. **(b)** Model predictive performance for a representative mouse in the transition-reversal two-stage task, across varying dataset sizes. Student RNN outperforms the best model-free RL model for all dataset sizes. Note different x-axis scales for *<* 3000 and *>* 3000 trials. **(c)** Human tasks structures. *Left:* Three-armed reversal learning: Subjects choose between actions *A*_1_-*A*_3_, each associated with a reward probability that changes over time. *Center:* Four-armed drifting bandit: Subjects choose between actions *A*_1_-*A*_4_, with each associated with a 0-100 reward that fluctuates over time. *Right:* Original two-stage task: Subjects first choose action *A*_1_ or *A*_2_ at the choice state, transitioning into one of two second-stage states, *S*_1_ or *S*_2_. Subjects then choose action *B*_1_/*B*_2_ or *C*_1_/*C*_2_, each probabilistically yielding a reward. Reward probabilities change over time. **(d)** Model performance (cross-validated trial-averaged negative log-likelihood; lower is better) vs. number of dynamical variables *d*, averaged over subjects using interspersed split protocol. Tiny RNNs outperform classical models in all tasks. Identical markers within a plot represent different variants of a model class. Error bars: SEM across individuals. *Left:* Three-armed reversal learning. *Center:* Four-armed drifting bandit. *Right:* Original two-stage.

Our approach leverages RNN models to predict the choices of individual animals in these tasks and to interpret the underlying cognitive mechanisms. As a benchmark, we also implemented over 30 classical cognitive models previously used to describe behavior in these tasks, including Bayesian inference models, RL models, and many of their variants (see Methods). Cognitive models and RNNs share the same input-output structure (Fig. 1a). Inputs include the agent’s previous action *a*_*t*−1_, second stage state *s*_*t*−1_, and reward *r*_*t*−1_ (the current state is always the same “choice state”, and thus is not included as input). Inputs update the agent’s internal state, described by a set of *d* internal *dynamical variables* that summarize the agent’s prior experience (e.g., action values, belief states) to guide future actions. The model outputs are the probabilities Pr(*a*_*t*_ = *A*_*i*_) of executing each action *A*_*i*_, known as the agent’s behavioral policy.

Despite sharing the same input-output structure, each model employs distinct update rules for its dynamical variables, causing these variables to represent different latent variables and, consequently, leading to different behavioral policies. The classical cognitive models can be clustered into three families: model-free RL, model-based RL, and Bayesian inference. In the two RL model families, dynamical variables represent action values updated via RL algorithms: model-free RL updates these values directly from reward input, and model-based RL updates them indirectly through a state-transition model. In the Bayesian inference family, dynamical variables represent belief states updated via Bayesian inference. We also implemented various extensions of these models accounting for factors like choice perseveration or learned state-transition probabilities (see Methods). The RNN models, on the other hand, have fewer constraints in the update of dynamical variables. Each network unit corresponds to one dynamical variable that, through the network training procedure, comes to represent a possibly complex function of network inputs and of previous unit activities (Fig. 1b). These functions are determined by the network weights, a set of adjustable parameters that enables the model to learn diverse mappings from past observations to policies. Our networks used gated recurrent units (GRUs), which employ a gating mechanism to selectively process past information when incorporating inputs^32^, though other recurrent architectures could also have been used (see Discussion).

### Predicting choices with tiny RNNs

To analyze the experimental data, we optimized the parameters in all RNNs and cognitive models to predict the animal’s recorded choices with maximum likelihood (equivalent to minimizing cross-entropy). Given the substantial difference in the number of free parameters between RNNs (e.g, 40-80 for 1-2 unit RNNs) and cognitive models (e.g, 2-10 parameters; Fig. S1), we compared models via nested cross-validation, a procedure that uses different trials to train, validate, and evaluate the models (Fig. 1c; see Methods for why AIC or BIC are not appropriate in this setting). We then averaged each model’s predictive performance across subjects in each dataset. Our primary focus was comparing models with an equal number of dynamical variables (*d*), as these use the same number of scalar variables to summarize past experiences and specify policies. Note that dynamical variables are distinct from the model parameters: dynamical variables evolve over time, representing the agent’s current beliefs about the task and dictating their actions; model parameters, in turn, are stable and specify the rules by which the dynamical variables evolve over time.

We found that very small RNNs predicted animals’ choices more accurately than classical cognitive models and all of their variants across all tested datasets (Fig. 1e for three datasets and Fig. S2 for two additional datasets, evidenced by the fact that the lowest (best) scores in each plot are achieved by an RNN; also see Fig. S10a-c for test accuracies). In particular, RNNs outperformed all ideal Bayesian observer models, which perform exact inference based on knowledge of the task structure, suggesting that animal behavior in these tasks is not optimal. At the group level, the highest predictive performance was achieved by very small RNNs — two-unit RNNs in the reversal learning and two-stage tasks, and four-unit RNNs in the transition-reversal two-stage task. At the individual subject level, the highest predictive performance was also achieved by similarly small RNNs (Fig. S3). Crucially, each RNN with *d* units outperformed all classical cognitive models with *d* dynamical variables (Fig. 1e; evidenced by the absence of data points dots below the blue line). The fitted RNNs also reproduced key behavioral metrics commonly used to analyze each task, including choice probabilities around reversals in the reversal learning task^11^, and stay probabilities in two-stage and transition-reversal two-stage tasks^12,13^ (Fig. 1f). Finally, these fitted RNNs produced highly robust and consistent predictions across model instances, suggesting that the strategies discovered by our approach are robust to variations intrinsic to empirical data (Fig. S12). These results demonstrate that tiny RNNs are versatile models of behavior, reproducing well-known behavioral patterns in reward learning tasks and capturing more variance in animal behavior than classical cognitive models.

Adding more dynamical variables to a tiny RNN did not always improve predictive performance, suggesting that animal behavior in these tasks is low-dimensional (Fig. 1e; notice the blue line flattening or curving upwards as *d* increases). The dimensionality of a given behavior (*d*_∗_) is defined as the minimal number of functions of the past required to optimize the predictability of future behavior^33,34^ (also see Supplementary Discussion 2.3). In the reversal learning and two-stage tasks, RNN predictions were optimized with just 1-2 units (Fig. S3a-d), suggesting that most individual animal behaviors in these tasks have dimensionality *d*_∗_ = 1 or *d*_∗_ = 2. In the transition-reversal two-stage task, RNN predictions were optimized with 1-4 units (Fig. S3e), suggesting that individual animal behavior in this task has dimensionality ranging from *d*_∗_ = 1 to *d*_∗_ = 4. In some cases, predictive performance even decayed when adding dynamical variables, indicating the lack of data for fitting more flexible models. The low *d*_∗_ values found for these tasks are unsurprising given the requirement for binary choices and the presence of a single choice state. While tasks with this level of simplicity are extremely common in neuroscience and psychology research due to a focus on experimental control and interpretability, one may wonder if the performance of tiny RNNs is limited to such low-dimensional tasks. To address this question, we will demonstrate in the next sections that tiny RNNs are also effective in more complex, higher-dimensional scenarios, highlighting their versatility across various experimental paradigms.

### Flexibility and data requirements of tiny RNNs

The superior performance of tiny RNNs compared to classical cognitive models stems from the increased flexibility afforded by a much larger (4-40 times) number of free parameters. This flexibility allows RNNs to be molded into a wider range of behaviors than possible with classical cognitive models. To evaluate the limits of this flexibility, we simulated the behavior of RL and Bayesian inference agents in the reversal learning and two-stage tasks and fitted both RNN and cognitive models to these synthetic data (see Supplementary Results 1.1, Fig. S4, S5, S6, S7). We found that the tiny RNNs achieved predictive performance similar to the ground-truth model that generated the behavior, with the best-performing RNN having the same dimensionality as the ground-truth model. These results suggest that RNN models can serve as a superset of classical cognitive models despite using a single architecture consistently across tasks and datasets and requiring only minimal manual engineering. Incidentally, these results also demonstrate that these cognitive strategies are identifiable and robustly recoverable (see Supplementary Results 1.1 and Fig. S11, S13), and that our training procedure successfully prevented overfitting, which can be diagnosed when the fitted model achieves higher performance on the training dataset and lower on the test dataset relative to the data-generating model.

An important caveat of the flexibility of RNNs is that these models often require more training data than simpler models to achieve optimal performance. This is not a concern if enough data is available to support the continual improvement of the RNN fits, which can improve beyond the point where cognitive models plateau (Fig. S8). If data is scarce, however, the training procedure may not adequately constrain the RNN parameters. Indeed, we found that the performance of RNNs decayed rapidly as fewer data were used for training, eventually dropping below the performance of simpler, data-efficient cognitive models. Specifically, we found that 500-3,000 trials were required for training and validation before RNNs could outperform cognitive models (Fig. S8). While datasets of this size are typical in animal experiments, this requirement presents a challenge for human studies, which typically rely on less data but multiple participants to achieve statistical power.

To overcome the limitation of RNNs in scenarios of limited data per participant, we developed a knowledge distillation framework whereby data from multiple participants can be used to enhance the predictive performance of individual-level RNNs^35^. This approach consists of first training a single large “teacher” model on data from all subjects and then using the teacher model to train smaller and more interpretable “student” RNNs for individual subjects (Fig. 2a). Each student RNN is trained on data from a single subject, but instead of predicting binary choices, it aims to match the probabilistic policy of the larger teacher model. Applying this method to a representative mouse in the transition-reversal two-stage task (Fig. 2b), we found that the tiny student RNNs outperformed cognitive models (the best model-free model with the same dimensionality) with as few as 350 trials per subject, compared to the 3,000 trials required without knowledge distillation (“solo RNNs”). This demonstrates that tiny RNNs, when leveraging data from multiple participants, can outperform classical cognitive models even with dataset sizes typical in human experiments.

### Predicting choices with tiny RNNs for human tasks

To expand the applicability of our approach, we next examined how tiny RNNs perform on human decision-making tasks, which typically involve fewer trials per subject and use slightly more complex designs than animal studies. We applied our method to analyze three tasks, representing a range of experimental paradigms commonly used in cognitive neuroscience research: a three-armed reversal learning task (three actions; 160 trials per subject), a four-armed drifting bandit task (four actions and continuous rewards; 150 trials per subject), and the original two-stage task (six actions and three choice states; 200 trials per subject) (Fig. 2c). Given the limited per-subject data, we used knowledge distillation and implemented an interspersed split protocol to train and evaluate the models (see Methods; also see the cross-subject split protocol in Fig. S39). As before, we compared tiny RNNs to over 10 established cognitive models for these tasks.

We found that student RNNs with 5-20 units provided the best fit to human behavior, despite the limited trials per subject. This suggests that human behavior in these tasks had higher dimensionality compared to the animal studies (Fig. 2d; also Fig. S9 for the distribution of dimensionality), likely because these tasks had more actions, and, in some cases, also more choice states and continuous rewards. Notably, however, even RNNs with just 2-4 units outperformed all cognitive models of equal dimensionality (Fig. 2d; also see Fig. S10d-f for test accuracies). These results demonstrate that tiny RNNs can efficiently capture complex behavior and outperform cognitive models in predicting single-subject behavior across a range of human decision-making tasks.

### Interpreting and comparing one-dimensional models

A major advantage of smaller RNNs is their potential to yield interpretable insights into cognitive processes, which has historically been challenging with larger neural networks due to their complexity. We propose an interpretative framework grounded in the theory of discrete dynamical systems, which describes how a system’s state (e.g., dynamical variables in an RNN or cognitive model) changes over time as a function of inputs (*r*ewards, observations, and past actions). We begin by examining models with a single dynamical variable (*d* = 1; Fig. 3a). In these models, the state at trial t is fully characterized by the policy’s logit (log-odds, defined as *L*(*t*) = log (Pr_*t*_(*A*_1_)/ Pr_*t*_(*A*_2_))), representing the agent’s current preference for one action over the other. The evolution of this state is given by the logit-change (Δ*L*(*t*) = *L*(*t* + 1) − *L*(*t*)), representing how different inputs alter the agent’s preferences between trials. Intuitively, the “logit” and “logit-change” incurred by an “input” are analogous to the “position” and “velocity” of a system subject to a “force”. By visualizing *L*(*t*) and Δ*L*(*t*) for each trial in a phase portrait, we can reveal important insights about the system’s behavior.

**Fig. 3.**
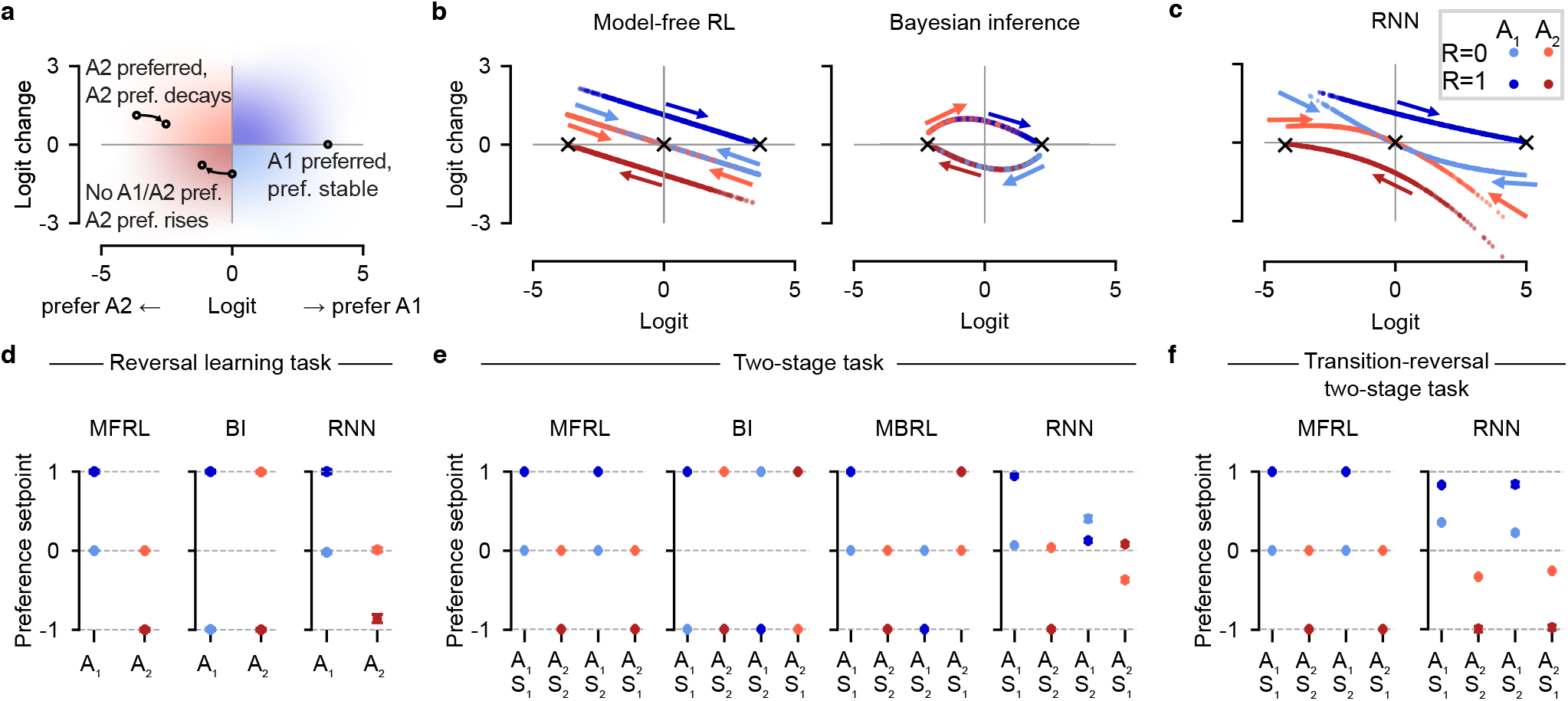
Dynamical systems analyses for interpretation and comparison of one-dimensional models. **(a)** Schematic showing how the agent’s preference evolves over consecutive trials. When action A2 is favored (logit *L*(*t*) *<* 0), a positive logit change (Δ*L*(*t*) *>* 0) results in a reduced preference for A2. When neither action is favored (indifference; logit *L*(*t*) = 0), a negative logit change (Δ*L*(*t*) *<* 0) results in a preference for A2. A preference level associated with a zero logit change (Δ*L*(*t*) = 0) is a stable fixed point. Colors indicate the preferred action (blue: *A*_1_; red: *A*_2_) and preference change (dark: increase; light: decrease). **(b-c)** Phase portraits illustrating how action preferences change as a function of current preferences (logit), action taken (*A*_1_, blue; *A*_2_, red), and reward received (*R* = 0, light; *R* = 1, dark). Points represent trials and are colored according to the trial input; colored arrows indicate the flow direction of the model state (logit) after receiving the corresponding input. **(b)** Two one-dimensional cognitive models fitted to the choices of one monkey in the reversal learning task. **(c)** One-unit GRU fitted to the same monkey data. **(d-f)** Preference setpoints (*u*_*I*_): Long-term action preferences after repeated exposure to input *I* (or, analogously, the instantaneous effect of *I* on normalized preferences). Colors indicate input types; error bars show standard deviations across model instantiations in nested cross-validation. **(d)** Reversal learning task; monkey data. **(e)** Two-stage task; rat data. **(f)** Transition-reversal two-stage task; mouse data.

To illustrate this interpretative framework, we first compared the phase portraits of two widely used cognitive models: a one-dimensional model-free RL and a one-dimensional Bayesian inference model, each fitted to the choices of one monkey in the reversal learning task (Fig. 3b). In model-free RL, a reward for action *A*_1_ (dark blue) is associated with a positive logit-change, while a reward for *A*_2_ (dark red) is associated with a negative logit-change, reflecting increases and decreases in preference for *A*_1_ over *A*_2_, respectively. Unrewarded actions (light blue and light red) generally lead to smaller preference changes. In contrast, the Bayesian inference model treats an unrewarded action as equivalent to a reward for the other action. We can also compare the fixed points of the models 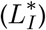, representing states in which preferences are unaffected by a given input (Δ*L*_*I*_ = 0). Model-free RL exhibits three types of stable fixed points (attractors), corresponding to high preference for *A*_1_, high preference for *A*_2_, and indifference. Bayesian inference has only two types of attractors corresponding to high preferences for either action, as the reward-action symmetry prevents convergence to an indifference state. The shape of the portraits (straight or curved) illustrates how each model processes unexpected rewards. When a disfavored action is rewarded (dark blue dots with extreme negative logit values), a model-free RL agent will experience a large state change (because of a large prediction error), while a Bayesian inference agent will experience only a small state change (because of a strong prior at extreme logit values).

Having shown how phase portraits enable insights into models for which the ground truth is known, we next applied this approach to analyze tiny RNNs, aiming to uncover cognitive processes underlying animal behavior. Since RNNs can mimic the behavior of both model-free RL and Bayesian inference agents (Supplementary Results 1.1, Fig. S4, S5, S6, S7), their phase portrait can reveal whether signatures of either cognitive model are present. Applying this rationale to a one-unit RNN fitted to the same data as above, we found that its phase portrait showed multiple model-free RL characteristics (Fig. 3c, see S14g for results with another monkey). For example, unrewarded trials moved the system towards indifference (note attractor at L = 0), unexpected rewards caused large preference changes (large positive logit-changes for dark blue dots in the left region), and unrewarded actions were not treated as rewards for the unchosen action (non-overlapping light red and dark blue curves). This suggests that the monkey’s behavior aligns more closely with a model-free RL agent than a Bayesian inference model. While these conclusions could have been achieved with conventional model comparisons, the phase portraits offer a more nuanced understanding of how specific aspects of the monkey’s behavior relate to each model.

Besides revealing signatures of known models in the RNN dynamics, phase portraits also support the discovery of entirely novel signatures. For example, curved lines suggest a non-constant, state-dependent learning rate (Fig. 3c). The decoupling of the two *R* = 0 (light blue and light red) curves, a sign of choice perseveration, is only present for extreme values of logit, suggesting a peculiar pattern of preference-dependent choice perseveration. The asymmetry between the two non-zero fixed points suggests greater sensitivity to rewards from action *A*_1_ than to rewards from *A*_2_, akin to a reward-dependent choice bias (Fig. 3c). These three signatures — “state-dependent learning rate”, “preference-dependent perseveration”, and “reward-dependent bias” — are absent from all cognitive model variants considered and from any published analysis of these tasks in the literature. Some of these signatures were found across animals, while others are individual-specific (e.g., Fig. S16a), highlighting the importance of modeling individual subjects. Crucially, we validated each of these insights using targeted hypothesis testing (see Supplementary Results 1.2, Fig. S14) or using a novel behavior-feature identifier approach (see Methods and Fig. S27).

Although phase portraits provide a comprehensive characterization of a system’s dynamics, they can be challenging to interpret, especially for tasks with a large (or infinite) number of inputs. In such cases, the essential information from phase portraits can be simplified and summarized by the model’s preference setpoints, which summarize the effect of each input on the model’s dynamics. A preference setpoint *u*_*I*_ for input *I* represents an agent’s normalized, asymptotic preference for one action over another after repeated exposure to input *I* (i.e., 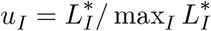). Beyond representing long-term behavior, preference setpoints indicate the instantaneous direction of change in the system’s state when presented with input I. Essentially, *u*_*I*_ summarizes the effect of input *I* in the model: |*u*_*I*_ | = 1 indicates convergence to a state of maximum preference, and *u*_*I*_ = 0 indicates convergence to a state of indifference. We computed *u*_*I*_ for all fitted models for the reversal learning task, the two-stage task, and the transition-reversal two-stage task (Fig. 3d-f). In the two-stage task, the RNN’s preference setpoints revealed a “reward-induced indifference” phenomenon where rewards following rare transitions led to indifference (|*u*_*I*_ | ≈ 0) rather than an increased preference for a specific action (Fig. 3e; dark blue marker for *A*_1_, *S*_2_ and the dark red marker for *A*_2_, *S*_1_). A similar but weaker effect (|*u*_*I*_ | < 1) was found in the transition-reversal two-stage task (Fig. 3f). This “reward-induced indifference” was absent from all cognitive model variants considered and from the literature more broadly, despite being found in several rats and mice (Fig. S15). Importantly, the strength of this effect correlated with better task performance across animals (*ρ* = 0.62, *p* = 0.008, Fig. S17), demonstrating that behavioral patterns discovered by our approach can have meaningful behavioral relevance.

### Interpreting and comparing multi-dimensional models

We next extended our interpretive framework to models with more than one dynamical variable (*d* > 1). While these models can still be analyzed with phase portraits, the policy logit can no longer fully characterize their state. For models with *d* = 2 dynamical variables, we can instead compute the 2D vector field, where axes represent both dynamical variables and arrows indicate how the state changes in each trial. To illustrate this approach, we analyzed vector fields of a two-dimensional model-free RL model and a two-unit RNN with diagonal readout (see Methods) fitted to the choices of one monkey in the reversal learning task (Fig. 4a-b; Fig.S20). We found that arrows in the model-free RL vector field were axis-aligned, indicating that only the value of the chosen action changed on each trial; in contrast, arrows in the RNN’s vector field were slightly tilted in rewarded trials, indicating a decay (forgetting) in the value of the unchosen action (Fig. 4a). Additionally, arrows in both models converged to a line in the space, indicating line attractor dynamics (white crosses in Fig. 4a). Interestingly, in the RNN’s vector field, arrows in unrewarded trials converged to the diagonal (*h*_1_ = *h*_2_) line, suggesting that the value of an unrewarded action drifted towards the value of the alternative action and not towards zero as expected. As a validation of this peculiar “drift-to-the-other” prediction, we found that a model-free RL model augmented with drift-to-the-other outperformed a model-free RL model augmented with conventional forgetting (drift-to-zero) and performed nearly as well as the two-unit RNN (Fig.S21).

**Fig. 4.**
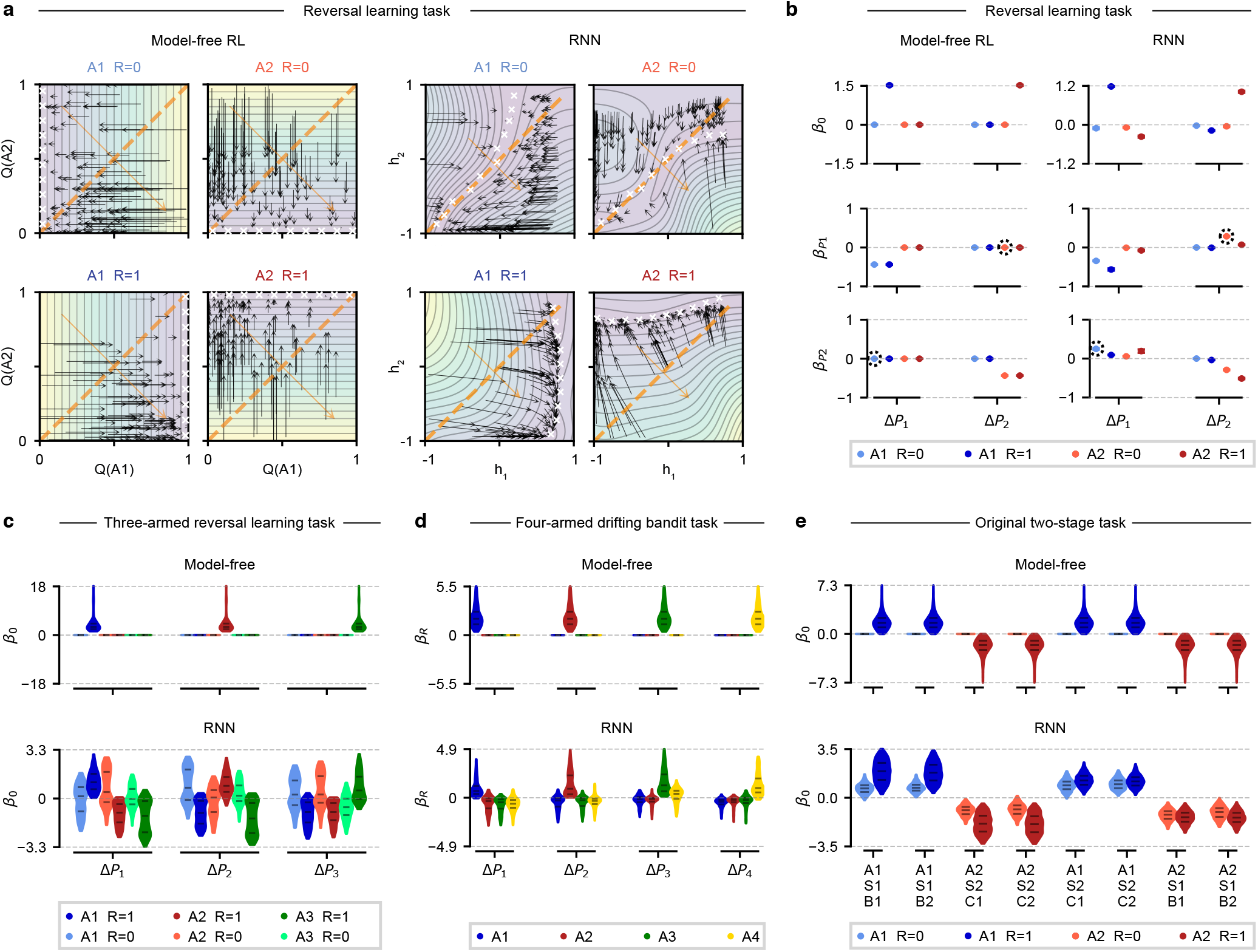
Dynamical systems analyses for interpretation and comparison of multi-dimensional models. **(a)** Vector field analysis of two-dimensional models fitted to the choices of one monkey in the reversal learning task. Each panel illustrates the effect of one input on the state variables (axes). Black arrows: flow lines indicating state changes per trial; white crosses: attractor states; dashed lines: indifference states; orange arrows: readout vectors; background-color: dynamics speed (purple: slow; green: medium; yellow: fast). *Left:* Model-free RL model; axis-aligned arrows indicate that only the value of the chosen action is updated in each trial, with values converging to the reward magnitude. *Right:* Two-unit RNN. *Top:* in unrewarded trials (*R* = 0), convergence of arrows to diagonal line suggest a drift-to-the-other pattern. **(b)** Dynamical regression analysis for 2D model-free RL (left) and two-unit RNN (right). *P*_*i*_, Δ*P*_*i*_: preference and preference change for action *A*_*i*_. Regression coefficients describe how each action preference changes in each trial as a function of its own current value 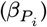, of the current preference for the other action 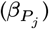, and independently of one’s current preferences (baseline_*i j*_ *β*_0_). Points represent mean coefficients across outer rounds and error bars represent standard deviations across outer rounds. Circled coefficients in the RNN model indicate a drift-to-the-other pattern. **(c-e)** Dynamical regression analysis for a model-free RL model and an RNN with equal dimensionality *d >* 2, fitted to human behavior. Violin plots show distributions over subject-level coefficients. **(c)** Three-dimensional models fitted to data from three-armed reversal learning task. *P*_*i*_, Δ*P*_*i*_: preference and preference change for action *A*_*i*_. *β*_0_ is the constant term in the linear regression. **(d)** Four-dimensional models fitted to data from four-armed drifting bandit task. *P*_*i*_, Δ*P*_*i*_: preference and preference change for action *A*_*i*_. *β*_*R*_ is the coefficient for the continuously-valued reward in the linear regression. **(e)** Three-dimensional models fitted to data from the original two-stage task. *L*_*i*_, Δ*L*_*i*_: logit and logit change for each choice state (*L*_1_: first-stage, *L*_2_, *L*_3_: second-stage states).

To interpret models with more than two dynamical variables (*d* > 2), we introduce an alternative method based on “dynamical regression”. This method approximates the one-step dynamics of the model’s internal state as a linear function of its current state. Applied to the two-unit RNN studied above, the dynamical regression coefficients indicate that the value of an unrewarded chosen action is positively influenced by the value of the alternative action — a re-discovery of the drift-to-the-other pattern (Fig. 4b). We then used the same approach to analyze human behavior in the three human tasks introduced previously (Fig. 2c), performing separate regressions for each subject-specific RNN trained with knowledge distillation. From the distributions of dynamical regression coefficients, we obtained several novel insights into human behavior in these tasks: (1) in the three-armed reversal learning task, trial outcomes affect preferences for all actions, but differently for chosen versus unchosen actions (Fig. 4c); (2) in the four-armed drifting bandit task, rewards can sometimes reduce the value of unchosen actions (Fig. 4d); (3) in the original two-stage task, the value of a rewarded action drifts towards a nonzero setpoint that depends on the transition type (Fig. 4e). None of these patterns are captured by classical cognitive models, and augmenting these models with the corresponding mechanisms derived from RNN analyses leads to improved performance (see Supplementary Results 1.4, 1.5, 1.6 and Fig.S28, S29, S30, S31, S32, S33, S34, S35, S36)

### Interpreting behavior in task-optimized neural networks

Our interpretative framework can be used not only to analyze models fit to behavior, but also to analyze the behavior of larger neural networks trained to achieve optimal task performance in decision-making tasks, offering a new way to compare behavior in biological and artificial systems. To illustrate this, we trained an RNN agent within a meta-reinforcement learning (meta-RL) framework to maximize total rewards in a two-stage task (Fig.5a; Fig.S25). While previous work suggested that such agents use a model-based RL strategy based on patterns “stay probability” and “prediction errors”^24^, these patterns could also have been generated by alternative strategies^36^.

To gain mechanistic insights into the behavior of meta-RL agents, we compared their phase portraits with those of different cognitive models. We found that the dynamics of the trained meta-RL agent (Fig. 5b) resembled a one-dimensional Bayesian inference agent (Fig. 5c), and not a model-based RL strategy as previously suggested (Fig. 5d). The dynamics of the trained meta-RL agent also did not resemble the dynamics we inferred in animals performing the same task (Fig. S15). Interestingly, before converging to a Bayesian inference strategy, the agent’s representations differed substantially from any known cognitive models (Fig. S26b). Even after convergence, we identified subtle deviations from the Bayesian inference model (Fig. 5b-c). In particular, each logit value in the meta-RL agent was associated with a range of logit-change values (Fig. 5b, note parallel lines), in contrast to the single logit-change value observed in the Bayesian inference agent. Our analyses determined that these adjacent curves reflect a “history effect” whereby exact Bayesian inference is distorted by the representation of historical input sequences (see Fig. S26c). This illustrates how our approach contributes to understanding computational processes in both biological and artificial systems.

**Fig. 5.**
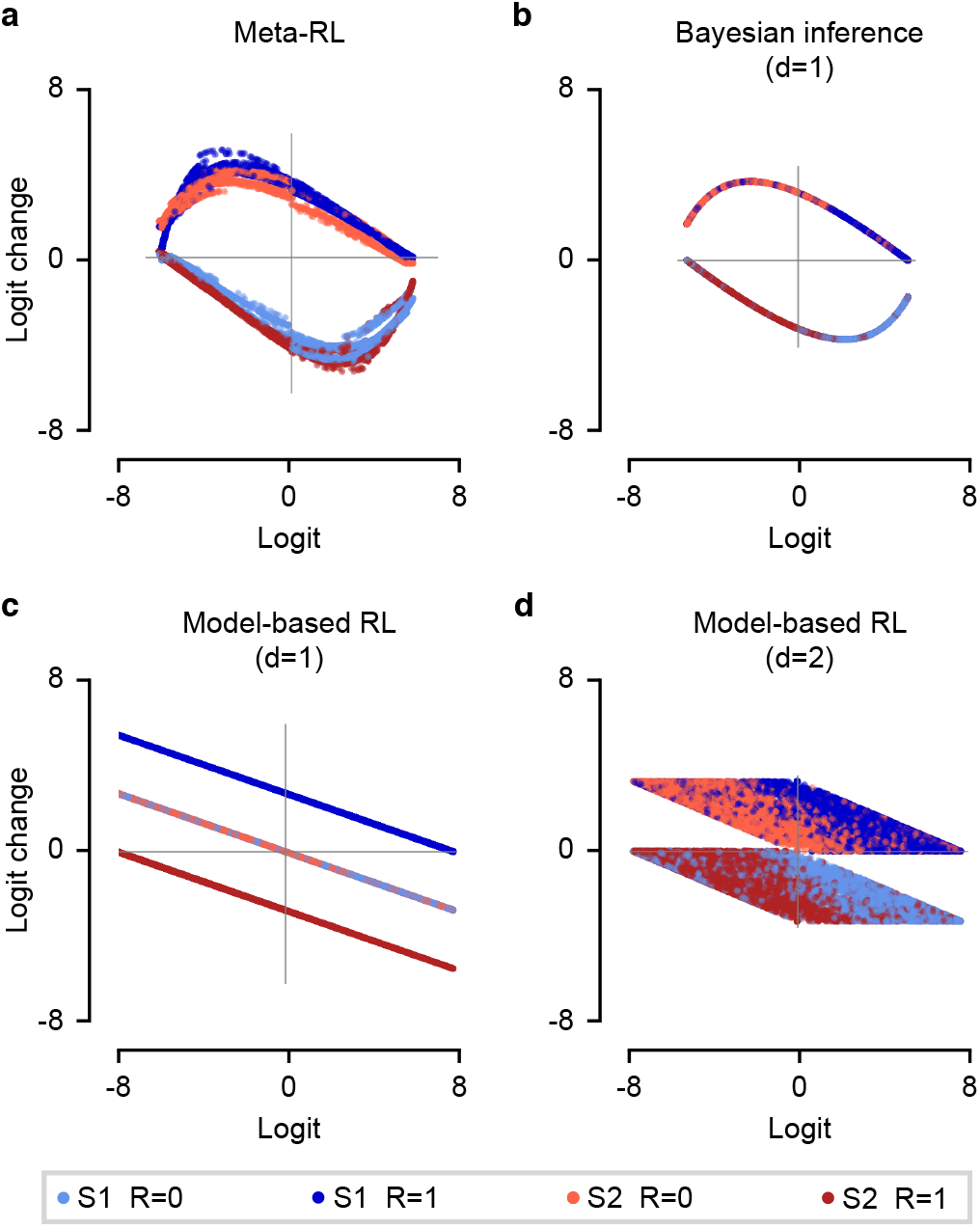
Logit analysis of meta-RL agent and cognitive models in the two-stage task. **(a)** Meta-RL agent. **(b)** One-dimensional Bayesian inference model fitted to meta-RL agent behavior (includes task-optimal Bayesian model). **(c)** One-dimensional model-based RL model fitted to meta-RL agent behavior. **(d)** Two-dimensional model-based RL model^24^ fitted to meta-RL agent behavior. Logit-change patterns suggest similarities between meta-RL agent and Bayesian inference model, and differences relative to model-based RL models and observed animal strategies (cf. Fig. S15).

## Discussion

We introduce a new method for modeling decision-making behavior using tiny recurrent neural networks (RNNs). We demonstrate its effectiveness across six reward learning tasks and eight datasets where tiny RNNs outperformed classical cognitive models in predicting individual subject behavior. Our approach leverages a dynamical systems framework to interpret the trained networks, revealing novel cognitive strategies without extensive model comparisons. These include state-dependent learning rates, a “reward-induced indifference” effect, and new forms of value updating, choice perseveration, and choice biases. Our approach also estimates the dimensionality of behavior and analyzes decision-making mechanisms in task-optimized neural networks. Overall, tiny RNNs offer a powerful tool for understanding decision-making across species, combining the flexibility of neural networks with the interpretability of cognitive models.

Despite their small size, tiny RNNs are remarkably flexible and can capture both normative and suboptimal behaviors. This flexibility stems from the increased number of free parameters, allowing tiny RNNs to model a wider range of behaviors than classical cognitive models. In particular, tiny RNNs can mimic the choices of normative agents based on reinforcement learning and Bayesian inference. Leveraging this flexibility, however, requires sufficiently large datasets. We found that tiny RNNs need up to a few thousand trials per subject to outperform cognitive models. While common in animal studies, this is rare in human studies. To mitigate this issue, we proposed a knowledge-distillation approach, which leverages data from multiple subjects to achieve excellent performance with only a few hundred trials per subject. This significantly expands the potential applications of our approach, particularly for human studies in cognitive neuroscience and computational psychiatry.

We developed a unified approach for interpreting and comparing cognitive models and RNNs, revealing novel insights into the underlying cognitive processes. This approach views a decision-making agent as a dynamical system whose internal state changes over time as a function of rewards, observations, and past actions. Given the low dimensionality of our fitted models, we could leverage graphical and analytical methods from dynamical systems theory to visualize, compare, and interpret the dynamics of each model. Additionally, our proposed dynamical regression approach could effectively summarize the dynamics of models of with higher dimensionalities. This unified framework enables direct comparisons between the strategies learned by RNNs and those assumed by cognitive models. Importantly, we validated these strategies identified by our approach using standard model recovery and validation techniques^14^.

The number of RNN units needed to optimally predict behavior estimates that behavior’s dimensionality^33,34^. In our study, tiny RNNs performed as well as or better than larger RNNs, suggesting that the studied behaviors have low dimensionality. This finding is consistent with other studies showing low-dimensional dynamics in the brain^23^ and the effectiveness of low-rank RNNs in modeling complex neural dynamics^37^. Our study also highlights the important distinction between the dimensionality of dynamics (number of dynamical variables) and their complexity (e.g., nonlinearity), mirroring a similar separation in low-rank RNNs^37–40^. We note that the identification of dynamical variables from experimental data is an active research area across neuroscience, complex systems, and physics, with efforts to extract key variables from neural data, physical system recordings, and multiscale complex systems^37,41–43^.

In developing our approach, we made several key technical decisions with specific advantages and limitations. First, we chose to use gated recurrent units (GRUs) due to their Markovian property and ability to process information selectively^32^. However, the GRU updating equation may limit the complexity of dynamics captured by tiny RNNs (see theoretical analysis in^44^ and Fig. S37 for the case of one-dimensional discrete dynamics). While sufficient for the tasks studied here, more complex behaviors may require different architectures. Additionally, our choices in model training and regularization balanced flexibility and generalization. While effective across multiple tasks, our approach may face challenges in scaling to more complex tasks. Future work should thus explore alternative architectures, training methods, or interpretability techniques to address these limitations.

Our findings have broad implications for understanding cognitive and neural mechanisms, with potential applications in computational psychiatry and other fields. The ability of tiny RNNs to accurately model individual-subject behavior makes them promising for studying individual differences in decision-making (e.g., Fig. S14, S3, S9, S15, S16), a key aspect of computational psychiatry. In the future, we aim to extend our framework to more complex, naturalistic settings and other cognitive domains, such as perceptual decision-making and memory. Finally, the potential of our approach to link neural activity with behavior could lead to more integrated models of cognition, bridging computational and neurobiological levels of analysis. In conclusion, our work with tiny RNNs opens new avenues for research in cognitive science, neuroscience, and AI, offering a powerful tool for uncovering the computational principles underlying adaptive behavior.

## Acknowledgments

We thank the support from the Swarma Club and Causal Emergence Reading Group supported by the Save 2050 Programme jointly sponsored by the Swarma Club and X-Order. We thank K. T. Jensen, H. Xiong, Z. Jin, and X. Li for their inspirational discussions. This work was supported by the Kavli Institute for Brain and Mind (KIBM) Innovative Research Grant #2022-2209 and in part by National Science Foundation (NSF) awards CNS-1730158, ACI-1540112, ACI-1541349, OAC-1826967, OAC-2112167, CNS-2100237, CNS-2120019, the University of California Office of the President, and the University of California San Diego’s California Institute for Telecommunications and Information Technology/Qualcomm Institute. Thanks to CENIC for the 100Gbps networks.

## Author Contributions

All authors conceived the study, participated in the discussions, wrote the paper, and acquired funding.

## Declaration of Interests

All authors declare no competing interests.

### Data availability

The monkey dataset on the reversal learning task can be found in^11,45^. The rat dataset on the two-stage task can be found in^13^. The three mice datasets on the reversal learning task, two-stage task, and transition-reversal two-stage task can be found in^12^. The human dataset on the three-armed reversal learning task can be found in^46^. The human dataset on the four-armed drifting bandit task can be found in^47^. The human dataset on the original two-stage task can be found in^48^.

## Code availability

All code used in this paper is available from the corresponding author upon request.

## Methods

### Tasks and datasets

#### Reversal learning task

The reversal learning task is a paradigm designed to assess subjects’ ability to adapt their behavior in response to changing reward contingencies. In each trial, subjects are presented with two actions, *A*_1_ and *A*_2_, yielding a unit reward with probability 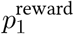 and 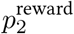, respectively. These reward probabilities remain constant for several trials before switching unpredictably and abruptly, without explicit cues. When this occurs, the action associated with the higher reward probability becomes linked to the lower reward probability, and vice versa. The task necessitates continuous exploration of which action currently has a higher reward probability in order to maximize total rewards. For consistency with the other animal tasks, we assume that actions (*A*_1_ and *A*_2_) are made at the choice state, and *A*_*i*_ deterministically leads to state *S*_*i*_, where the reward is delivered.

In the Bartolo dataset^11^, two monkeys completed a total of 15,500 trials of the reversal learning task with two state-reward types: (1) 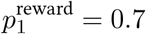 and 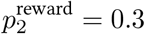 (2) 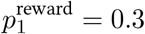 and 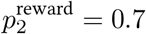 Blocks were 80 trials long, and the switch happened at a “reversal trial” between 30 and 50. We predicted the behavior from trials 10 to 70, similar to the original preprocessing procedure 11 because the monkeys were inferring the current block type (“what” block: choosing from two objects; “where” block: choosing from two locations) in the first few trials.

In the Akam dataset^12^, ten mice completed a total of 67,009 trials of the reversal learning task with three state-reward types: (1) 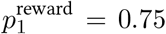 and 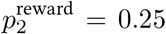 (2) 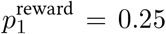 and 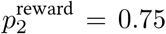 (3) 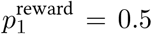 and 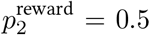 (neutral trials). Block transitions from non-neutral blocks were triggered 10 trials after an exponential moving average (tau = 8 trials) crossed a 75% correct threshold. Block transitions from neutral blocks occurred with a probability of 10% on each trial after the 15th of the block to give an average neutral block length of 25 trials.

#### Two-stage task

The two-stage task is a paradigm commonly used to distinguish between the influences of model-free and model-based reinforcement learning on animal behavior^49^ and later reduced in^36^. In each trial, subjects are presented with two actions, *A*_1_ and *A*_2_, while at the choice state. Action *A*_1_ leads with a high probability to state *S*_1_ and a low probability to state *S*_2_, while action *A*_2_ leads with a high probability to state *S*_2_ and a low probability to state *S*_1_. From second-stage states *S*_1_ and *S*_2_, the animal can execute an action for a chance of receiving a unit reward. Second-stage states are distinguishable by visual cues and have different probabilities of yielding a unit reward: 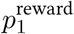 for *S*_1_ and 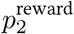 for *S*_2_ These reward probabilities remain constant for several trials before switching unpredictably and abruptly. When this occurs, the second-stage state associated with the higher reward probability becomes linked to the lower reward probability, and vice versa.

In the Miller dataset^13^, four rats completed a total of 33,957 trials of the two-stage task with two state-reward types: (1) 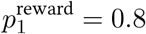 and 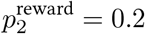 (2) 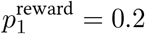 and 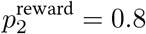. Block switches occurred with a 2% probability on each trial after a minimum block length of 10 trials.

In the Akam dataset^12^, ten mice completed a total of 133,974 trials of the two-stage task with three state-reward types: (1) 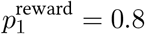 8 and 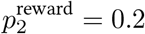 (2) 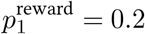 and 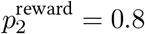 (3) 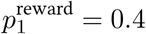 and 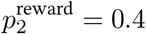 (neutral trials). Block transitions occur 20 trials after an exponential moving average (tau = 8 trials) of the subject’s choices crossed a 75% correct threshold. In neutral blocks, block transitions occurred with 10% probability on each trial after the 40th. Transitions from non-neutral blocks occurred with equal probability either to another non-neutral block or to the neutral block. Transitions from neutral blocks occurred with equal probability to one of the non-neutral blocks.

#### Transition-reversal two-stage task

The transition-reversal two-stage task is a modified version of the original two-stage task, with the introduction of occasional reversals in action-state transition probabilities^12^. This modification was proposed to facilitate the dissociation of state prediction and reward prediction in neural activity and to prevent habit-like strategies that may produce model-based control-like behavior without forward planning. In each trial, subjects are presented with two actions, *A*_1_ and *A*_2_, at the choice state. One action commonly leads to state *S*_1_ and rarely to state *S*_2_, while the other action commonly leads to state *S*_2_ and rarely to state *S*_1_. These action-state transition probabilities remain constant for several trials before switching unpredictably and abruptly, without explicit cues. In the second-stage states *S*_1_ and *S*_2_, subjects execute an action for a chance of receiving a unit reward. The second-stage states are visually distinguishable and have different reward probabilities that also switch unpredictably and abruptly, without explicit cues, similar to the other two tasks.

In the Akam dataset^12^, seventeen mice completed a total of 230,237 trials of the transition-reversal two-stage task with two action-state types: (1) Pr(*S*_1_|*A*_1_) = Pr(*S*_2_|*A*_2_) = 0.8 and Pr(*S*_2_|*A*_1_) = Pr(*S*_1_|*A*_2_) = 0.2; (2)Pr(*S*_1_|*A*_1_) = Pr(*S*_2_|*A*_2_) = 0.2 and Pr(*S*_2_|*A*_1_) = Pr(*S*_1_|*A*_2_) = 0.8. There were also three state-reward types: (1) 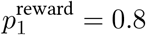 and 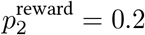 (2) 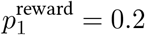 and 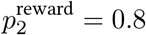 (3) 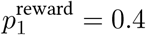 and 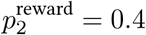 (neutral trials). Block transitions occur 20 trials after an exponential moving average (tau = 8 trials) of the subject’s choices crossed a 75% correct threshold. In neutral blocks, block transitions occurred with 10% probability on each trial after the 40th. Transitions from non-neutral blocks occurred with equal probability (25%) either to another non-neutral block via reversal in the reward or transition probabilities, or to one of the two neutral blocks. Transitions from neutral blocks occurred via a change in the reward probabilities only to one of the non-neutral blocks with the same transition probabilities.

#### Three-armed reversal learning task

In the Suthaharan dataset^46^, 1010 participants (605 participants from the pandemic group and 405 participants from the replication group) completed a three-armed probabilistic reversal learning task. This task was framed as either a non-social (card deck) or social (partner) domain, each lasting 160 trials divided evenly into 4 blocks. Participants were presented with 3 actions (*A*_1_, *A*_2_, *A*_3_; 3 decks of cards in the non-social domain frame or 3 avatar partners in the social domain frame), each containing different amounts of winning (+100) and losing (−50) points. The objective was to find the best option and earn as many points as possible, knowing that the best option could change.

The task contingencies started with 90%, 50%, and 10% reward probabilities, with the best deck/partner switching after 9 out of 10 consecutive rewards. Unbeknownst to the participants, the underlying contingencies transitioned to 80%, 40%, and 20% reward probabilities at the end of the second block, making it more challenging to distinguish between probabilistic noise and genuine changes in the best option.

#### Four-armed drifting bandit task

The Bahrami dataset^47^ includes 975 participants who completed the 4-arm bandit task^50^. Participants were asked to choose between four options on 150 trials. On each trial, they chose an option and were given a reward. The rewards for each option drifted over time in a manner known as a restless bandit, forcing the participants to constantly explore the different options to obtain the maximum reward. The rewards followed one of three predefined drift schedules^47^.

During preprocessing, we removed 57 participants (5.9%) who missed more than 10% of trials. For model fitting, missing trials from other subjects are excluded from the loss calculation.

#### Original two-stage task

In the Gillan dataset^48^, the original version of the two-stage task^49^ was used to assess goal-directed (model-based) and habitual (model-free) learning in individuals with diverse psychiatric symptoms. 1961 subjects (548 from the first experiment and 1413 from the second experiment) completed the task. In each trial, subjects were presented with a choice between two options (*A*_1_ or *A*_2_). Each option commonly (70%) led to a particular second-stage state (*A*_1_ → *S*_1_ or *A*_2_ → *S*_2_). However, on 30% of “rare” trials, choices led to the alternative second-stage state (*A*_1_ → *S*_2_ or *A*_2_ → *S*_1_). In the second-stage states, subjects chose between two options (*B*_1_/*B*_2_ in *S*_1_ or *C*_1_/*C*_2_ in *S*_2_), each associated with a distinct probability of being rewarded. The reward probabilities associated with each second-stage option drifted slowly and independently over time, remaining within the range of 0.25 to 0.75. To maximize rewards, subjects had to track which second-stage options were currently best as they changed over time.

For model fitting, missing stages/trials from some subjects are excluded from the loss calculation.

### Recurrent neural networks

#### Network architectures

We investigated several architectures, as described below. Our primary goal is to capture the maximum possible behavioral variance with *d* dynamical variables. While we generally prefer more flexible models due to their reduced bias, such models typically require more data for training, and insufficient data can result in underfitting and poorer performance in comparison to less flexible (simpler) models. Therefore, we aimed to balance data efficiency and model capacity through cross-validation.

After finding the best-performing model class, we performed an investigation of the network properties that contributed the most to the successfully explained variance. Analogous to ablation studies, our approach consisted of gradually removing components or adding constraints to the architectures, such as eliminating nonlinearity or introducing symmetric weight constraints. The unaffected predictive performance suggests that the examined components are not essential for the successfully explained variance. If affected, this indicates that these components can contribute to explaining additional behavioral patterns. Following this approach, we can establish connections between architectural components and their corresponding underlying behavioral patterns. The primary objective of this approach is to capture maximum variance with minimal components in the models, resulting in highly interpretable models.

##### Recurrent layer

The neural network models in this paper used the vanilla gated recurrent units (GRU) in their hidden layers^32^. The hidden state *h*_*t*_ at the beginning of trial *t* consists of *d* elements (dynamical variables). The initial hidden state *h*_1_ is set to 0 and *h*_*t*_ (*t* > 1) is updated as follows:

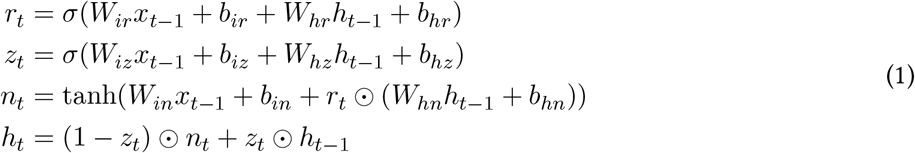

where σ is the sigmoid function, ⊙ is the Hadamard (element-wise) product, *x*_*t*−1_ and *h*_*t*−1_ are the input and hidden state from the last trial *t* − 1, and *r*_*t*_, *z*_*t*_, *n*_*t*_ are the reset, update, new gates (intermediate variables) at trial *t*, respectively. The weight matrices *W*.. and biases *b*.. are trainable parameters. The d-dimensional hidden state of the network, *h*_*t*_, represents a summary of past inputs and is the only information used to generate outputs.

Importantly, the use of GRUs means that the set of d unit activations fully specifies the network’s internal state, rendering the system Markovian (i.e., *h*_*t*_ is fully determined by *h*_*t*−1_ and *x*_*t*−1_). This is in contrast to alternative RNN architectures such as the long short-term memory^51^, where the use of a cell state renders the system non-Markovian (i.e., the output state ht cannot be fully determined by *h*_*t*−1_ and *x*_*t*−1_).

To accommodate discrete inputs, we also introduce a modified architecture called switching GRU, where recurrent weights and biases are input-dependent, similar to discrete-latent-variable-dependent switching linear dynamical systems^52^. In this architecture, the hidden state *h*_*t*_ (*t* > 1) is updated as follows:

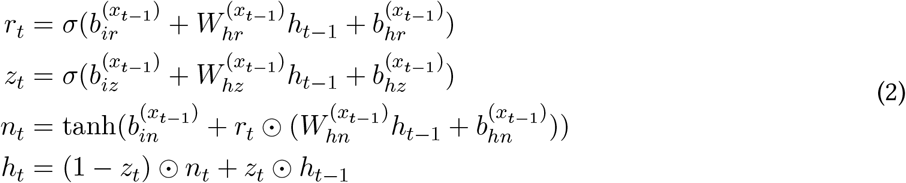

where 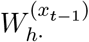 and 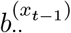 are the weight matrices and biases selected by the input *x*_*t*-1_ (i.e., each input *x*_*t*-1_ induces an independent set of weights *W*_*h*_. and biases *b*..).

For discrete inputs, switching GRUs are a generalization of vanilla GRUs (i.e., a vanilla GRU can be viewed as a switching GRU whose recurrent weights do not vary with the input). Generalizations of switching GRUs from discrete to continuous inputs are closely related to multiplicative integration GRUs^53^.

For animal datasets, we found that the switching GRU models performed similarly to the vanilla GRU models for *d* ≥ 2, but consistently outperformed the vanilla GRU models for *d* = 1. Therefore, for the results of animal datasets in the main text, we reported the performance of the switching GRU models for *d* = 1 and the performance of the vanilla GRU models for *d* ≥ 2. Mathematically, these vanilla GRU models can be directly transformed into corresponding switching GRU models:

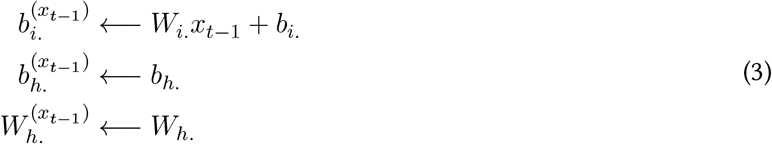

We also proposed the switching linear neural networks (SLIN), where the hidden state *h*_*t*_ (*t* > 1) is updated as follows:

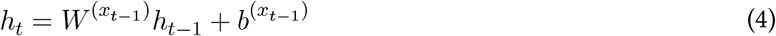

where 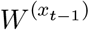 and 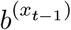 are the weight matrices and biases selected by the input *x*_*t*−1_. In some variants, we constrained 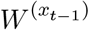 to be symmetric.

##### Input layer

The network’s input xt consists of the previous action *a*_*t*−1_, the previous second-stage state *s*_*t*−1_, and the previous reward *r*_*t*−1_ (but at = st in the reversal learning task). In the vanilla GRU networks, the input *x*_*t*_ is three-dimensional and projects with linear weights to the recurrent layer. In the switching GRU networks, the input *x*_*t*_ is used as a selector variable where the network’s recurrent weights and biases depend on the network’s inputs. Thus, switching GRUs trained on the reversal learning task have four sets of recurrent weights and biases corresponding to all combinations of *a*_*t*−1_ and *r*_*t*−1_, and switching GRUs trained on the two-stage and transition-reversal two-stage tasks have eight sets of recurrent weights and biases corresponding to all combinations of *a*_*t*−1_, *s*_*t*−1_, and *r*_*t*−1_.

##### Output layer

The network’s output consists of two units whose activities are linear functions of the hidden state *h*_*t*_. A softmax function (a generalization of the logistic function) is used to convert these activities into a probability distribution (a policy). In the first trial, the network’s output is read out from the initial hidden state *h*_1_, which has not yet been updated on the basis of any input. For d-unit networks, the network’s output scores were computed either from a fully-connected readout layer (i.e., 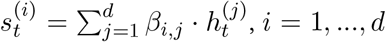) or from a diagonal readout layer (i.e., 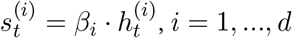). The output scores are sent to the softmax layer to produce action probabilities.

#### Network training

Networks were trained using the Adam optimizer (learning rate of 0.005) on batched training data with crossentropy loss, recurrent weight L1-regularization loss (coefficient drawn between 1e-5 and 1e-1, depending on experiments), and early-stop (if the validation loss does not improve for 200 iteration steps). All networks were implemented with PyTorch.

### Classical cognitive models

#### Models for the reversal learning task

In this task, we implemented one model from the Bayesian inference family and eight models from the model-free family (adopted from^36^ and^13^, or constructed from GRU phase portraits).

##### Bayesian inference strategy (d=1)

This model (also known as latent-state) assumes the existence of the latent-state *h*, with *h* = *i* representing a higher reward probability following action *A*_*i*_ (state *S*_*i*_). The probability Pr_*t*_(*h* = 1), as the dynamical variable, is first updated via Bayesian inference:

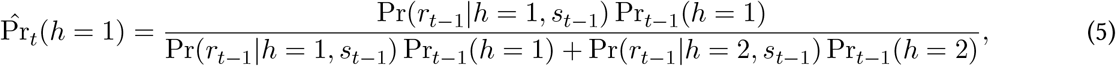

where the left-hand side is the posterior probability (we omit the conditions for simplicity). The agent also incorporates the knowledge that, in each trial, the latent-state *h* can switch (e.g., from *h* = 1 to *h* = 2) with a small probability *p*_*r*_. Thus the probability Pr_*t*_(*h*) reads,

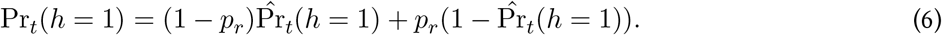

The action probability is then derived from softmax(*β*Pr_*t*_(*h* = 1), *β*Pr_*t*_(*h* = 2)) with inverse temperature *β* (*β* ≥ 0).

##### Model-free strategy (d=1)

This model hypothesizes that the two action values *Q*_*t*_(*A*_*i*_) are fully anti-correlated (*Q*_*t*_(*A*_1_) = −*Q*_*t*_(*A*_2_)) as follows:

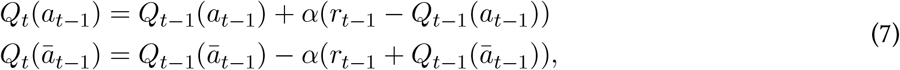

where *ā*_t−1_ is the unchosen action, and *α* is the learning rate (0 ≤ *α* ≤ 1). We specify the *Q*_*t*_(*A*_1_) as the dynamical variable.

##### Model-free strategy (d=2)

This model hypothesizes that the two action values *Q*_*t*_(*A*_*i*_), as two dynamical variables, are updated independently:

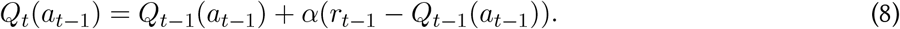

The unchosen action value *Q*_*t*_(*ā*_*t*−1_) is unaffected.

Model-free strategy with value forgetting (d=2). The chosen action value is updated as in the previous model. The unchosen action value *Q*_*t*_(*ā*_*t*−1_), instead, is gradually forgotten:

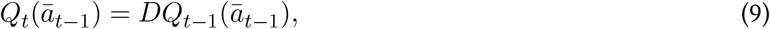

where *D* is the value forgetting rate (0 ≤ *D* ≤ 1).

##### Model-free strategy with value forgetting to mean (d=2)

This model is the “forgetful model-free strategy” proposed in^54^. The chosen action value is updated as in the previous model. The unchosen action value *Q*_*t*_(*ā*_*t*−1_), instead, is gradually forgotten to a initial value 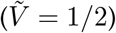:

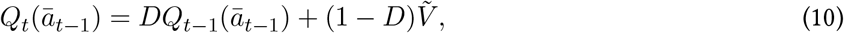

where *D* is the value forgetting rate (0 ≤ *D* ≤ 1).

##### Model-free strategy with the drift-to-the-other rule (d=2)

This strategy is constructed from the phase diagram of the two-unit GRU. When there is a reward, the chosen action value is updated as follows,

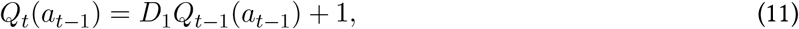

where *D*_1_ is the value drifting rate (0 ≤ *D*_1_ ≤ 1). The unchosen action value is slightly decreased:

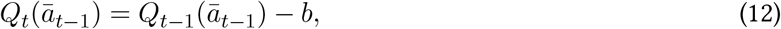

where *b* is the decaying bias (0 ≤ b ≤ 1, usually small). When there is no reward, the unchosen action value is unchanged, and the chosen action value drifts to the other:

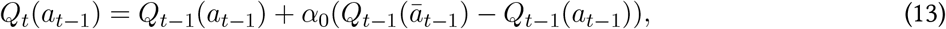

where *α*_0_ is the drifting rate (0 ≤ *α*_0_ ≤ 1).

For all model-free RL models with d = 2, the action probability is determined by softmax(*βQ*_*t*_(*A*_1_), *βQ*_*t*_(*A*_2_)).

##### Model-free strategy with inertia (d=2)

The action values are updated as the model-free strategy (d=1). The action perseveration (inertia) is updated by:

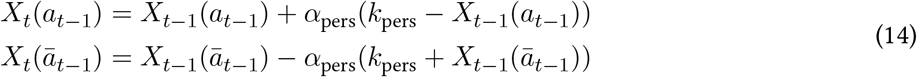

where *α*_pers_ is the perseveration learning rate (0 ≤ *α*_pers_ ≤ 1), and k_pers_ is the single-trial perseveration term, affecting the balance between action values and action perseverations.

##### Model-free strategy with inertia (d=3)

The action values are updated as the model-free strategy (d=2). The action perseveration (inertia) is updated by the same rule in the model-free strategy with inertia (d=2).

The action probabilities in all model-free models with inertia are generated via softmax({*β*(*Q*_*t*_(Ai) + *X*_*t*_(*A*_*i*_))}_*i*_). Both the action values and action perseverations are dynamical variables.

##### Model-free reward-as-cue strategy (d=8)

This model assumes that the animal considers the combination of the second-stage state *s*_t−1_ and the reward *r*_*t*−1_ from the trial *t* − 1 as the augmented state *S*_*t*_ for trial *t*. The eight dynamical variables are the values for the two actions at the four augmented states. The action values are updated as follows:

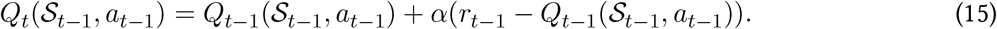

The action probability at trial *t* is determined by softmax(*βQ*_*t*_(*S*_*t*_, *A*_1_), *βQ*_*t*_(*S*_*t*_, *A*_2_)).

#### Models for the two-stage task

We implemented one model from the Bayesian inference family, four models from the model-free family, and four from the model-based family (adopted from^36^ and^13^).

##### Bayesian inference strategy (d=1)

Same as Bayesian inference strategy (d=1) in the reversal learning task, except that *h = i* represents a higher reward probability following state *S*_*i*_ (not action *A*_*i*_).

##### Model-free strategy (d=1)

Same as the model-free strategy (d=1) in the reversal learning task by ignoring the second-stage states *s*_*t*−1_.

##### Model-free Q(1) strategy (d=2)

Same as the model-free strategy (d=2) in the reversal learning task by ignoring the second-stage states *s*_*t*−1_.

##### Model-free Q(0) strategy (d=4)

This model first updates the first-stage action values *Q*_*t*_(*a*_*t*−1_) with the second-stage state values *V*_t−1_(*s*_*t*−1_):

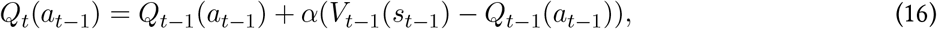

while the unchosen action value *Q*_*t*_(*ā*_*t*−1_) is unaffected. Then the second-stage state value *V*_*t*_(*s*_t−1_) is updated by the observed reward:

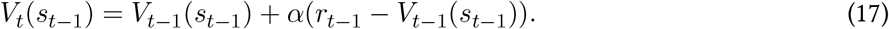

The four dynamical variables are the two action values and two state values.

##### Model-free reward-as-cue strategy (d=8)

Same as model-free reward-as-cue strategy (d=8) in the reversal learning task.

##### Model-based strategy (d=1)

In this model, the two state values *V*_*t*_(*S*_*i*_) are fully anti-correlated (*V*_*t*_(*S*_1_) = −*V*_*t*_(*S*_2_)):

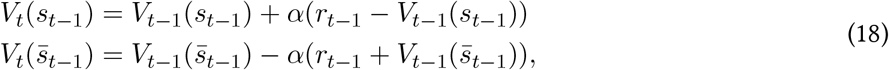

where 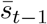 is the unvisited state. The dynamical variable is the state value *V*_*t*_(*S*_1_).

##### Model-based strategy (d=2)

The visited state value is updated:

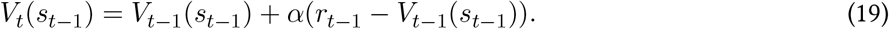

The unvisited state value is unchanged. The two dynamical variables are the two state values.

##### Model-based strategy with value forgetting (d=2)

The visited state value is updated as in the previous model. The unvisited state value is gradually forgotten:

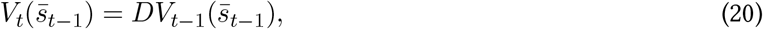

where *D* is the value forgetting rate (0 ≤ *D* ≤ 1).

For all model-based RL models, the action values at the first stage are directly computed using the state transition model:

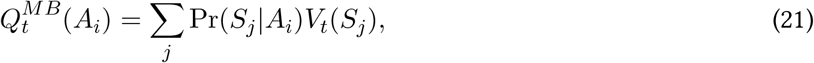

where *P*_*r*_(*S*_*j*_ |*A*_*i*_) is known. The action probability is determined by softmax 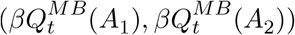.

##### Model-based mixture strategy (d=2)

model-based strategy (d=1). The net action values are determined by:

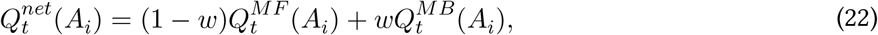

where *w* controls the strength of the model-based component. The action probabilities are generated via softmax 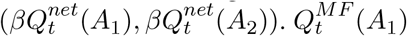 and *V*_*t*_ (*S*_1_) are the dynamical variables.

#### Models for the transition-reversal two-stage task

For this task, we further include cognitive models proposed in^12^. We first describe different model components (ingredients) and corresponding numbers of dynamical variables, and then specify the components employed in each model.

##### Second-stage state value component

The visited state value is updated:

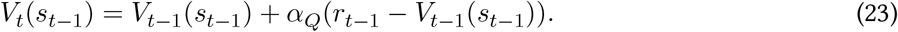

The unvisited state value 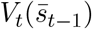 is either unchanged or gradually forgotten with *f*_*Q*_ as the value forgetting rate. This component requires two dynamical variables.

##### Model-free action value component

The first-stage action values 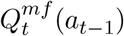 are updated by the second-stage state values *V*_*t*−1_(*s*_*t*−1_) and the observed reward:

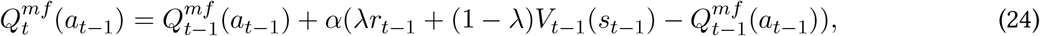

where λ is the eligibility trace. The unchosen action value 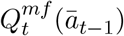 is unaffected or gradually forgotten with *f*_*Q*_ as the value forgetting rate. This component requires two dynamical variables.

##### Model-based component

The action-state transition probabilities are updated as:

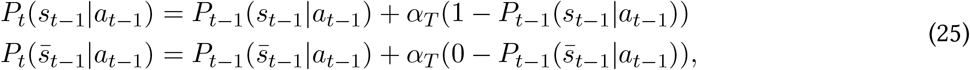

where *α*_*T*_ is the transition probability learning rate. For the unchosen action, the action-state transition probabilities are either unchanged or forgotten:

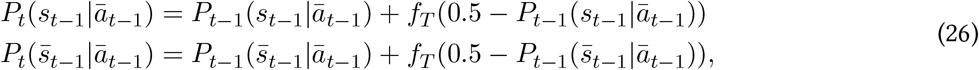

where *f*_*T*_ is the transition probability forgetting rate.

The model-based action values at the first stage are directly computed using the learned state transition model:

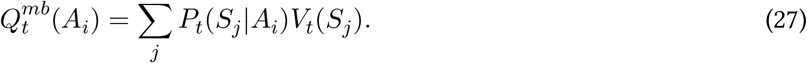

This component requires two dynamical variables (*P*_*t*_(*S*_1_|*A*_1_) and *P*_*t*_(*S*_1_|*A*_2_)), since other variables can be directly inferred.

##### Motor-level model-free action component

Due to the apparatus design in this task^12^, it is proposed that the mice consider the motor-level actions 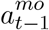, defined as the combination of the last-trial action *a*_t−1_ and the second-stage state *s*_t−2_ before it. The motor-level action values 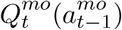 are updated as:

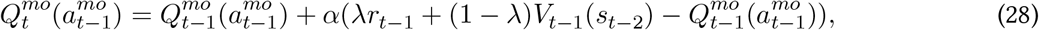

where *λ* is the eligibility trace. The unchosen motor-level action value 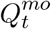 is unaffected or gradually forgotten with *f*_*Q*_ as the value forgetting rate. This component requires four dynamical variables (four motor-level actions).

##### Choice perseveration component

The single-trial perseveration 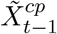 is set to -0.5 for *a*_t−1_ = *A*_1_ and 0.5 for *a*_t−1_ = *A*_2_. The multi-trial perseveration 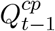 (exponential moving average of choices) is updated as:

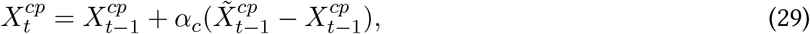

where *α*_*c*_ is the choice perseveration learning rate. In some models, the *α*_*c*_ is less than 1, so one dynamical variable is required; while in some other models, the *α*_*c*_ is fixed to 1, suggesting that it is reduced to the single-trial perseveration and no dynamical variable is required.

##### Motor-level choice perseveration component

The multi-trial motor-level perseveration 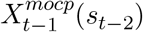 is updated as:

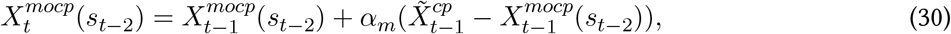

where *α*_*m*_ is the motor-level choice perseveration learning rate. This component requires two dynamical variables.

##### Action selection component

The net action values are computed as follows:

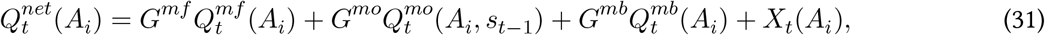

where *G*^*mf*^, *G*^*mo*^, *G*^mb^ are model-free, motor-level model-free, model-based inverse temperatures, and *X*_*t*_(*A*_*i*_) is:

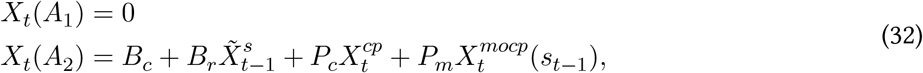

where *B*_*c*_ (bias), *B*_*r*_ (*r*otation bias), *P*_*c*_, Pm are weights controlling each component, and 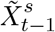 is -0.5 for *s*_*t* −1_ = *S*_1_and 0.5 for *s*_t−1_ = *S*_2_.

The action probabilities are generated via softmax 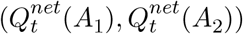

##### Model-free strategies

We include five model-free RL models:

- the model-free strategy (d=1) same as the two-stage task;
- the model-free Q(1) strategy (d=2) same as the two-stage task;
- state value [2] + model-free action value [2] + bias [0] + rotation bias [0] + single-trial choice perseveration [0];
- state value [2] + model-free action value with forgetting [2] + bias [0] + rotation bias [0] + single-trial choice perseveration [0];
- state value [2] + model-free action value with forgetting [2] + motor-level model-free action value with forgetting [4] + bias [0] + rotation bias [0] + multi-trial choice perseveration [1] + multi-trial motor-level choice perseveration [2].

Here, we use the format of “model component [required number of dynamical variables]” (more details in^12^).

##### Model-based strategies

We include twelve model-based RL models:

- state value [2] + model-based [2] + bias [0] + rotation bias [0] + single-trial choice perseveration [0];
- state value [2] + model-free action value [2] + model-based [2] + bias [0] + rotation bias [0] + single-trial choice perseveration [0];
- state value [2] + model-based with forgetting [2] + bias [0] + rotation bias [0] + single-trial choice perseveration [0];
- state value [2] + model-free action value with forgetting [2] + model-based with forgetting [2] + bias [0] + rotation bias [0] + single-trial choice perseveration [0];
- state value [2] + model-free action value with forgetting [2] + model-based [2] + bias [0] + rotation bias [0] + single-trial choice perseveration [0];
- state value [2] + model-free action value [2] + model-based [2] + bias [0] + rotation bias [0] + multi-trial choice perseveration [1];
- state value [2] + model-free action value with forgetting [2] + model-based with forgetting [2] + bias [0] + rotation bias [0] + multi-trial choice perseveration [1];
- state value [2] + model-free action value with forgetting [2] + model-based [2] + bias [0] + rotation bias [0] + multi-trial choice perseveration [1];
- state value [2] + model-free action value with forgetting [2] + model-based with forgetting [2] + bias [0] + rotation bias [0] + multi-trial motor-level choice perseveration [2];
- state value [2] + model-based with forgetting [2] + bias [0] + rotation bias [0] + multi-trial choice perseveration [1] + multi-trial motor-level choice perseveration [2];
- state value [2] + model-free action value with forgetting [2] + model-based with forgetting [2] + bias [0] + rotation bias [0] + multi-trial choice perseveration [1] + multi-trial motor-level choice perseveration [2];
- state value [2] + model-free action value with forgetting [2] + model-based with forgetting [2] + motor-level model-free action value with forgetting [4] + bias [0] + rotation bias [0] + multi-trial choice perseveration [1] + multi-trial motor-level choice perseveration [2].

Here, we use the format of model component [required number of dynamical variables] (more details in^12^).

#### Models for the three-armed reversal learning task

We implemented four models (*n* = 3 actions) from the model-free family, one of which is constructed from the strategies discovered by the GRU.

##### Model-free strategy (d=n)

This model hypothesizes that each action value *Q*_*t*_(*A*_*i*_), as a dynamical variable, is updated independently. The chosen action value is updated by:

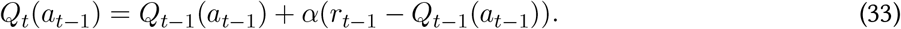

The unchosen action values *Q*_*t*_(*A*_*j*_) (*A*_*j*_≠*a*_t−1_) are unaffected.

##### Model-free strategy with value forgetting (d=n)

The chosen action value is updated as in the previous model. The unchosen action value *Q*_*t*_(*A*_*j*_) (*A*_*j*_≠*a*_t−1_), instead, is gradually forgotten:

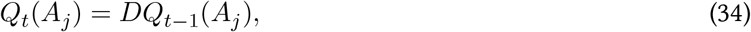

where *D* is the value forgetting rate (0 ≤ *D* ≤ 1).

##### Model-free strategy with value forgetting and action perseveration (d=2n)

The action values are updated as the model-free strategy with value forgetting. The chosen action perseveration is updated by:

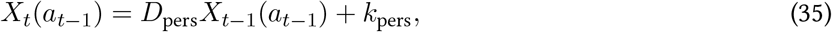

and the unchosen action perseverations are updated by:

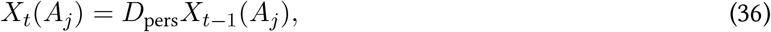

where *D*_pers_ is the perseveration forgetting rate (0 ≤ *D*_pers_ ≤ 1), and *k*_pers_ is the single-trial perseveration term, affecting the balance between action values and action perseverations.

##### Model-free strategy with unchosen value updating and reward utility (d=n)

This model is constructed from the strategy discovered by the GRU (see Supplementary Results 1.4). It assumes that the reward utility *U* (*r*) (equivalent to the preference setpoint) is different in four cases (corresponding to four free parameters): no reward for chosen action (*Uc*(0)), one reward for chosen action (*Uc*(1)), no reward for unchosen action (*Uu*(0)), and one reward for chosen action (*Uu*(1)).

The chosen action value is updated by:

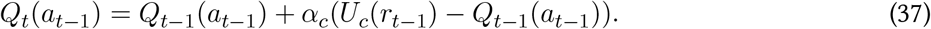

The unchosen action value *Q*_*t*_(*A*_*j*_) (*A*_*j*_≠*a*_*t*−1_) is updated by:

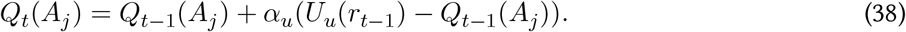

The action probabilities for these models are generated via softmax({*β*(*Q*_*t*_(*A*_*i*_) + *X*_*t*_(*A*_*i*_))}_*i*_) (*X*_*t*_ = 0 for models without action perseverations). Both the action values and action perseverations are dynamical variables.

#### Models for the four-armed drifting bandit task

We implemented five models (*n* = 4 actions) from the model-free family, two of which are constructed from the strategies discovered by the GRU.

##### Model-free strategy (d=n)

This model is the same as the model-free strategy in the three-armed reversal learning task.

##### Model-free strategy with value forgetting (d=n)

This model is the same as the model-free strategy with value forgetting in the three-armed reversal learning task.

##### Model-free strategy with value forgetting and action perseveration (d=2n)

This model is the same as the model-free strategy with value forgetting and action perseveration in the three-armed reversal learning task.

##### Model-free strategy with unchosen value updating and reward reference point (d=n)

This model is constructed from the strategy discovered by the GRU (see Supplementary Results 1.5). It assumes that the reward utility *U* (*r*) is different for chosen action (*U*_*c*_(*r*) = *β*_*c*_(*r* − *R*_*c*_)) and for unchosen action (*U*_*u*_(*r*) = *β*_*u*_(*r* − *R*_*u*_)), where *β*_*c*_ and *β*_*u*_ are reward sensitivities, and *R*_*c*_ and *R*_*u*_ are reward reference points.

The chosen action value is updated by:

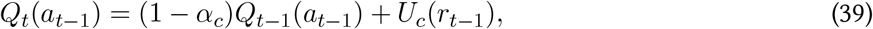

where 1 − *α*_*c*_ is the decay rate for chosen actions. The unchosen action value *Q*_*t*_(*A*_*j*_) (*A*_*j*_≠*a*_*t*−1_) is updated by:

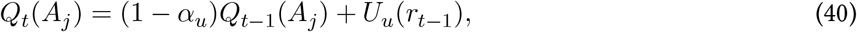

where 1 − *α*_*u*_ is the decay rate for unchosen actions. We additionally fit a reduced model of this strategy where *β*_*c*_ = *α*_*c*_ and *β*_*u*_ = *α*u (similarly inspired by the GRU’s solution).

The action probabilities for these models are generated via softmax({*β*(*Q*_*t*_(*A*_*i*_) + *X*_*t*_(*A*_*i*_))}_*i*_) (*X*_*t*_ = 0 for models without action perseverations). Both the action values and action perseverations are dynamical variables.

#### Models for the original two-stage task

##### Model-free strategy (d=3)

This model hypothesizes that the action values for each task state (first-stage state *S*_*0*_, second-stage states *S*_*1*_ and *S*_*2*_) are fully anti-correlated 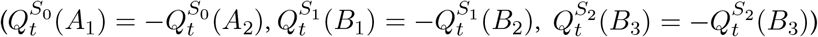

The action values at the chosen second-stage state (e.g., assuming *B*_1_ or *B*_2_ at *S*_1_ is chosen) are updated by:

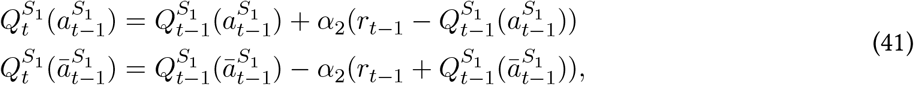

where 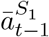 is the unchosen second-stage action at the chosen second-stage state, and *α*_2_ is the learning rate for the second-stage states (0 ≤ *α*_2_ ≤_1_). The second-stage action probabilities are generated via softmax 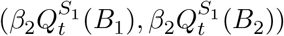

The action values at the first-stage state (*A*_1_ or *A*_2_ at *S*_0_) are updated by:

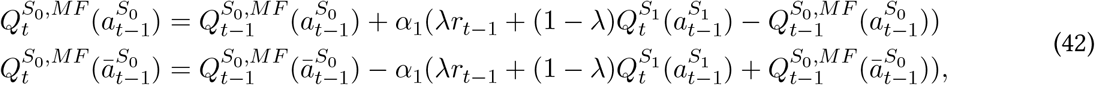

where 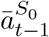 is the unchosen first-stage action, *α*_1_ is the learning rate for the first-stage state (0 ≤ *α*_1_ ≤_1_), and λ specifies the (λ) learning rule. The first-stage action probabilities are generated via softmax 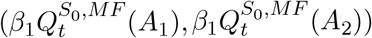

Here 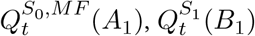 and 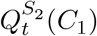 are the dynamical variables.

##### Model-based strategy (d=2)

The update of action values at the chosen second-stage state is the same as the model-free strategy. The action values at the first-stage state (*A*_1_ or *A*_2_ at *S*_0_) are determined by:

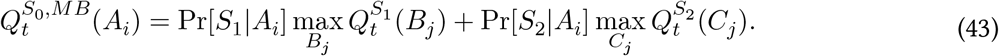

The first-stage action probabilities are generated via softmax 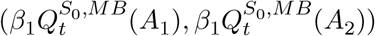 only 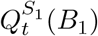 and 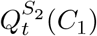 are the dynamical variables.

##### Model-based mixture strategy (d=3)

This model considers the mixture of model-free and model-based strategies for the first-stage states. The net action values are determined by:

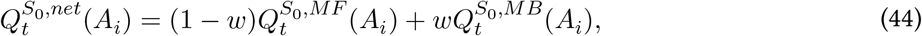

where w controls the strength of the model-based component. The first-stage action probabilities are generated via softmax 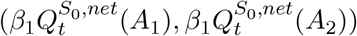

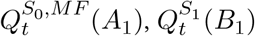 and 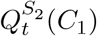 are the dynamical variables

##### Model-free strategy (d=6)

Compared to the model-free strategy (d=3), only the chosen action values at *S*_*0*_, *S*_1_, and *S*_2_ are updated. The unchosen values are unchanged. 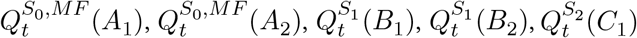 and 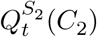 are the dynamical variables.

##### Model-based strategy (d=4)

Compared to the model-based strategy (d=2), only the chosen action values at *S*_1_, and *S*_2_ are updated. The unchosen values are unchanged 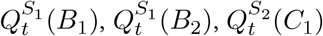 and 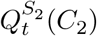 are the dynamical variables.

##### Model-based mixture strategy (d=6)

Compared to the model-based mixture strategy (d=3), only the chosen action values at *S*_0_, *S*_1_, and *S*_2_ are updated. The unchosen values are unchanged. 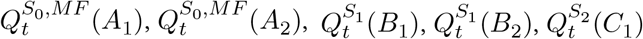 and 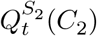 are the dynamical variables.

##### Model-free strategy with reward utility (d=3)

This model is constructed from the GRU’s strategy. Similar to the model-free strategy (d=3), it hypothesizes that the action values for each task state (first-stage state *S*_0_, second-stage states *S*_1_ and *S*_2_) are fully anti-correlated 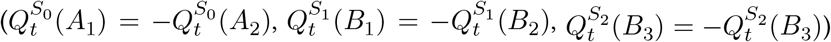

It assumes that when receiving one reward, the reward utility (i.e., equivalently, the preference setpoint) for the chosen action at the first-stage state *S*_0_ is 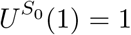, for the chosen action at the chosen second-stage state *S*_1_ (or *S*_2_) is 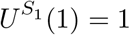, and for the (motor-level) chosen action at the unchosen second-stage state *S*_2_ (or *S*_1_) is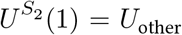 (e.g., *B*_1_ at the chosen *S*_1_ and *C*_1_ at unchosen *S*_2_ are the same motor-level action). When receiving no reward, the reward utility for the chosen action at the first-stage state *S*_0_ is 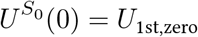 for the chosen action at the chosen second-stage state (assuming *S*_1_) is 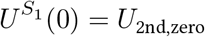, and for the (motor-level) chosen action at the unchosen second-stage state (assuming *S*_2_) is 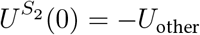 The chosen action values at the chosen second-stage state (e.g., assuming B1 or B2 at S1) are updated by:

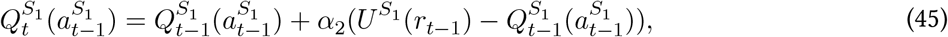

where *α*_2_ is the learning rate for the second-stage states (0 ≤ *α*_2_ ≤_1_). The (motor-level) chosen action values (i.e., 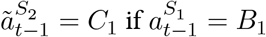 and, 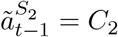 if 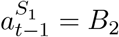) at the unchosen second-stage state (e.g., assuming *C*_1_ or *C*_2_ at *S*_2_) are updated by:

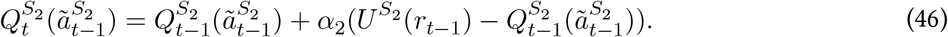

The second-stage action probabilities are generated via softmax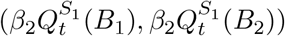

The action values at the first-stage state (*A*_1_ or *A*_2_ at *S*_0_) are updated by:

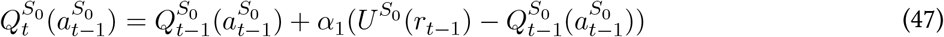

where *α*_1_ is the learning rate for the first-stage state (0 ≤ *α*_1_ ≤_1_). The first-stage action probabilities are generated via softmax, 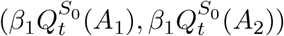

Here, 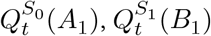 and, 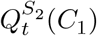 are the dynamical variables.

### Model fitting

#### Maximum likelihood estimation

The parameters in all models were optimized on the training dataset to maximize the log-likelihood (i.e., minimize the negative log-likelihood, or cross-entropy) for the next-action prediction. The loss function is defined as follows:

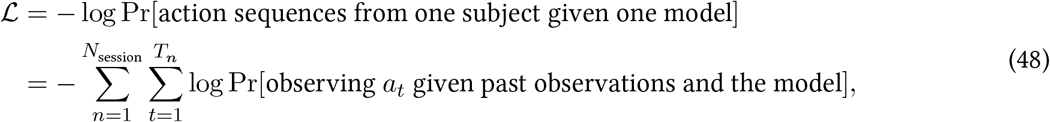

where N_session_ is the number of sessions and T_*n*_ is the number of trials in session *n*.

#### Nested cross-validation

To avoid overfitting and ensure a fair comparison between models with varying numbers of parameters, we implemented nested cross-validation. For each animal, we first divided sessions into non-overlapping shorter blocks (approximately 150 trials per block) and allocated these blocks into ten folds. In the outer loop, nine folds were designated for training and validation, while the remaining fold was reserved for testing. In the inner loop, eight of the nine folds were assigned for training (optimizing a model’s parameters for a given set of hyperparameters), and the remaining fold of the nine was allocated for validation (selecting the best-performing model across all hyperparameter sets). Notice that this procedure allows different hyperparameters for each test set.

RNNs’ hyperparameters encompassed the L1-regularization coefficient on recurrent weights (drawn from 1e-5, 1e-4, 1e-3, 1e-2, 1e-1, depending on the experiments), the number of training epochs (i.e., early stopping), and the random seed (three seeds). For cognitive models, the only hyperparameter was the random seed (used for parameter initialization). The inner loop produced nine models, with the best-performing model, based on average performance in the training and validation datasets, being selected and evaluated on the unseen testing fold. The final testing performance was computed as the average across all ten testing folds, weighted by the number of trials per block. This approach ensures that test data is exclusively used for evaluation and is never encountered during training or selection.

During RNN training, we employed early stopping if the validation performance failed to improve after 200 training epochs. This method effectively prevents RNN overfitting on the training data. According to this criterion, a more flexible model may demonstrate worse performance than a less flexible one, as the training for the former could be halted early due to insufficient training data. However, it is expected that the more flexible model would continue to improve with additional training data (e.g., see Fig. *S*_8_).

We note that, in the rich-data situation, this training-validation-test split in (nested) cross-validation is better than the typical usage of Akaike information criteria (AIC)^55^, corrected AIC (AICc)^56^, or Bayesian information criteria (BIC)^57^ in cognitive modeling, due to the following reasons^58^: the (nested) cross-validation provides a direct and unbiased estimate of the expected extra-sample test error, which reflects the generalization performance on new data points with inputs not necessarily appearing in the training dataset; in contrast, AIC, AICc, and BIC can only provide asymptotically unbiased estimates of in-sample test error under some conditions (e.g., models are linear in their parameters), measuring the generalization performance on new data points with inputs always appearing in the training dataset (the labels could be different from those in the training dataset due to noise). Furthermore, in contrast to regular statistical models, neural networks are singular statistical models with degenerate Fisher information matrices. Consequently, estimating the model complexity (the number of effective parameters, as used in AIC/*A*ICc/*B*IC) in neural networks requires estimating the real log canonical threshold^59^, which falls outside the scope of this study.

##### Estimating the dimensionality of behavior

For each animal, we observed that the predictive performance of RNN models initially improves and then saturates or even declines as the number d of dynamical variables increases. To operationally estimate the dimensionality *d*_∗_ of behavior, we implemented a statistical procedure that satisfies two criteria: (1) the RNN model with *d*_∗_ dynamical variables significantly outperforms all RNN models with *d*_<_ *d*_∗_ dynamical variables (using a significance level of 0.05 in the t-tests of predictive performance conducted over outer folds); (2) any RNN model with *d*_′_ (*d*^′^ > *d*_∗)_ dynamical variables does not exhibit significant improvement over all RNN models with *d* ^<^ *d*_′_ dynamical variables.

Our primary objective is to estimate the intrinsic dimensionality (*r*eflecting the latent variables in the data-generating process), not the embedding dimensionality^60^. However, it is important to consider the practical limitations associated with the estimation procedure. For instance, RNN models may fail to uncover certain latent variables due to factors such as limited training data or variables operating over very long time scales, leading to an underestimation of *d*_∗._ Additionally, even if all *d*_∗_ latent variables are accurately captured, the RNN models may still require *d*_≥_ *d*_∗_ dynamical variables to effectively and losslessly embed d∗-dimensional dynamics, particularly if they exhibit high nonlinearity, potentially resulting in an overestimation of *d*_∗._ A comprehensive understanding of these factors is crucial for future studies.

##### Knowledge distillation

We employ the knowledge distillation framework^35^ to fit models to individual subjects, while simultaneously leveraging group data: first fitting a teacher network to data from multiple subjects, and then fitting a student network to the outputs of the teacher network corresponding to an individual subject.

#### Teacher network

In the teacher network (TN), each subject is represented by a one-hot vector. This vector projects through a fully connected linear layer into a subject-embedding vector *e*_sub_, which is provided as an additional input to the RNN. The TN uses 20 units in its hidden layer and uses the same output layer and loss (cross-entropy between the next-trial action and the predicted next-trial action probability) as in previous GRU models.

#### Student network

The student network (SN) has the same architecture as previous tiny GRUs. The only difference is that, during training and validation, the loss is defined as cross-entropy between the next-trial action probability provided by the teacher and the next-trial action probability predicted by the student:

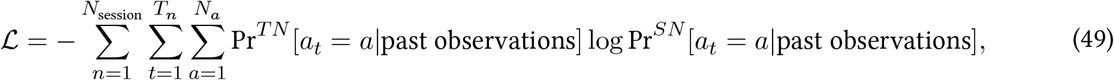

where *N*_*session*_ is the number of sessions, *T*_*n*_ is the number of trials in session *n*, and *N*_*a*_ is the number of actions (*N*_*a*_ = 2 in our tasks).

#### Training, validation, and test data in knowledge distillation for the mouse in the Akam dataset

To study the influence of the number of training trials from one representative mouse on the performance of knowledge distillation, we employed a procedure different from nested cross-validation. This procedure splits the data from animal *M* into two sets. The first set consisted of 25% of the trials and was used as a hold-out M-test dataset. The second set consisted of the remaining 75% trials, from which smaller datasets of different sizes were sampled. From each sampled dataset, 90% of the trials were used for training (M-training dataset) and 10% for validation (M-validation dataset). Next, we split the data from all other animals, with 90% of the data used for training (O-training dataset) and 10% for validation (O-validation dataset).

After dividing the datasets as described above, we trained the models. The solo GRUs were trained to predict choices on the M-training dataset and selected on the M-validation dataset. The teacher GRUs were trained to predict choices on the M- and O-training datasets and selected on the M- and O-validation datasets. The number of embedding units in the teacher GRUs was selected based on the M-validation dataset. The student GRUs were trained on the M-training dataset and selected on the M-validation dataset, but with the training target of action probabilities provided by the teacher GRUs. Here the student GRUs and the corresponding teacher GRUs were trained on the same M-training dataset. Finally, all models were evaluated on the unseen M-test data.

When training the student GRUs, due to symmetry in the task, we augment the M-training datasets by flipping the action and second-stage states, resulting in an augmented dataset that is four times the size of the original one, similar to^30^. One key difference between our augmentation procedure and that of^30^ is that the authors augmented the data for training the group RNNs, where the potential action bias presented in the original dataset (and other related biases) becomes invisible to the RNNs. In contrast, our teacher RNNs are trained only on the original dataset, where any potential action biases can be learned. Even if we augment the training data later for the student networks, the biases learned by the teacher network can still be transferred into the student networks. In addition to direct augmentation, simulating the teacher network can be another method to generate pseudo-data. The benefit of these pseudo-data was discussed in model compression^61^.

### Protocols for training, validating, and testing models in human datasets

#### Interspersed split protocol

In the three human datasets, each subject only performs one block of 100-200 trials. In the standard practice of cognitive modeling, the cognitive models are trained and tested on the same block, leading to potential overfitting and exaggerated performance. While it is possible to directly segment one block into three sequences for training, validation, and testing, this might introduce undesired distributional shifts in the sequences due to the learning effect.

To ensure a fair comparison between RNNs and cognitive models, here we propose a new interspersed split protocol to define the training, validation, and testing trials, similar to the usage of goldfish loss to prevent the memorization of training data in language models^62^. Specifically, we randomly sample without replacement ∼ 75% trial indexes for training, ∼ 12.5% trial indexes for validation, and ∼ 12.5% trial indexes for testing (three-armed reversal learning task: 120/20/20; four-armed drifting bandit task: 110/20/20; original two-stage task: 150/25/25). We then feed in the whole block of trials as the model’s inputs, obtain the output probabilities for each trial, and calculate the training, validation, and testing losses for each set of trial indexes, separately. This protocol guarantees the identical distribution between three sets of trials.

One possible concern is whether the test data is leaked into the training data in this protocol. For instance, the models are trained on the input sequence ((*a*_1_, *r*_1_), (*a*_2_, *r*_2_), (*a*_3_, r3)) to predict a4 and later tested on the input sequence ((*a*_1_, *r*_*1*_), (*a*_2_, r2)) to predict *a*_3_. In this scenario, while the models see a3 in the input during training, they never see *a*_3_ in the output. Thus, models are not trained to learn the input-output mapping from ((*a*_1_, *r*_1_), (*a*_2_, *r*_2_)) to *a*_3_, which is evaluated during testing. We confirmed that this procedure prevents data leakage on artificially generated choices (Fig. S38).

#### Cross-subject split protocol

In addition to the interspersed split protocol, it is possible to train the RNNs on a proportion of subjects and evaluate them on held-out subjects (i.e., zero-shot generalization), a cross-subject split protocol. To illustrate this protocol, we first divided all subjects into 6 folds of cross-validation. The teacher network was trained and validated using 5 folds and tested on the remaining 1 fold. For each subject in the test fold, because each subject only completed one task block, student networks are trained on the action-augmented blocks (to predict the teacher’s choice probabilities for the subject), validated on the original block (to predict the teacher’s choice probabilities for the subject), and tested on the original block (to predict actual choices of the subject). By design, both teacher networks and student networks will not overfit the subjects’ choices in the test data. The cognitive models were trained and validated using 5 folds and tested on the remaining 1 fold. We presented the results in Fig. S39).

### Phase portraits

#### Models with d = 1

##### Logit

In each trial t, a model predicts the action probabilities Pr(*a*_*t*_ = *A*_1_) and Pr(*a*_*t*_ = *A*_2_). We define the logit *L*(*t*) (log-odds) at trial *t* as *L*(*t*) = log (Pr(*a*_*t*_ = *A*_1_)/ Pr(*a*_*t*_ = *A*_2_)). When applied to probabilities computed via softmax, the logit yields 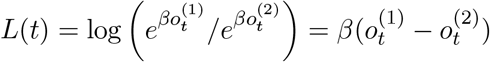 where 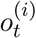 is the model’s output for action *a*_*t*_ = *A*_*i*_ before softmax. Thus, the logit can be viewed as reflecting the preference for action *A*_1_ over *A*_2_: in RNNs, the logit corresponds to the score difference 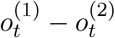 in model-free and model-based RL models, the logit is proportional to the difference in first-stage action values *Q*_*t*_(*A*_1_) − *Q*_*t*_(*A*_2_); in Bayesian inference models, the logit is proportional to the difference in latent-state probabilities Pr_*t*_(*h* = 1) − Pr_*t*_(*h* = 2) = 2 Pr_*t*_(*h* = 1) − 1.

##### Logit change

We define the logit change, ΔL(*t*), in trial t as the difference between L(t + 1) and L(*t*). In one-dimensional models, ΔL(*t*) is a function of the input and L(*t*), forming a vector field.

##### Stability of fixed points

Here we derive the stability of a fixed point in one-dimensional discrete dynamical systems. The system’s dynamics update according to:

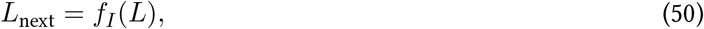

where *L* is the current-trial logit, *L*_next_ is the next-trial logit, and *f*_*I*_ is a function determined by input *I* (omitted for simplicity). At a fixed point, denoted by *L* = *L*^∗^, we have

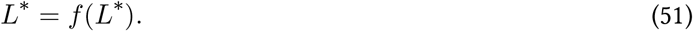

Next, we consider a small perturbation δL around the fixed point:

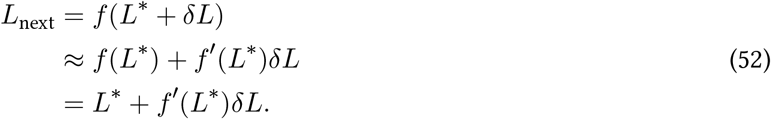

The fixed point is stable only when −1 < *f*′(*L*∗) < 1. Because the logit change Δ*L* is defined as Δ*L* = *g*(*L*) = *f* (*L*) − *L*, we have the stability condition −2 < *g*′(*L*∗) < 0.

##### Effective learning rate and slope

In the one-dimensional RL models with prediction error updates and constant learning rate *α*, we have

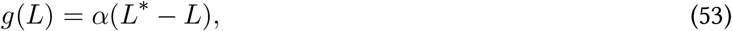

where *g*(*L*) is the logit change at *L*. In general, to obtain a generalized form of *g*(*L*) = *α*(*L*)(*L*− *L*) with a non-constant learning rate, we define the effective learning rate *α*(*L*) at *L* relative to a stable fixed point *L*∗ as:

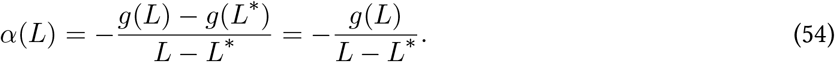

At *L*∗, *α*(*L*∗) is the negative slope −*g*′(*L*∗) of the tangent at *L*∗. However, for general *L*≠*L*∗, *α*(*L*) is the negative slope of the secant connecting (*L, g*(*L*)) and (*L*∗, 0), which is different from −*g*′(*L*).

We have

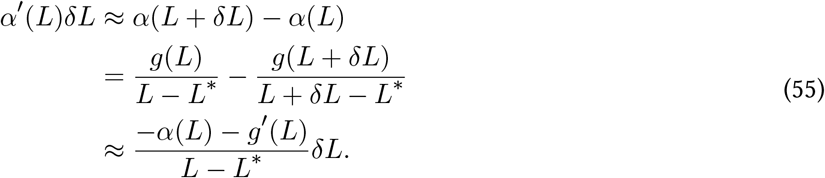

Letting δ*L* go to zero, we have:

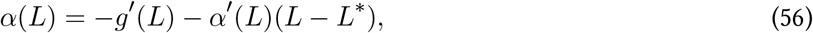

which provides the relationship between the effective learning rate *α*(L) and the slope of the tangent *g*′(*L*).

##### Models with d > 1

In models with more dynamical variables, Δ*L*(*t*) is no longer solely a function of the input and *L*(*t*) due to added degrees of freedom. In these models, the state space is spanned by a set of dynamical variables, collected by the vector *F*(*t*). For example, the action value vector is the *F* (*t*) = (*Q*_*t*_(*A*_1_), *Q*_*t*_(*A*_2_))^*T*^ in the two-dimensional RL models. The vector field ΔF (*t*) can be defined as ΔF (*t*) = *F* (*t* +1) − *F* (*t*) = (*Q*_*t*_+1(*A*_1_) − *Q*_*t*_(*A*_1_), *Q*t+_1_(*A*_2_) − *Q*_*t*_(*A*_2_))^*T*^, a function of *F* (*t*) and the input in trial t.

#### Dynamical regression

For one-dimensional models with states characterized by the policy logit *L*(*t*), we can approximate the one-step dynamics for a given input with a linear function, i.e., Δ_*L*_ ∼ *β*_0_ + *β*_*L*_*L*. The coefficients *β*_0_ and *β*_*L*_ can be computed via linear regression, or “dynamical regression” given its use in modeling dynamical systems. Here, *β*_0_ is similar to the preference setpoint and *βL* is similar to learning rates in RL models.

For models with more than one dynamical variable, we can use a similar dynamical regression approach to extract a first-order approximation of the model dynamics via linearizations of vector fields. To facilitate interpretation, we consider only d-dimensional RNNs with a d-unit diagonal readout layer (denoted by *Li*(*t*) or *Pi*(*t*); a non-degenerate case).

For tasks with a single choice state (Supplementary Results 1.4, 1.5), the diagonal readout layer means that d is equal to the number of actions. Thus Pi(*t*) corresponds to the action preference for Ai at trial t (before softmax). A special case of *P*_*i*_(*t*) is equal to *βV*_*t*_(*A*_*i*_) in cognitive models. We use Δ *P*_*i*_ (*t*) = *P*_*i*_(*t* + 1) − *P*_*i*_(*t*) to denote preference changes between two consecutive trials. For the reversal learning task and three-armed reversal learning task, we consider Δ*P*_*i*_(*t*) as an (approximate) linear function of *P*_1_(*t*), …, *P*_*d*_(*t*) for different (discrete) task inputs (i.e., 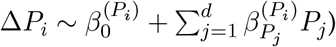 For the four-armed drifting bandit task, we further include the continuous reward r as an independent variable (i.e., 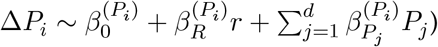

For the original two-stage task, where there are 3 choice states (Supplementary Result 1.6), we focus on the 3-dimensional model with a diagonal readout layer. Here, *L*_1_, *L*_2_, and *L*_3_ represent the logits for *A*_1_/*A*_2_ at the first-stage state, logits for *B*_1_/*B*_2_ at the second-stage state *S*_1_ and logits for *C*_1_/*C*_2_ at the second-stage state *S*_2_, respectively. We similarly consider the regression 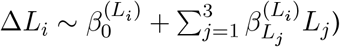

Collecting all the 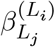 (similarly for 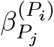 regression coefficients for a given input condition, we have the input-dependent state-transition matrix A, akin to the Jacobian matrix of nonlinear dynamical systems:

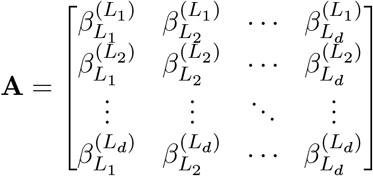

Note that the model-free RL models in these tasks are fully characterized by the collection of all regression coefficients in our dynamical regression.

#### Symbolic regression

Apart from the two-dimensional vector field analysis, symbolic regression is another method for discovering concise equations that summarize the dynamics learned by GRUs. To accomplish this, we used PySR 63 to search for simple symbolic expressions of the updated dynamical variables as functions of the current dynamical variables for each possible input I (for the GRU with d = 2 and a diagonal readout matrix). Ultimately, this process revealed a model-free strategy featuring the “drift-to-the-other” rule.

#### Model validation via behavior-feature identifier

We proposed a general and scalable approach based on a “behavior-feature identifier”. In contrast to conventional model recovery, this approach provides a model-agnostic form of validation to identify and verify the hallmark of the discovered strategy in the empirical data.

For a given task, we collect the behavioral sequences generated by models that exhibit a specific feature (positive class) and by those that do not (negative class). An RNN identifier is then trained on these sequences to discern their classes. Subsequently, this identifier is applied to the actual behavioral sequences produced by subjects.

We built identifiers to distinguish between the GRU models (positive class) and model-free RL models (negative class) in the reversal learning task, and between the GRU models (positive class) and model-based RL models (negative class) in the two-stage task. We presented the results in Fig. S27).

#### Meta-reinforcement learning models

We trained meta-RL agents on the two-stage task (common transition: Pr(*S*_1_|*A*_1_) = Pr(*S*_2_|*A*_2_) = 0.8, rare transition: Pr(*S*_2_|*A*_1_) = Pr(*S*_1_|*A*_2_) = 0.2; see Fig. S25) implemented in NeuroGym 64. Each second-stage state leads to a different probability of a unit reward, with the most valuable state switching stochastically (Pr(*r* = 1|*S*_1_) = 1 − Pr(*r* = 1|*S*_2_) = 0.8 or 0.2 with a probability of 0.025 on each trial). There are three periods (discrete time steps) on one trial: Delay 1, Go, and Delay 2. During Delay 1, the agent receives the observation (choice state S0 and a fixation signal), and the reward (1 or 0) from second-stage states on the last trial. During Go, the agent receives the observation of the choice state and a go signal. During Delay 2, the agent receives the observation of state *S*_1_/*S*_2_ and a fixation signal. If the agent does not select action *A*_1_ or *A*_2_ during Go or select action *F* (Fixate) during Delay periods, a small negative reward (−0.1) is given. The contributions of second-stage states, rewards, and actions on networks are thus separated in time.

The agent architecture is a fully connected, gated recurrent neural network (long short-term memory^51^) with 48 units^24^. The input to the network consists of the current observation (state *S*_0_/*S*_1_/*S*_2_ and a scalar fixation/go signal), a scalar reward signal of the previous time step, and a one-hot action vector of the previous time step. The network outputs a scalar baseline (value function for the current state) serving as the critic and a real-valued action vector (passed through a softmax layer to sample one action from *A*_1_/*A*_2_/*F*) serving as the actor. The agents are trained using the Advantage Actor-Critic RL algorithm 65 with the policy gradient loss, value estimate loss, and entropy regularization. We trained and analyzed agents for five seeds. Our agents obtained 0.64 rewards on average on each trial (0.5 rewards for chance level), close to optimal performance (0.68 rewards obtained by an oracle agent knowing the correct action).

## References

[1] Allen Newell and Herbert A Simon. Computer science as empirical inquiry: Symbols and search. In ACM Turing award lectures, page 1975. 2007.

[2] James L McClelland, David E Rumelhart, PDP Research Group, et al. Parallel Distributed Processing, Volume 2: Explorations in the Microstructure of Cognition: Psychological and Biological Models, volume 2. MIT press, 1987.

[3] Joshua B Tenenbaum, Charles Kemp, Thomas L Griffiths, and Noah D Goodman. How to grow a mind: Statistics, structure, and abstraction. science, 331(6022):1279–1285, 2011.

[4] Brenden M Lake, Tomer D Ullman, Joshua B Tenenbaum, and Samuel J Gershman. Building machines that learn and think like people. Behavioral and brain sciences, 40:e253, 2017.

[5] Richard S Sutton and Andrew G Barto. Reinforcement learning: An introduction. MIT press, 2018.

[6] Yael Niv. Reinforcement learning in the brain. Journal of Mathematical Psychology, 53(3):139–154, 2009.

[7] Matthew Botvinick, Sam Ritter, Jane X Wang, Zeb Kurth-Nelson, Charles Blundell, and Demis Hassabis. Reinforcement learning, fast and slow. Trends in cognitive sciences, 23(5):408–422, 2019.

[8] Anne Gabrielle Eva Collins. Reinforcement learning: Bringing together computation and cognition. Current Opinion in Behavioral Sciences, 29:63–68, 2019.

[9] Paul Cisek and John F Kalaska. Neural mechanisms for interacting with a world full of action choices. Annual review of neuroscience, 33:269–298, 2010.

[10] Marcelo G Mattar and Máté Lengyel. Planning in the brain. Neuron, 2022.

[11] Ramon Bartolo and Bruno B Averbeck. Prefrontal cortex predicts state switches during reversal learning. Neuron, 106(6):1044–1054, 2020.

[12] Thomas Akam, Ines Rodrigues-Vaz, Ivo Marcelo, Xiangyu Zhang, Michael Pereira, Rodrigo Freire Oliveira, Peter Dayan, and Rui M Costa. The anterior cingulate cortex predicts future states to mediate model-based action selection. Neuron, 109(1):149–163, 2021.

[13] Kevin J Miller, Matthew M Botvinick, and Carlos D Brody. Value representations in the rodent orbitofrontal cortex drive learning, not choice. Elife, 11:e64575, 2022.

[14] Robert C Wilson and Anne GE Collins. Ten simple rules for the computational modeling of behavioral data. Elife, 8:e49547, 2019.

[15] Carolina Feher da Silva and Todd A Hare. Humans primarily use model-based inference in the two-stage task. Nature Human Behaviour, 4(10):1053–1066, 2020.

[16] Blake A Richards, Timothy P Lillicrap, Philippe Beaudoin, Yoshua Bengio, Rafal Bogacz, Amelia Christensen, Claudia Clopath, Rui Ponte Costa, Archy de Berker, Surya Ganguli, et al. A deep learning framework for neuroscience. Nature neuroscience, 22(11):1761–1770, 2019.

[17] Guangyu Robert Yang and Xiao-Jing Wang. Artificial neural networks for neuroscientists: a primer. Neuron, 107(6):1048–1070, 2020.

[18] Daniel LK Yamins, Ha Hong, Charles F Cadieu, Ethan A Solomon, Darren Seibert, and James J DiCarlo. Performance-optimized hierarchical models predict neural responses in higher visual cortex. Proceedings of the national academy of sciences, 111(23):8619–8624, 2014.

[19] Daniel LK Yamins and James J DiCarlo. Using goal-driven deep learning models to understand sensory cortex. Nature neuroscience, 19(3):356–365, 2016.

[20] Christopher J Cueva and Xue-Xin Wei. Emergence of grid-like representations by training recurrent neural networks to perform spatial localization. arXiv preprint arXiv:1803.07770, 2018.

[21] Andrea Banino, Caswell Barry, Benigno Uria, Charles Blundell, Timothy Lillicrap, Piotr Mirowski, Alexander Pritzel, Martin J Chadwick, Thomas Degris, Joseph Modayil, et al. Vector-based navigation using grid-like representations in artificial agents. Nature, 557(7705):429–433, 2018.

[22] James CR Whittington, Timothy H Muller, Shirley Mark, Guifen Chen, Caswell Barry, Neil Burgess, and Timothy EJ Behrens. The tolman-eichenbaum machine: unifying space and relational memory through generalization in the hippocampal formation. Cell, 183(5):1249–1263, 2020.

[23] Valerio Mante, David Sussillo, Krishna V Shenoy, and William T Newsome. Context-dependent computation by recurrent dynamics in prefrontal cortex. nature, 503(7474):78–84, 2013.

[24] Jane X Wang, Zeb Kurth-Nelson, Dharshan Kumaran, Dhruva Tirumala, Hubert Soyer, Joel Z Leibo, Demis Hassabis, and Matthew Botvinick. Prefrontal cortex as a meta-reinforcement learning system. Nature neuroscience, 21(6):860–868, 2018.

[25] Valeria Fascianelli, Fabio Stefanini, Satoshi Tsujimoto, Aldo Genovesio, and Stefano Fusi. Neural representational geometry correlates with behavioral differences between monkeys. bioRxiv, pages 2022–10, 2022.

[26] Guangyu Robert Yang, Madhura R Joglekar, H Francis Song, William T Newsome, and Xiao-Jing Wang. Task representations in neural networks trained to perform many cognitive tasks. Nature neuroscience, 22(2): 297–306, 2019.

[27] Kristopher T Jensen, Guillaume Hennequin, and Marcelo G Mattar. A recurrent network model of planning explains hippocampal replay and human behavior. bioRxiv, pages 2023–01, 2023.

[28] Joshua C Peterson, David D Bourgin, Mayank Agrawal, Daniel Reichman, and Thomas L Griffiths. Using large-scale experiments and machine learning to discover theories of human decision-making. Science, 372 (6547):1209–1214, 2021.

[29] Amir Dezfouli, Kristi Griffiths, Fabio Ramos, Peter Dayan, and Bernard W Balleine. Models that learn how humans learn: The case of decision-making and its disorders. PLoS computational biology, 15(6):e1006903, 2019.

[30] Mingyu Song, Yael Niv, and Mingbo Cai. Using recurrent neural networks to understand human reward learning. In Proceedings of the Annual Meeting of the Cognitive Science Society, volume 43, 2021.

[31] Paul I Jaffe, Russell A Poldrack, Robert J Schafer, and Patrick G Bissett. Modelling human behaviour in cognitive tasks with latent dynamical systems. Nature Human Behaviour, pages 1–15, 2023.

[32] Kyunghyun Cho, Bart Van Merriënboer, Dzmitry Bahdanau, and Yoshua Bengio. On the properties of neural machine translation: Encoder-decoder approaches. arXiv preprint arXiv:1409.1259, 2014.

[33] William Bialek. On the dimensionality of behavior. Proceedings of the National Academy of Sciences, 119(18): e2021860119, 2022.

[34] Surya Ganguli. Measuring the dimensionality of behavior. Proceedings of the National Academy of Sciences, 119(43):e2205791119, 2022.

[35] Jianping Gou, Baosheng Yu, Stephen J Maybank, and Dacheng Tao. Knowledge distillation: A survey. International Journal of Computer Vision, 129:1789–1819, 2021.

[36] Thomas Akam, Rui Costa, and Peter Dayan. Simple plans or sophisticated habits? state, transition and learning interactions in the two-step task. PLoS computational biology, 11(12):e1004648, 2015.

[37] Adrian Valente, Jonathan W Pillow, and Srdjan Ostojic. Extracting computational mechanisms from neural data using low-rank rnns. Advances in Neural Information Processing Systems, 35:24072–24086, 2022.

[38] Francesca Mastrogiuseppe and Srdjan Ostojic. Linking connectivity, dynamics, and computations in low-rank recurrent neural networks. Neuron, 99(3):609–623, 2018.

[39] Manuel Beiran, Alexis Dubreuil, Adrian Valente, Francesca Mastrogiuseppe, and Srdjan Ostojic. Shaping dynamics with multiple populations in low-rank recurrent networks. Neural Computation, 33(6):1572–1615, 2021.

[40] Alexis Dubreuil, Adrian Valente, Manuel Beiran, Francesca Mastrogiuseppe, and Srdjan Ostojic. The role of population structure in computations through neural dynamics. Nature neuroscience, 25(6):783–794, 2022.

[41] Christopher Langdon and Tatiana A Engel. Latent circuit inference from heterogeneous neural responses during cognitive tasks. bioRxiv, pages 2022–01, 2022.

[42] Boyuan Chen, Kuang Huang, Sunand Raghupathi, Ishaan Chandratreya, Qiang Du, and Hod Lipson. Automated discovery of fundamental variables hidden in experimental data. Nature Computational Science, 2(7): 433–442, 2022.

[43] Pantelis R Vlachas, Georgios Arampatzis, Caroline Uhler, and Petros Koumoutsakos. Multiscale simulations of complex systems by learning their effective dynamics. Nature Machine Intelligence, 4(4):359–366, 2022.

[44] Ian D Jordan, Piotr Aleksander Sokół, and Il Memming Park. Gated recurrent units viewed through the lens of continuous time dynamical systems. Frontiers in computational neuroscience, 15:678158, 2021.

[45] Ramon Bartolo, Richard C Saunders, Andrew R Mitz, and Bruno B Averbeck. Dimensionality, information and learning in prefrontal cortex. PLoS computational biology, 16(4):e1007514, 2020.

[46] Praveen Suthaharan, Erin J Reed, Pantelis Leptourgos, Joshua G Kenney, Stefan Uddenberg, Christoph D Mathys, Leib Litman, Jonathan Robinson, Aaron J Moss, Jane R Taylor, et al. Paranoia and belief updating during the covid-19 crisis. Nature human behaviour, 5(9):1190–1202, 2021.

[47] Bahador Bahrami and Joaquin Navajas. 4 arm bandit task dataset. 10.17605/OSF.IO/F3T2A, 2020.

[48] Claire M Gillan, Michal Kosinski, Robert Whelan, Elizabeth A Phelps, and Nathaniel D Daw. Characterizing a psychiatric symptom dimension related to deficits in goal-directed control. elife, 5:e11305, 2016.

[49] Nathaniel D Daw, Samuel J Gershman, Ben Seymour, Peter Dayan, and Raymond J Dolan. Model-based influences on humans’ choices and striatal prediction errors. Neuron, 69(6):1204–1215, 2011.

[50] Nathaniel D Daw, John P O’doherty, Peter Dayan, Ben Seymour, and Raymond J Dolan. Cortical substrates for exploratory decisions in humans. Nature, 441(7095):876–879, 2006.

[51] Sepp Hochreiter and Jürgen Schmidhuber. Long short-term memory. Neural computation, 9(8):1735–1780, 1997.

[52] Scott W Linderman, Andrew C Miller, Ryan P Adams, David M Blei, Liam Paninski, and Matthew J Johnson. Recurrent switching linear dynamical systems. arXiv preprint arXiv:1610.08466, 2016.

[53] Yuhuai Wu, Saizheng Zhang, Ying Zhang, Yoshua Bengio, and Russ R Salakhutdinov. On multiplicative integration with recurrent neural networks. Advances in neural information processing systems, 29, 2016.

[54] Anne GE Collins and Michael J Frank. How much of reinforcement learning is working memory, not reinforcement learning? a behavioral, computational, and neurogenetic analysis. European Journal of Neuroscience, 35(7):1024–1035, 2012.

[55] Hirotugu Akaike. A new look at the statistical model identification. IEEE transactions on automatic control, 19(6):716–723, 1974.

[56] Joseph E Cavanaugh. Unifying the derivations for the akaike and corrected akaike information criteria. Statistics & Probability Letters, 33(2):201–208, 1997.

[57] Gideon Schwarz. Estimating the dimension of a model. The annals of statistics, pages 461–464, 1978.

[58] Trevor Hastie, Robert Tibshirani, Jerome H Friedman, and Jerome H Friedman. The elements of statistical learning: data mining, inference, and prediction, volume 2. Springer, 2009.

[59] Sumio Watanabe. A widely applicable bayesian information criterion. The Journal of Machine Learning Research, 14(1):867–897, 2013.

[60] Mehrdad Jazayeri and Srdjan Ostojic. Interpreting neural computations by examining intrinsic and embedding dimensionality of neural activity. Current opinion in neurobiology, 70:113–120, 2021.

[61] Cristian Buciluă, Rich Caruana, and Alexandru Niculescu-Mizil. Model compression. In Proceedings of the 12th ACM SIGKDD international conference on Knowledge discovery and data mining, pages 535–541, 2006.

[62] Abhimanyu Hans, Yuxin Wen, Neel Jain, John Kirchenbauer, Hamid Kazemi, Prajwal Singhania, Siddharth Singh, Gowthami Somepalli, Jonas Geiping, Abhinav Bhatele, et al. Be like a goldfish, don’t memorize! mitigating memorization in generative llms. arXiv preprint arXiv:2406.10209, 2024.

[63] Miles Cranmer. Pysr: Fast & parallelized symbolic regression in python/julia. September 2020. doi: 10.5281/zenodo.4041459. URL 10.5281/zenodo.4041459.

[64] Manuel Molano-Mazon, Joao Barbosa, Jordi Pastor-Ciurana, Marta Fradera, Ru-Yuan Zhang, Jeremy Forest, Jorge del Pozo Lerida, Li Ji-An, Christopher J Cueva, Jaime de la Rocha, et al. Neurogym: An open resource for developing and sharing neuroscience tasks. 2022.

[65] Volodymyr Mnih, Adria Puigdomenech Badia, Mehdi Mirza, Alex Graves, Timothy Lillicrap, Tim Harley, David Silver, and Koray Kavukcuoglu. Asynchronous methods for deep reinforcement learning. In International conference on machine learning, pages 1928–1937. PMLR, 2016.

