## Supplementary Materials for "Discovering Cognitive Strategies with Tiny Recurrent Neural Networks"

### Supplementary Materials
### Automatic Discovery of Cognitive Strategies with Tiny Recurrent Neural Networks

Li Ji-An[1], Marcus K. Benna[2], and Marcelo G. Mattar[3, @]

[1]Neurosciences Graduate Program, University of California San Diego, La Jolla, CA 92093
[2]Department of Neurobiology, School of Biological Sciences, University of California San Diego, La Jolla, CA 92093
[3]Department of Psychology, New York University, New York, NY 10012

#### 1 Supplementary Results

##### 1.1 Tiny RNNs can identify ground-truth strategies in simulated behavior

What if biological agents exhibited a different behavioral pattern other than observed in these datasets, such as those exhibited by an RL or Bayesian inference agent? One possibility is that RNN models can only approximate, but not fully account for, the behavior of such agents. In this scenario, a cognitive model well-matched to the behavior would outperform the RNN models. Alternatively, the RNN models might be flexible enough to fully account for the behavior of an RL or Bayesian inference agent. In this scenario, the RNN models may be viewed as a superset of the cognitive models, and their predictive performance would be no worse than that of any cognitive model.

To determine how well can an RNN model predict the behavior of RL and Bayesian inference agents, we simulated the behavior of such agents over 10,000 choices in the reversal learning task (with parameters fitted to one example monkey from the Bartolo dataset) and the two-stage task (with parameters fitted to one example rat from the Miller dataset). By design, thus, these choices embodied the behavior of the RL, Bayesian inference, or GRU agents with one or two dynamical variables. We then fitted all cognitive models and RNN models to optimize predictive performance for this simulated behavior. We found that the best RNN models achieved identical predictive performance as the ground-truth model of equal dimensionality that generated the behavior (Fig. S4, S5, S6, S7).

A further concern arises from the identifiability of strategies that RNNs learn in these tasks, given the limited choice data available. Specifically, there is a possibility that RNNs could learn various strategies that equally well fit identical sequences of behavior. In such scenarios, it becomes challenging to determine whether strategies learned by the RNNs accurately reflect the underlying ground-truth strategies that generate the behavior. However, our findings indicate that these RNNs indeed accurately recovered the ground-truth model's dynamics (see Fig.S11), suggesting the identifiability of the strategies under study is not problematic.

#### 1.2 Detailed analysis of the logit diagram in the reversal learning task

An important detail in the GRU diagram ($d = 1$) is the non-overlapping curves representing state changes after no reward (light blue and light red curves in Fig. 3c, compared to the model-free model predictions in Fig. 3b). To understand what this means, note that choice perseveration in these diagrams would be diagnosed by an upward displacement of the two curves corresponding to $A_1$ (indicating an increase in preference for $A_1$ regardless of the outcome), and by a downward displacement of the two curves corresponding to $A_2$ (indicating an increase in preference for $A_2$ regardless of the outcome). In the GRU diagrams, the two curves representing no reward are only non-overlapping when the monkey highly favors one of the two options, suggesting a logit-dependent choice perseveration.

This rather nuanced pattern of behavior, while was identified from the phase portrait of a one-unit GRU, can also be confirmed by directly interrogating the data. By reanalyzing the relations between next-trial action probability and the current-trial logit in the behavioral data, we confirmed the existence of such a logit-dependent choice perseveration effect (Fig. S14a-d). Similar results were also obtained for the second monkey (Fig. S14e-h).

#### 1.3 Analysis of the switching linear neural networks

In addition to the GRUs (vanilla and switching GRUs) studied in the main text, we found that the switching linear neural networks (SLIN, see Methods) can achieve comparable performance to the GRUs in some animals and agents (equipped with different cognitive strategies) (Fig. S22), providing alternative transparent descriptions of the underlying cognitive strategies given their linear updating equations.

In SLINs, given an input $I$, the hidden state $h_t$ ($t > 1$) is updated as follows (we omit the subscript $I$ for simplicity):

$$h_t = Wh_{t-1} + b \tag{S1}$$

where $W$ and $b$ are the input-dependent weight matrices and biases (see Methods). The logit is read out as $r^T h_t$, where $r$ is the readout vector. Note that this model is invariant under some transformations, such as rotating or scaling $h_t$, $b$, and $r$ simultaneously.

To analyze the dynamics of SLINs, we can consider the singular value decomposition of $W$. Here we constrain $W$ to be symmetric and consider its eigenvalue decomposition for two practical reasons: we observed that SLINs with symmetry constraints usually demonstrate comparable performance to those without constraints, and the RL models can be treated as SLINs with diagonal weight matrices.

In the two-dimensional case, the fixed point is given by $h_* = (I - W)^{-1}b$. The stability of the fixed point is determined by the two eigenvalues of $W$, $\lambda_1$ and $\lambda_2$ ($\lambda_1 \geq \lambda_2$):

1. if $|\lambda_1| < 1$ and $|\lambda_2| < 1$, then $h_*$ is an attractor;
2. if $|\lambda_1| = 1$ and $|\lambda_2| < 1$, then there is a line attractor;
3. if $|\lambda_1| > 1$ or $|\lambda_2| > 1$, then $h_*$ is a saddle point (unstable);
4. if $|\lambda_1| > 1$ and $|\lambda_2| > 1$, then $h_*$ is a repellor.

Specifically, for symmetric $W$, the two eigenvalues $\lambda_1$ and $\lambda_2$ are always real numbers and we can write $W$ as $W = \lambda_1 v_1 v_1^T + \lambda_2 v_2 v_2^T$, where $v_i$ are the associated eigenvectors and are mutually orthogonal. Therefore, the projection of $h_t$ on $v_i$ evolves as follows:

$$
\begin{aligned}
h_t^{(i)} &= \langle v_i, h_t \rangle \\
&= \lambda_i \langle v_i, h_{t-1} \rangle + \langle v_i, b \rangle \\
&= \lambda_i h_{t-1}^{(i)} + b^{(i)},
\end{aligned} \tag{S2}
$$

where $\langle \cdot, \cdot \rangle$ is the inner product. Importantly, $h_t^{(1)}$ and $h_t^{(2)}$ evolve independently, similar to the RL models with $v_1 = (1, 0)^T$, $v_2 = (0, 1)^T$, and $\lambda_i$ act as decaying rates. In the RL models, the readout weight $r = (\beta, -\beta)^T$, where $\beta$ is the inverse temperature, is $45^\circ$ away from either eigenvector. In contrast, the angles between the readout weight and eigenvectors in the general SLINs are not constrained.

The symmetric SLINs fitted to one example rat performing the two-stage task (Fig. S23) and one example mouse performing the transition-reversal two-stage task (Fig. S24) show that the locations of fixed points (relative to the decision boundary) are consistent with one-dimensional results. An approximate line attractor occurs when $\lambda_1 \approx 1$.

For the rat performing the two-stage task, the eigenvectors $v_2$ associated with the smaller $\lambda_2$ are always aligned with the readout vector, suggesting that this eigenvector direction is conceptually similar to the logit. This result indicates that these two-dimensional dynamics can be considered as one-dimensional logit dynamics modulated by the dynamics on the orthogonal eigenvector $v_1$.

For the mouse performing the transition-reversal two-stage task, we found that the angles between the readout vector and the eigenvectors might change for different input conditions. This suggests a more complex strategy this mouse used, evading all classical cognitive models with $d = 2$.

##### 1.4 Dynamical regression analysis of the three-armed reversal learning task

In this section we describe dynamical regression coefficients for three-dimensional models fitted to subjects' behavior in the three-armed reversal learning task. $P_i$ and $\Delta P_i$ represents the preference and the preference change for action $A_i$ ($i = 1, 2, 3$), respectively. We consider the regression $\Delta P_i \sim \beta_0^{(P_i)} + \beta_{P_1}^{(P_i)} P_1 + \beta_{P_2}^{(P_i)} P_2 + \beta_{P_3}^{(P_i)} P_3$ for each input condition.

In the model-free model with value forgetting (Fig. S28), only the chosen action value is influenced by the reward ($\beta_0 > 0$ for $\Delta P_1$ in [A1 R=1] trials, $\Delta P_2$ in [A2 R=1] trials, $\Delta P_3$ in [A3 R=1] trials; the magnitude of $\beta_0$ is related to the inverse temperature of each subject), but not the unchosen action values ($\beta_0 = 0$ for all other coefficients). However, in the GRU model (Fig. S29), we observed diverse $\beta_0^{(P_i)}$ coefficients across trial conditions and subjects. The features of $\beta_0$ in the GRU model are summarized as follows:

(1) $\beta_0$ is mostly larger than zero for the chosen action value after receiving a reward (for $\Delta P_1$ in [A1 R=1] trials, for $\Delta P_2$ in [A2 R=1] trials, for $\Delta P_3$ in [A3 R=1] trials), suggesting a positive impact on the chosen action value;

(2) $\beta_0$ is non-zero for the chosen action value after receiving no reward (for $\Delta P_1$ in [A1 R=0] trials, for $\Delta P_2$ in [A2 R=0] trials, for $\Delta P_3$ in [A3 R=0] trials), and is on average smaller than $\beta_0$ for the chosen action value after receiving a reward, suggesting either a positive or negative impact on the chosen action value (depending on the subject);

(3) $\beta_0$ is mostly smaller than zero for the unchosen action value after receiving a reward (for $\Delta P_1$ in [A2

R=1] & [A3 R=1] trials, for $\Delta P_2$ in [A1 R=1] & [A3 R=1] trials, for $\Delta P_3$ in [A1 R=1] & [A2 R=1] trials), suggesting a negative impact on the unchosen action values;

(4) $\beta_0$ is non-zero for the unchosen action value after receiving no reward (for $\Delta P_1$ in [A2 R=0] & [A3 R=0] trials, for $\Delta P_2$ in [A1 R=0] & [A3 R=0] trials, for $\Delta P_3$ in [A1 R=0] & [A2 R=0] trials), suggesting either a positive or negative impact on the unchosen action values (depending on the subject).

In short, receiving a reward will mostly positively impact the chosen action value and negatively impact the unchosen action value, but receiving no reward can either positively or negatively impact action values for all actions, with the directions dependent on the subjects.

For other coefficients, we found that $\beta_{P_i}$ for $\Delta P_j$ $(i \neq j)$ in the GRU model is close to zero, suggesting weak (or zero) cross-action dependence. $\beta_{P_i}$ for $\Delta P_i$ lies between 0 and -1, suggesting value decaying in all trial conditions. These are similar to the assumptions in the model-free RL models.

We also visualized the regression coefficients of a new model-free RL model (model-free strategy with unchosen value updating and reward utility (d=3)) inspired by GRU models (see Methods and Fig. S30). By design, it updates both the chosen action values and the unchosen action values (the non-zero $\beta_0$ for all trial conditions and $\Delta P_i$). This model achieves a test loss of 0.477, outperforming the best existing model (model-free strategy) with a test loss of 0.602.

##### 1.5 Dynamical regression analysis of the four-armed drifting bandit task

In this section we describe the dynamical regression coefficients for four-dimensional models fitted to subjects' behavior in the four-armed drifting bandit task. $P_i$ and $\Delta P_i$ represents the preference and the preference change for action $A_i$ $(i = 1, 2, 3, 4)$, respectively. We consider the regression $\Delta P_i \sim \beta_0^{(P_i)} + \beta_R^{(P_i)} r + \beta_{P_1}^{(P_i)} P_1 + \beta_{P_2}^{(P_i)} P_2 + \beta_{P_3}^{(P_i)} P_3 + \beta_{P_4}^{(P_i)} P_4$ for each input condition.

In the model-free model with value forgetting (Fig. S31), only the chosen action value is updated by the reward ($\beta_R > 0$ for $\Delta P_i$ in $[A_i]$ trials; the magnitude of $\beta_R$ is related to the inverse temperature of each subject), but not the unchosen action values ($\beta_R = 0$ for all other coefficients). However, in the GRU model (Fig. S32), we observed that the reward (mostly) positively impacts the chosen action value ($\beta_R > 0$ for $\Delta P_i$ in $[A_i]$ trials) and (usually) negatively impacts the unchosen action values ($\beta_R$ usually smaller than zero for $\Delta P_i$ in $[A_j]$ trials $(j \neq i)$). Additionally, we observed non-zero $\beta_0$ across all trial conditions in the GRU model, suggesting a non-zero reference point effect (because $\beta_0^{(P_i)} + \beta_R^{(P_i)} r$ can be transformed into $\beta_R^{(P_i)}(r - \beta_0^{(P_i)}/\beta_R^{(P_i)})$, where $\beta_0^{(P_i)}/\beta_R^{(P_i)}$ is a reference point).

In addition, we found that $\beta_{P_i}$ for $\Delta P_j$ $(i \neq j)$ in the GRU model is close to zero, suggesting weak (or zero) cross-action dependence. $\beta_{P_i}$ for $\Delta P_i$ lies between 0 and -1, suggesting value decaying in all trial conditions. These are similar to the assumptions in the model-free RL models.

We also visualized the regression coefficients of a new model-free RL model (model-free strategy with unchosen value updating and reward reference point (d=4)) inspired by GRU models (see Methods and Fig. S33). By design, it updates both the chosen action values and unchosen action values with rewards shifted by reference points (non-zero $\beta_0$ and $\beta_R$ for all trial conditions and $\Delta P_i$), capturing major features of the GRU model. This model achieves a test loss of 0.796, outperforming the best existing model (model-free strategy with value forgetting and action perseveration) with a test loss of 0.815.

##### 1.6 Dynamical regression analysis of the original two-stage task

In this section we describe dynamical regression coefficients for three-dimensional models fitted to subjects' behavior in the original two-stage task. $L_1$ represents the logits for $A_1/A_2$ at the first-stage state, $L_2$ represents logits for $B_1/B_2$ at the second-stage state $S_1$, and $L_3$ represents logits for $C_1/C_2$ at the second-stage state $S_2$. we consider $\Delta L_i \sim \beta_0^{(L_i)} + \sum_{j=1}^{3} \beta_{L_j}^{(L_i)} L_j$ for each input condition.

The comparison of the regression coefficients for $\Delta L_1$ in the model-free RL model and the GRU model is summarized below.

(1) For $\Delta L_1$, both the model-free RL model and the GRU model shows $\beta_0 > 0$ for [A1 R=1] trials (positively impacting the preference for $A_1$) and $\beta_0 < 0$ for [A2 R=1] trials (negatively impacting the preference for $A_1$). The GRU model additionally shows that the magnitude of $\beta_0$ for [A1 S2 R=1] & [A2 S1 R=1] trials (both are rare transitions) are smaller than that for [A1 S1 R=1] & [A2 S2 R=1] trials (both are common transitions), suggesting a modulation effect of task transitions. The model-free RL model shows $\beta_0 = 0$ for [R=0] trials, but GRU model shows $\beta_0 > 0$ for [A1 R=0] trials (positively impacting the preference for $A_1$) and $\beta_0 < 0$ for [A2 R=0] trials (negatively impacting the preference for $A_1$), suggesting non-zero impact on the preference for $A_1$ (not modulated by the state S1/S2).

(2) For $\Delta L_1$, both the model-free RL model and the GRU model shows $0 > \beta_{L_1} > -1$, suggesting typical value decaying.

(3) For $\Delta L_1$, both the model-free RL model and the GRU model shows that $\beta_{L_2}$ and $\beta_{L_3}$ are close to zero for most subjects, suggesting that the $L_1$ is mostly not affected by $L_2$ or $L_3$ (i.e., TD(1)-like).

(4) For $\Delta L_2$, both the model-free RL model and the GRU model shows that $\beta_0 > 0$ for [B1 R=1] trials and $\beta_0 < 0$ for [B2 R=1] trials. The model-free RL model shows $\beta_0 = 0$ for [R=0] trials, but GRU model shows $\beta_0 < 0$ for [B1 R=0] trials (negatively impacting the preference for $B_1$) and $\beta_0 > 0$ for [B2 R=0] trials (positively impacting the preference for $B_1$). Notice that the direction of the impact is opposite to the $\beta_0$ for $\Delta L_1$ (e.g., comparing [A1 S1 B1] for $\Delta L_1$ and for $\Delta L_2$).

(5) For $\Delta L_2$, interestingly, the GRU model discovers that taking action C1/C2 in state S2 has a small impact on the logit $L_2$ for state S1: $\beta_0$ is positive for [S2 C1 R=1] and [S2 C2 R=0] trials, and negative for [S2 C1 R=0] and [S2 C2 R=1] trials. This pattern is similar to $\beta_0$ for $\Delta L_3$ (e.g., comparing [S2] trials for $\Delta L_2$ and for $\Delta L_3$), suggesting a motor-level influence (different actions on different second-stage states sharing the same motor-level action).

(6) For $\Delta L_2$, the model-free RL model shows $\beta_{L_2} < 0$ only for [S1] trials, but the GRU model shows $\beta_{L_2} < 0$ for all trials, suggesting value decaying no matter which second-stage states are experienced.

The comparisons for $\Delta L_3$ are omitted since they are mostly symmetric to $\Delta L_2$.

We also visualized the regression coefficients of a new model-free RL model (model-free strategy with reward utility (d=3)) inspired by the GRU model (see Methods and Fig. S36). This model, which incorporates most of the above features discovered by the GRU model, achieves a test loss of 0.448, outperforming the best existing cognitive model (model-based mixture strategy (d=3)) with a test loss of 0.535.

#### 2 Supplementary Discussion

##### 2.1 Interpretation and insights from discrete dynamical systems approach for the animal datasets

Our results also suggest that animals treated the presence and absence of reward differently (Fig. 3). This contradicts the ideal-observer behavior assumed in Bayesian inference models where an animal should leverage its knowledge of the task structure to infer that the absence of reward for one action indicates the presence of reward for the other action. One possible reason for the asymmetrical treatment of reward is that animals were never shown the reward for the unchosen action, and thus were unable to learn the symmetric nature of the task. Another possibility is that animals exhibited a "reward-induced indifference effect" whereby uncertainty is reduced by confirmatory evidence (e.g., a reward when a reward is expected) but increased otherwise (e.g., a lack of reward when a reward is expected). This explanation is indeed consistent with the suboptimal behavior commonly observed in perceptual decision tasks where animals process past success and failure differently[1], reflecting qualitatively different neural mechanisms triggered by outcomes with valence[2]. Our analyses of the two-stage task data provide further support for a reward-induced indifference effect (Fig. 3e). Here, when a rare transition leads to a reward, animals do not increase their preference for the chosen (model-free) or the unchosen (model-based and Bayesian inference) actions, but rather become increasingly indifferent. Overall, our results suggest that, in the three examined tasks but possibly also many others, the modulatory effect of reward on behavior may depend on past events, including past actions and observations.

Additionally, we observed a novel pattern of state-dependent choice perseveration in the reversal learning task (Fig. 3c), where monkeys tended to repeat an action regardless of the reward outcome, but only when they displayed a high preference for either action. Note that we identified such a pattern simply by visualizing the logit dynamics, without the requirement of a prior hypothesis or the specification of an augmented behavioral model, as traditionally done[3]. The pattern of state-dependent perseveration could then be confirmed using standard hypothesis testing (Fig. S14).

##### 2.2 Emergent algorithms in meta learning

Our analysis of meta-RL agents trained for the two-stage task has revealed that they learned a Bayesian-inference-like representation, and updated the internal variables using Bayesian inference. While prior research has suggested that such meta-trained agents learn either a model-based RL algorithm[4] or a Bayes-optimal algorithm[5–7], we have strictly tested these competing hypotheses together. Our logit analysis visualizes the dynamics of the meta-RL agents in detail, leading us to discover that the emergent Bayesian inference algorithm is imperfect and negatively affected by the history effect in the recurrent dynamics. Identifying emergent algorithms in this way provides a principled approach for "opening the black box" of meta-trained agents.

An intriguing question is whether these meta-RL agents can learn suboptimal behavior similar to that of animals under certain circumstances. Previous research has shown that RNNs can replicate rats' suboptimal behavior in the two-alternative forced-choice task when equipped with adaptive structural priors implemented by pre-training in a more naturalistic environment with multiple alternative choices, providing normative explanations for animals' behavior[1]. Identifying the naturalistic environmental factors that give rise to the emergence of animal-like strategies in meta-RL agents would be an interesting direction for future research.

##### 2.3 Justification for counting dynamical variables in the models

One may question whether only one variable is needed to pass information about the past into the future. While it is true that one can pack several *discrete* variables into a single variable, the same is not true about *continuous* variables — i.e., it is *not* possible to pack multiple continuous variables into one continuous variable without significant loss of precision. Intuitively, well-behaved functions (e.g., smooth functions) cannot unpack multiple continuous variables from a single continuous variable without exhibiting pathological behavior. Below we present two theorems with proofs to support this claim.

**Theorem 1.** *A one-dimensional continuous RNN $h_{t+1} = f(h_t; Input)$ with a linear readout, cannot accurately represent a two-dimensional dynamical system without significant precision loss.*

*Proof.* Consider a two-dimensional RL model where the value of the chosen actions depend on reward prediction error, and the value of unchosen action remains unchanged (Fig. S40a). The model's initial state is $(Q_L, Q_R) = (0, 0)$, and the first inputs alternate between [$A_L$;R=1] and [$A_R$ R=1] five times, for a total of 10 trials. Given that the readout from the one-dimensional hidden state $h_t$ of the RNN to the output layer is linear, $h_t$ can *only* represent a linear function of the logit $L_t$, i.e., $h_t = aL_t + b$, where $a$ and $b$ are coefficients and $L_t = \log[\Pr(A_L)/\Pr(A_R)]$. At the red and blue points explored by the RL model, since $Q_L = Q_R$, $L_t = 0$ (indicating equal probability), resulting in $h_t = b$. This implies that red and blue points should represented approximately at $b$ in the RNN ($h_{red} \approx h_{blue} \approx b$). By continuity, we have $f(h_{red}; A_L\ R = 1) \approx f(h_{blue}; A_L\ R = 1) \approx f(b; A_L\ R = 1)$. However, $h_{orange} = f(h_{red}; A_L\ R = 1)$, which strongly favors $Q_L$, should differ from $h_{green} = f(h_{blue}; A_L\ R = 1)$, which shows almost equal preference. This disparity contradicts the continuity of the function $f$.

□

**Theorem 2.** *A one-dimensional continuous RNN $h_{t+1} = f(h_t; Input)$ with an arbitrarily nonlinear readout cannot accurately represent a two-dimensional dynamical system without significant precision loss.*

*Proof.* Given the unconstrained nature of the readout, we assume that the one-dimensional RNN can learn an extremely nonlinear representation of two-dimensional state-space of the RL model. Generally, according to Netto's theorem, continuous bijections between smooth manifolds preserve the dimension[8], meaning that there does not exist a continuous bijective mapping between two smooth manifolds of different dimensions (e.g., one-dimensional state space and a two-dimensional state space). However, by relaxing the conditions, we can consider space-filling curves, which are surjective continuous functions from one-dimensional spaces to two-dimensional spaces (e.g., a Hilbert curve). True space-filling curves are non-differentiable everywhere, and therefore, cannot be learned by RNNs. Consequently, we consider approximations to space-filling curves through finite iterations (thus only non-differentiable at finite points). For an illustration of the Hilbert curve in two-dimensional state space, see its third iteration in Fig. S40b and four iteration in Fig. S40c. We define the 2D-to-1D approximation function $G$, such that each two-dimensional point is mapped to its nearest position on the Hilbert curve. This method allows the approximation of two-dimensional space by a one-dimensional curve to achieve higher accuracy in higher-order iterations.

Consider the red and blue points, which are close in the two-dimensional space and similarly mapped to proximate positions on the Hilbert curve ($G(red) \approx G(blue)$). After a single trial of [$A_L$ R=1], the red point transitions to the orange point $G(orange) = f(G(red))$ and the blue point transitions to the green point $G(green) = f(G(blue))$. According to the continuity of $f$, the representations of orange and blue points should be similar ($G(orange) \approx G(green)$). However, paradoxically, these points should occupy positions on the Hilbert curve that are far apart, as the orange and green points fall into different grid sections, thereby being mapped to disparate regions of the Hilbert curve ($G(orange)$ and $G(green)$). This indicates that, if the loss incurred from representing two-dimensional space by the one-dimensional curve ($G$) is minimized to be very small (e.g., through higher iterations), then the loss in accurately representing two-dimensional RL dynamics by one-dimensional RNN dynamics $f$ will conversely be significantly large.

□

The theorems above prove that two continuous variables cannot be packed into a single continuous variable without incurring significant loss. By extension, a $d$-dimensional dynamical system with continuous state variables cannot be characterized by less than $d$ dynamical variables. The relevance of this dimensionality is evidence in the fact that each of the $d$ variables that characterize a dynamical system describe one function mapping $d$ variables in trial $t$ to $d$ variables in trial $t + 1$. According to one of the most classical textbooks on dynamical systems, the dimensionality of the system is one of its most relevant characteristics, and systems of equal dimensionality can be directly compared to one another[9]. Clearly, this dimensionality would lose its meaning if a dynamical system could be compressed into fewer dynamical variables without substantial lost.

#### 3 Supplementary Figures

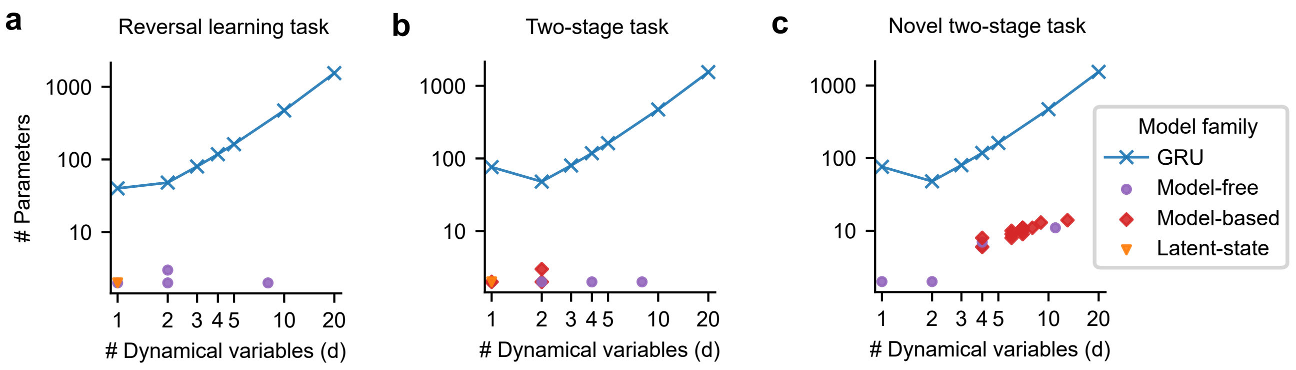


**Fig. S1. Number of trainable parameters in RNNs and cognitive models.** (a) Models in the reversal learning task. (b) Models in the two-stage task. (c) Models in the transition-reversal two-stage task. Here we reported the number of trainable parameters of the switching GRU models for $d = 1$ and that of the vanilla GRU models for $d \geq 2$, consistent with the main text (see Methods). Each model family may include multiple models, leading to two or more identical markers for a given $d$.

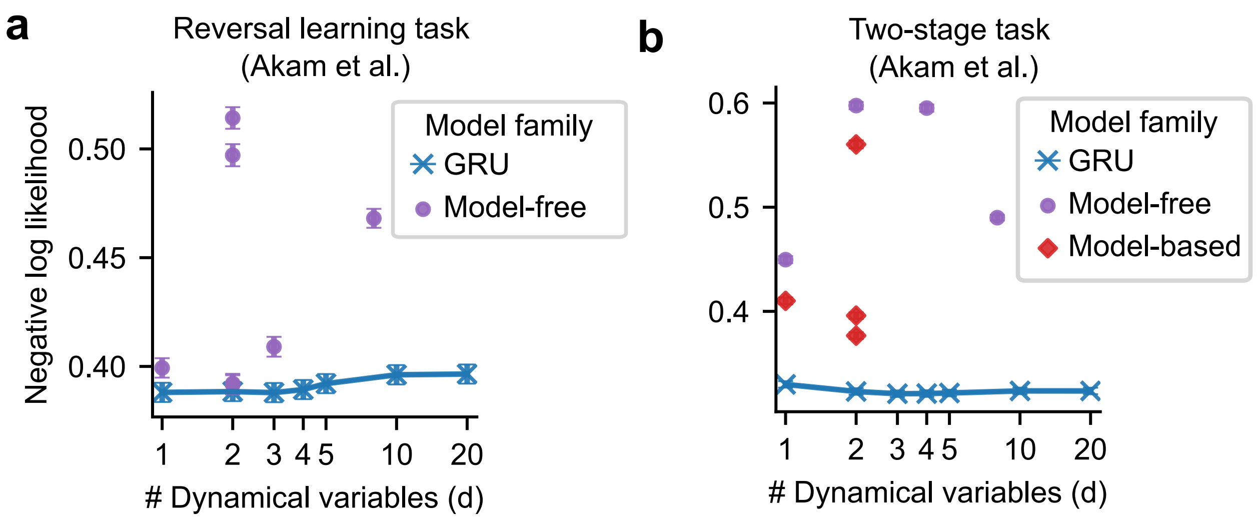


**Fig. S2. Tiny RNNs outperform classical cognitive models in predicting animals' choices in two additional datasets. (a-b)** Predictive performances (trial-averaged negative log-likelihood; lower is better) in two additional datasets. Performances are displayed as a function of the number of dynamical variables $d$, plotted on a log scale along the x-axis. The reported test performance for each model is the average over 10 outer folds in the nested cross-validation. Error bars show the average SEM (standard errors first calculated across outer and inner rounds, then averaged across individuals). Each dynamical variables in the GRU model family corresponds to one network unit; in cognitive models, dynamical variables can correspond to action values, state values, multi-trial choice perseveration, learned state-transition probabilities, among others. Each model family may include multiple model variants, leading to two or more identical markers for a given $d$. **(a)** In the reversal learning task with mice, the GRU models with $d = 1$ or $d = 2$ outperform all other models. **(b)** In the two-stage task with mice, the GRU model with $d = 2$ outperforms all other models.

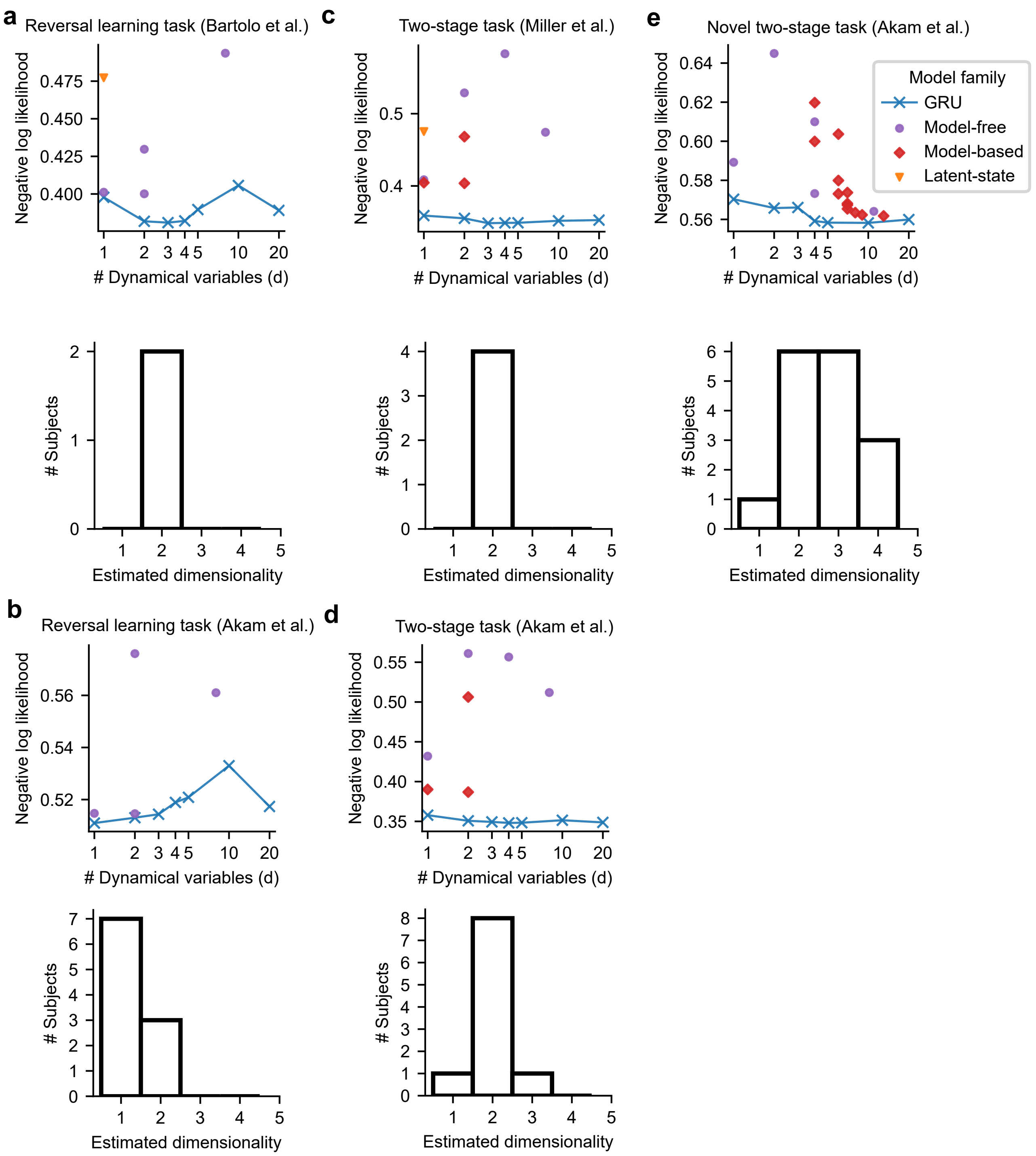
a
Reversal learning task (Bartolo et al.)
c
Two-stage task (Miller et al.)
e
Novel two-stage task (Akam et al.)
b
Reversal learning task (Akam et al.)
d
Two-stage task (Akam et al.)
Negative log likelihood
### Dynamical variables (d)
### Subjects
Estimated dimensionality
Model family
GRU
Model-free
Model-based
Latent-state

**Fig. S3. Tiny RNNs outperform classical cognitive models in predicting animals' choices at the individual level, providing estimated dimensionalities in three tasks.** (a-e) The predictive performance of models for one example animal (top) and the distribution of the number $d_*$ of dynamical variables for each animal (bottom). The estimated $d_*$ is determined such that an RNN model (GRU) with $d = d_*$ (but not $d > d_*$) dynamical variables significantly outperforms all RNN models with $d < d_*$ dynamical variables. (a) Monkeys (Bartolo et al.) performing the reversal learning task: for one example monkey, the GRU model with $d = 2$ outperforms all other models (top). Both monkeys have a dimensionality of $d_* = 2$ (bottom). (b) Mice (Akam et al.) performing the reversal learning task: for one example mouse, the GRU model with $d = 1$ outperforms all other models. Seven mice have a dimensionality of $d_* = 1$ and the remaining three have a dimensionality of $d_* = 2$ (bottom). (c) Rats (Miller et al.) performing the two-stage task: for one example rat, the GRU model with $d = 2$ outperforms all other models. Four rats have a dimensionality of $d_* = 2$ (bottom). (d) Mice (Akam et al.) performing the two-stage task: for one example mouse, the GRU model with $d = 2$ outperforms all other models. Eight of ten mice have a dimensionality of $d_* = 2$ (bottom). (e) Mice (Akam et al.) performing the transition-reversal two-stage task: for one example mouse, the GRU model with $d = 4$ outperforms all other models. Most mice have a dimensionality $d_*$ between 2 and 4 (bottom). Each model family may include multiple models, leading to two or more identical markers for a given $d$.

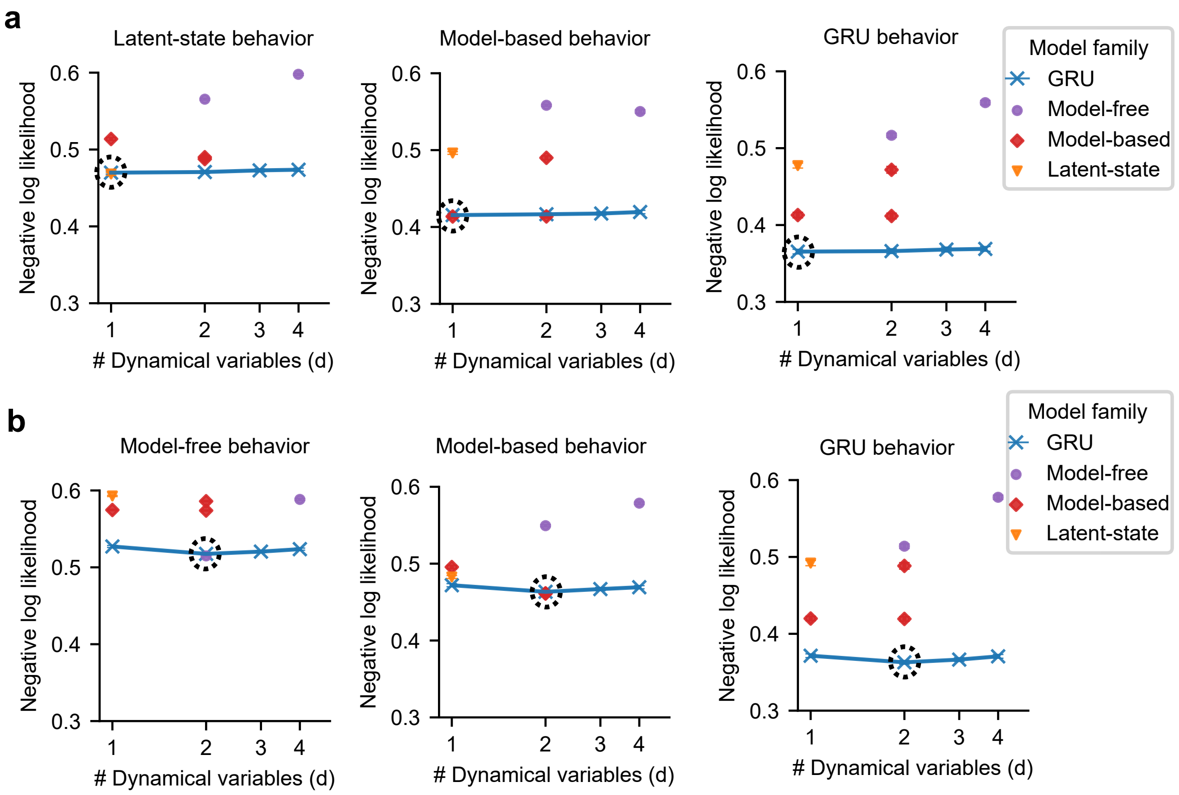


**Fig. S4. Tiny RNNs can predict choices as well as the ground-truth model in the two-stage task, measured by negative log-likelihood. (a-b)** The performance of each model in predicting the choices generated by a cognitive model in the two-stage task, with choice stochasticity arising due to sampling from the policy. Performances are displayed as a function of the number of dynamical variables $d$, plotted on a log scale along the x-axis. Each panel corresponds to choices generated by a different cognitive model. Dashed circles indicate the predictive performance of the ground-truth cognitive model — i.e., a model fitted to the choice data generated by the same model. Note that the RNNs have a similar predictive performance as the ground-truth model, suggesting that it can accurately identify the ground-truth strategy. **(a)** Simulated behavior generated by three models fitted to an example rat: a Bayesian inference (latent-state) agent with $d = 1$ (left), a model-based RL agent with $d = 1$ (middle), and a GRU agent with $d = 1$ (right). **(b)** Simulated behavior generated by three models fitted to an example rat: a model-free (Q(1)) agent with $d = 2$ (left), a model-based RL agent with $d = 2$ without value forgetting (middle), and a GRU agent with $d = 2$ (right). The reported test performance for each model is the average over 10 outer folds in the nested cross-validation. Error bars (almost invisible) show the average SEM (standard errors first calculated across outer and inner loops, then averaged across individuals). Each model family may include multiple model variants, leading to two or more identical markers for a given $d$.

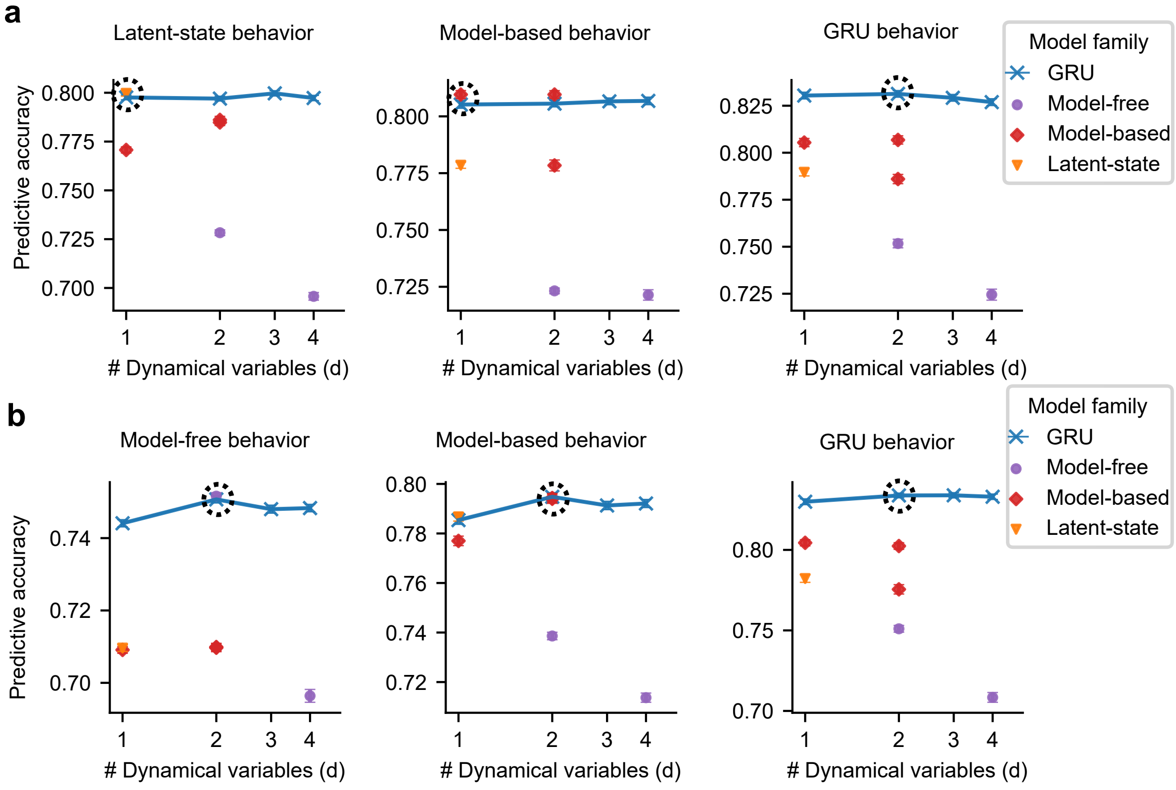


**Fig. S5. Tiny RNNs can predict choices as well as the ground-truth model in the two-stage task, measured by predictive accuracy. (a-b)** The performance of each model in predicting the choices generated by a cognitive model in the two-stage task, with choice stochasticity arising due to sampling from the policy. Performances are displayed as a function of the number of dynamical variables $d$, plotted on a log scale along the x-axis. Each panel corresponds to choices generated by a different cognitive model. Dashed circles indicate the predictive performance of the ground-truth cognitive model — i.e., a model fitted to the choice data generated by the same model. Note that the RNNs have a similar predictive performance as the ground-truth model, suggesting that it can accurately identify the ground-truth strategy. **(a)** Simulated behavior generated by three models fitted to an example rat: a Bayesian inference (latent-state) agent with $d = 1$ (left), a model-based RL agent with $d = 1$ (middle), and a GRU agent with $d = 1$ (right). **(b)** Simulated behavior generated by three models fitted to an example rat: a model-free (Q(1)) agent with $d = 2$ (left), a model-based RL agent with $d = 2$ without value forgetting (middle), and a GRU agent with $d = 2$ (right). The reported test performance for each model is the average over 10 outer folds in the nested cross-validation. Error bars (almost invisible) show the average SEM (standard errors first calculated across outer and inner loops, then averaged across individuals). Each model family may include multiple model variants, leading to two or more identical markers for a given $d$.

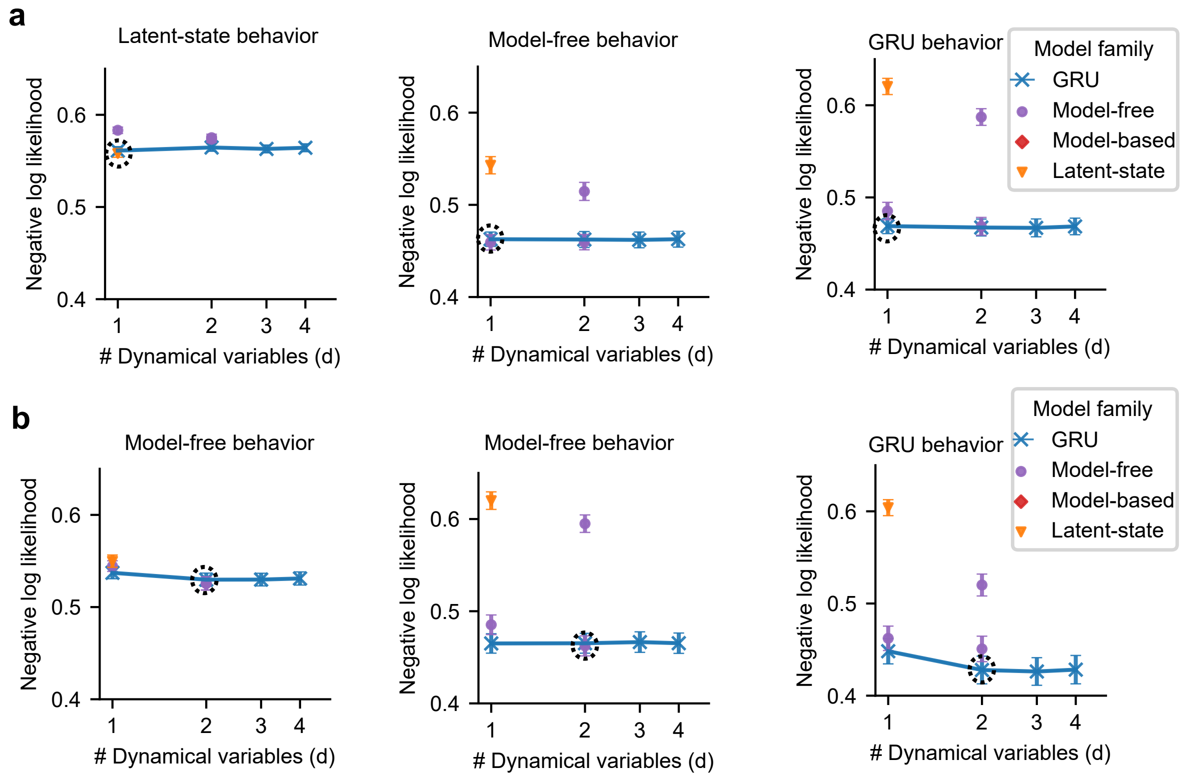


**Fig. S6. Tiny RNNs can predict choices as well as the ground-truth model in the reversal learning task, measured by negative log-likelihood. (a-b)** The performance of each model in predicting the choices generated by a cognitive model in the reversal learning task, with choice stochasticity arising due to sampling from the policy. Performances are displayed as a function of the number of dynamical variables $d$, plotted on a log scale along the x-axis. Each panel corresponds to choices generated by a different cognitive model. Dashed circles indicate the predictive performance of the ground-truth cognitive model — i.e., a model fitted to the choice data generated by the same model. Note that the RNNs have a similar predictive performance as the ground-truth model, suggesting that it can accurately identify the ground-truth strategy. **(a)** Simulated behavior generated by three models fitted to an example monkey: a Bayesian inference (latent-state) agent with $d = 1$ (left), a model-free RL agent with $d = 1$ (middle), and a GRU agent with $d = 1$ (right). **(b)** Simulated behavior generated by three models fitted to an example monkey: a model-free agent with $d = 2$ without value forgetting (left), a model-free RL agent with $d = 2$ with value forgetting (middle), and a GRU agent with $d = 2$ (right). The reported test performance for each model is the average over 10 outer folds in the nested cross-validation. Error bars show the average SEM (standard errors first calculated across outer loops, then averaged across individuals). Each model family may include multiple model variants, leading to two or more identical markers for a given $d$.

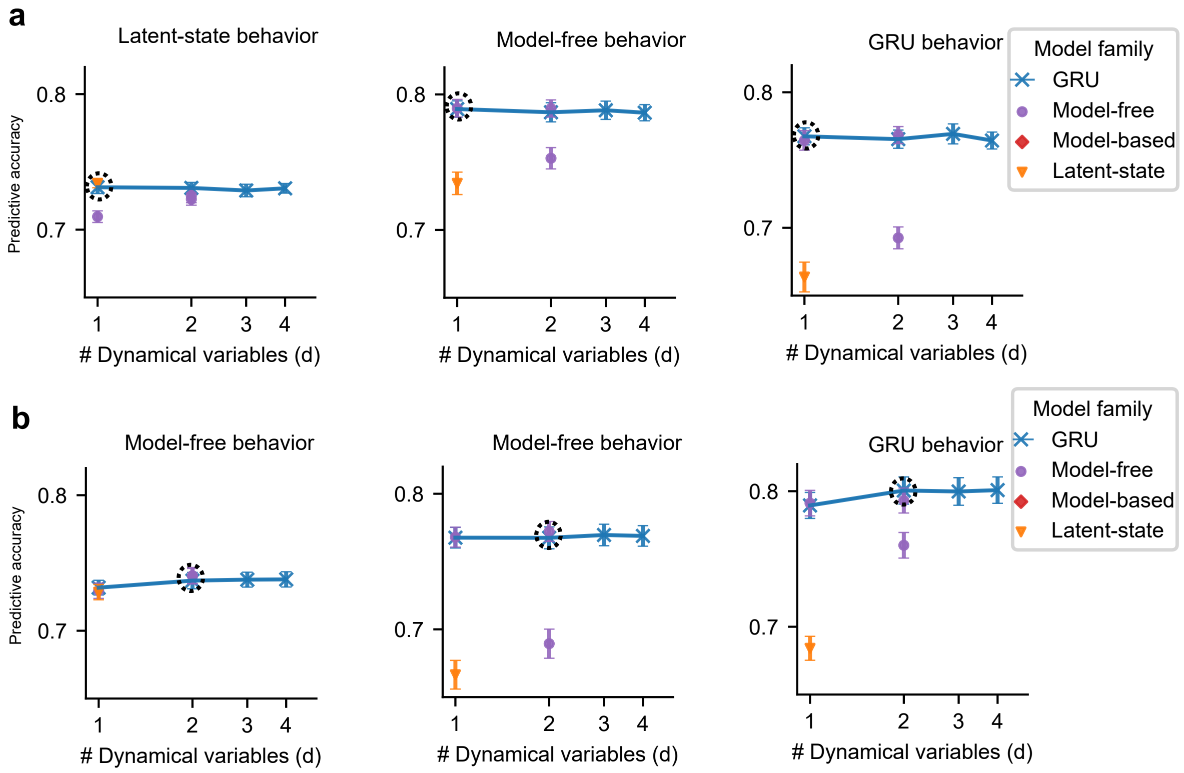


**Fig. S7. Tiny RNNs can predict choices as well as the ground-truth model in the reversal learning task, measured by predictive accuracy. (a-b)** The performance of each model in predicting the choices generated by a cognitive model in the reversal learning task, with choice stochasticity arising due to sampling from the policy. Performances are displayed as a function of the number of dynamical variables $d$, plotted on a log scale along the x-axis. Each panel corresponds to choices generated by a different cognitive model. Dashed circles indicate the predictive performance of the ground-truth cognitive model — i.e., a model fitted to the choice data generated by the same model. Note that the RNNs have a similar predictive performance as the ground-truth model, suggesting that it can accurately identify the ground-truth strategy. **(a)** Simulated behavior generated by three models fitted to an example monkey: a Bayesian inference (latent-state) agent with $d = 1$ (left), a model-free RL agent with $d = 1$ (middle), and a GRU agent with $d = 1$ (right). **(b)** Simulated behavior generated by three models fitted to an example monkey: a model-free agent with $d = 2$ without value forgetting (left), a model-free RL agent with $d = 2$ with value forgetting (middle), and a GRU agent with $d = 2$ (right). The reported test performance for each model is the average over 10 outer folds in the nested cross-validation. Error bars show the average SEM (standard errors first calculated across outer loops, then averaged across individuals). Each model family may include multiple model variants, leading to two or more identical markers for a given $d$.

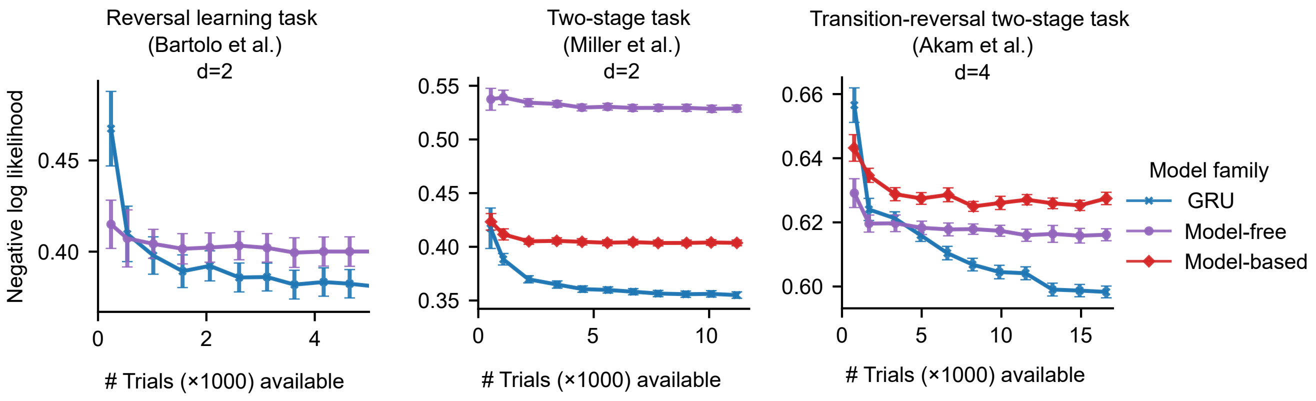


**Fig. S8. The predictive performance of tiny RNNs improves as the amount of available data increases.** Performance of each model (negative log-likelihood) when trained with varying amounts of data (number of trials for training and validation) to predict the choices of a representative animal in each dataset. Error bars show the average SEM (standard errors first calculated across outer and inner rounds, then averaged across individuals). The reported test performance for each model is the average over 10 outer folds in the nested cross-validation. **(Left)** Two-dimensional models (GRU: two units; model-free: two action values) applied to reversal learning task data from a representative monkey (Bartolo et al.). **(Middle)** Two-dimensional models (GRU: two units; model-free: two action values; model-based: two state values) applied to two-stage task data from a representative rat (Miller et al.). **(Right)** Four-dimensional models (GRU: four units; model-free: two action values and two state values; model-based: two state values and two state-transition probabilities) applied to transition-reversal two-stage task data from a representative mouse (Akam et al.).

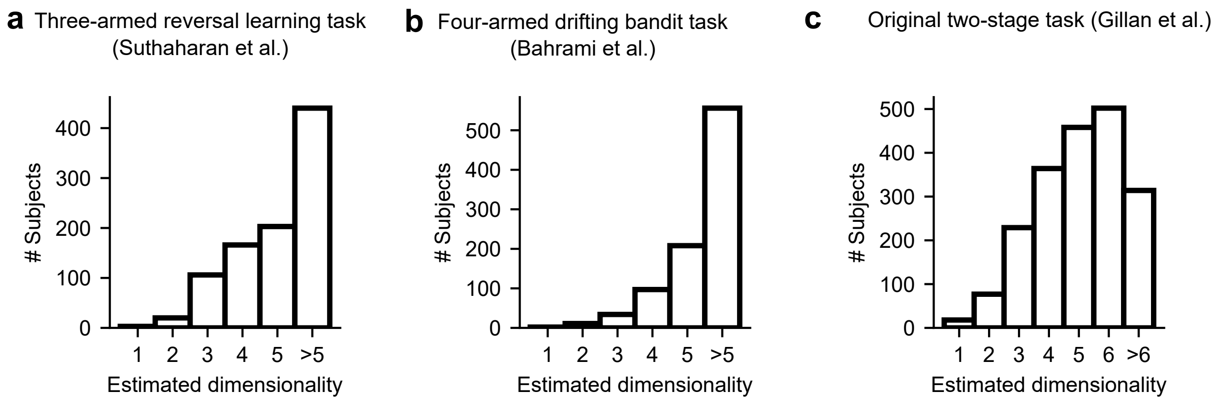


**Fig. S9. Estimated dimensionalities in three human reward learning tasks.** (a-c) The distribution of the number $d_*$ of dynamical variables for each subject. The estimated $d_*$ is determined such that an RNN model (GRU) with $d = d_*$ (but not $d > d_*$) dynamical variables significantly outperforms all RNN models with $d < d_*$ dynamical variables. (a) Human subjects (Suthaharan et al.) performing the three-armed reversal learning task. (b) Human subjects (Bahrami et al.) performing the four-armed drifting bandit task. (c) Human subjects (Gillan et al.) performing the original two-stage task.

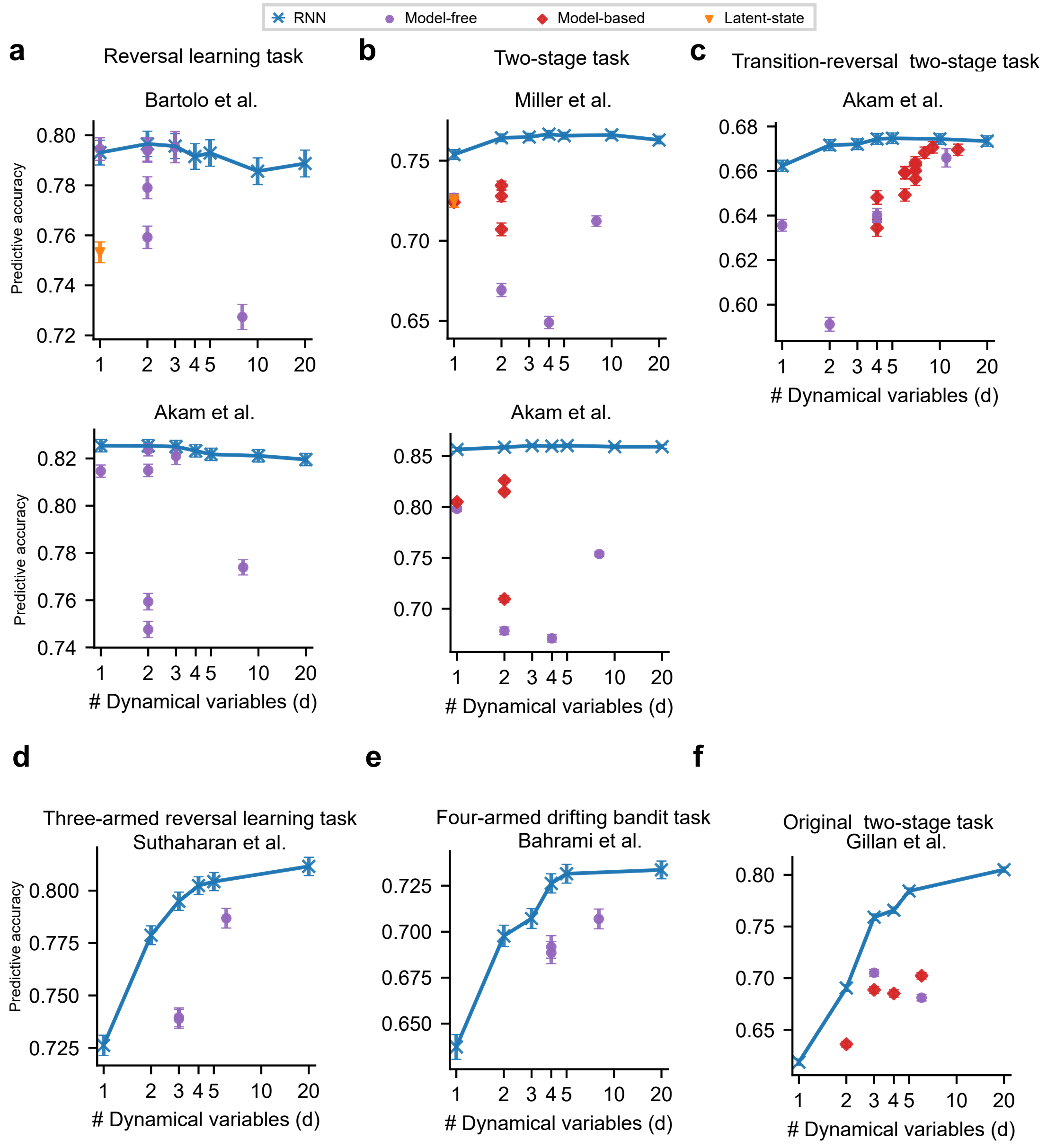


**Fig. S10. Tiny RNNs outperform classical cognitive models in predicting animals' and humans' choices at the group level, measured by predictive test accuracy.** Performances are displayed as a function of the number of dynamical variables *d*, plotted on a log scale along the x-axis. (a-c) The reported test accuracy for each model is the average over 10 outer folds in the nested cross-validation and then over individuals. Error bars show the average SEM (standard errors first calculated across outer and inner rounds, then averaged across individuals). (a) (Top) Monkeys (Bartolo et al.) performing the reversal learning task. (Bottom) Mice (Akam et al.) performing the reversal learning task. (b) (Top) Rats (Miller et al.) performing the two-stage task. (Bottom) Mice (Akam et al.) performing the two-stage task. (c) Mice (Akam et al.) performing the transition-reversal two-stage task. (d-f) The reported test accuracy for each model is the average over unseen test trials (using the interspersed split protocol) and then over individuals. Error bars show the SEM across individuals. (d) Humans (Suthaharan et al.) performing the three-armed reversal learning task. (e) Humans (Bahrami et al.) performing the four-armed drifting bandit task. (f) Humans (Gillan et al.) performing the original two-stage task. Each model family may include multiple models, leading to two or more identical markers for a given *d*.

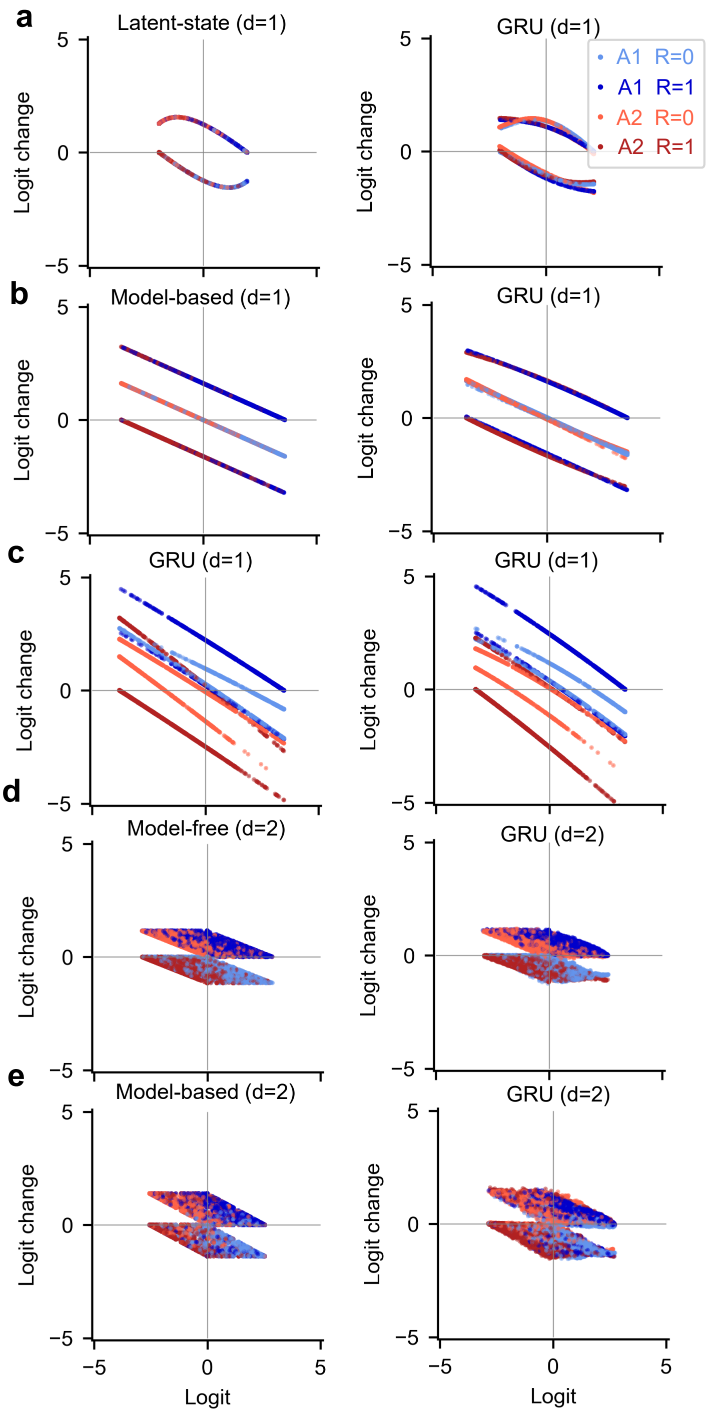


**Fig. S11. Tiny RNN models can recover the ground-truth phase portrait from simulated behavior generated by a cognitive model performing the two-stage task.** Ground-truth phase portrait (left) and RNN-recovered phase portrait (right). (a) A Bayesian inference (latent-state) model with $d = 1$. (b) A model-based model with $d = 1$. (c) A model-free (Q(1)) RL model with $d = 2$. (d) A model-based RL model (without value forgetting) with $d = 2$.

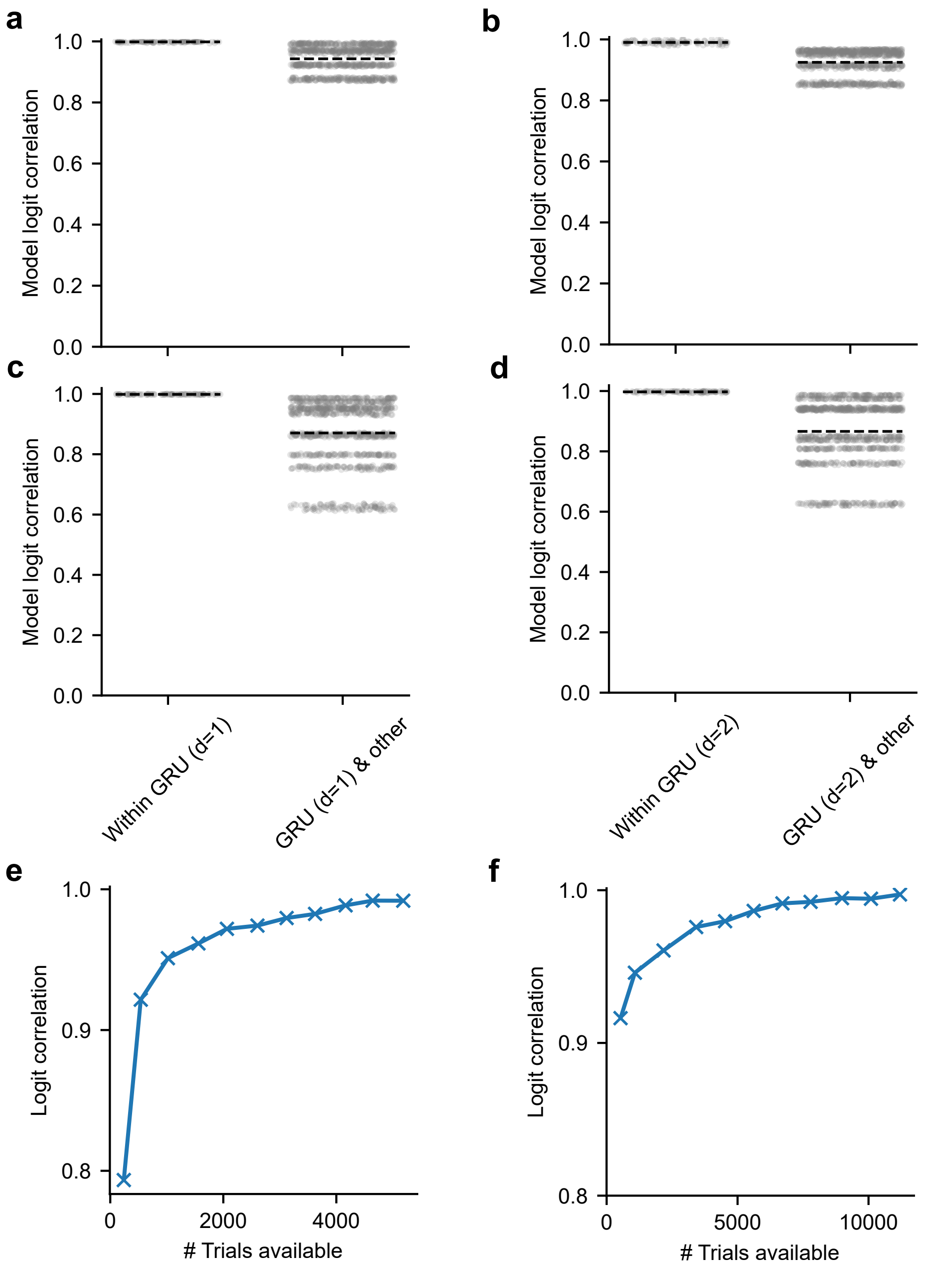


**Fig. S12. The strategies extracted by tiny RNN models (fitted to animal's behavior) are robust across outer rounds.** For each model, there are 10 model instances from the 10 outer rounds (in the nested cross-validation procedure) that are trained on different trials and with different hyperparameters. Each point is the correlation coefficient between the logits provided by two model instances. Dashed lines are the average across points. The correlations of logits generated by GRU models across outer rounds are close to one, significantly higher than the the correlations between logits generated by GRU models and by other cognitive models. (a) One-dimensional GRU model and (b) two-dimensional GRU model for one example monkey in the reversal learning task. (c) One-dimensional GRU model and (d) two-dimensional GRU model for one example rat in the two-stage task. (e) The logit correlation between pairs of GRU model instances (from outer rounds) improves as the amount of available data increases, suggesting convergence of extracted strategy (showing one example monkey in the reversal learning task). (f) The logit correlation between pairs of GRU model instances (from outer rounds) improves as the amount of available data increases, suggesting convergence of extracted strategy (showing one example rat in the two-stage task).

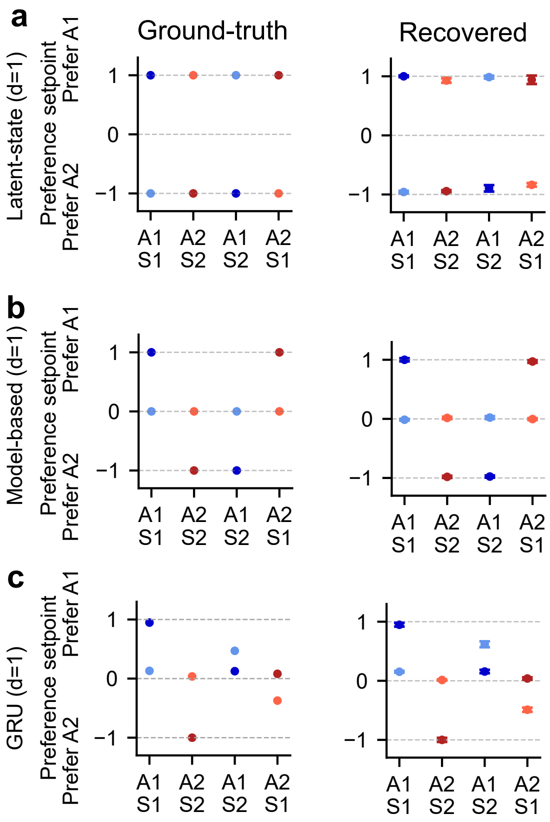


**Fig. S13. The strategies extracted by tiny RNN models (fitted to artificial behavior generated by one model) are robust across outer rounds.** For each model, there are 10 model instances from the 10 outer rounds (in the nested cross-validation procedure) that are trained on different trials and with different hyperparameters. Left column: Each point is the preference setpoint extracted from the ground-truth model. Right column: Each point is the preference setpoint recovered by the GRU model fitted to the behavior generated by the ground-truth model (averaged over 10 outer rounds). Error bars are the standard deviations across outer rounds. (a-c) One-dimensional models (with ground-truth models fitted to one example rat) in the two-stage task. (a) Bayesian inference (latent-state) strategy ($d = 1$). (b) model-based strategy ($d = 1$). (c) GRU strategy ($d = 1$).

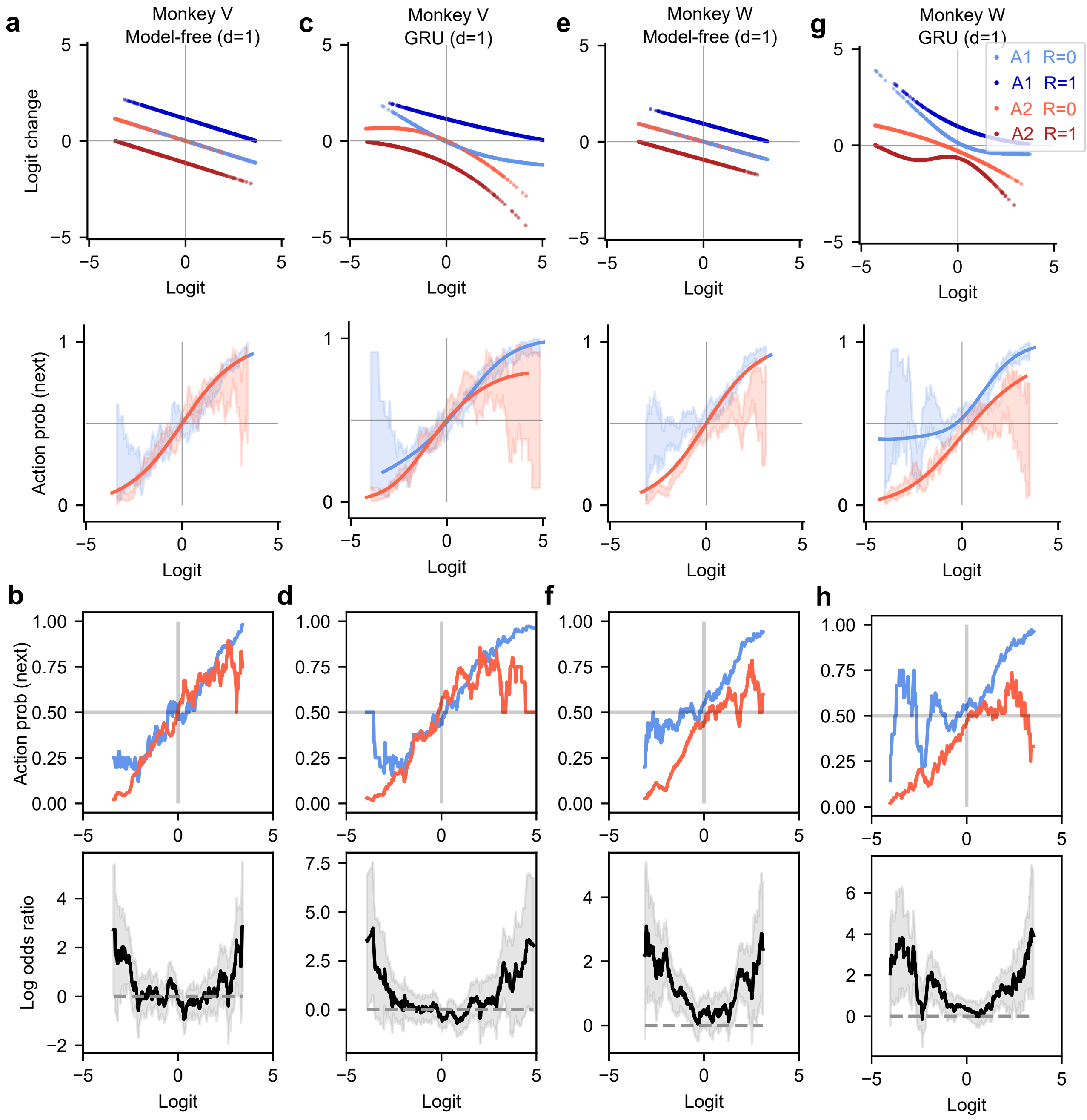

a
Monkey V
Model-free (d=1)
c
Monkey V
GRU (d=1)
e
Monkey W
Model-free (d=1)
g
Monkey W
GRU (d=1)
A1 R=0
A1 R=1
A2 R=0
A2 R=1
Logit change
Logit
Action prob (next)
b
d
f
h
Log odds ratio

**Fig. S14. Analysis of phase portraits of one-dimensional models after no reward.** (a) The phase portrait of the model-free RL model fitted to monkey V, showing logit change (top) or next-trial action probability (bottom) as a function of the current-trial logit and input. The next-trial action probability is a sigmoid transformation of the next-trial logit, which is the sum of the current-trial logit and logit change. The two curves for unrewarded $A_1$ (light blue) and unrewarded $A_2$ (light red) overlap completely, which is an assumption of the model-free model. The shaded regions are the 68% confidence interval of the proportion of selecting action $A_1$ (empirical action probability) in the next trial conditioning on the current-trial logit. (b) The proportion of selecting action $A_1$ (empirical action probability) in the next trial conditioning on the current-trial logit provided by the model-free model (top). This empirical action probability is higher in trials with $A_1$ followed by no reward (light blue) than with $A_2$ followed by no reward (light red), but only when the animal strongly prefers one action (i.e., the logit is away from zero). The difference between these two empirical action probability curves is measured by the log odds ratio of the two curves (bottom). The log odds ratio is positive when the light blue curve is above the light red curve and zero when the two curves overlap. The shaded regions are the 95% confidence interval (normal approximation) of the log odds ratio. The model-free RL model predicts zero log-odds ratio for all logits, but a U-effect in the log odds ratio is found. (c) The phase portrait of the GRU model fitted to monkey V. The GRU model discovered the non-overlapping features of the two curves representing state changes after no reward (i.e., light blue and light red). (d) The proportion of selecting action $A_1$ (empirical action probability) in the next trial conditioning on the current-trial logit provided by the GRU model (top). The non-overlapping effect aligns with the U-effect observed in the log odds ratio. (e-h) The phase portrait, empirical action probability, and log odds ratio of the model-free RL model fitted to monkey W. The model-free RL model's prediction about fully overlapping curves for no reward following $A_1$ (light blue) and $A_2$ (light red) deviates substantially from the empirical action probability marked by the shaded regions (e, bottom). In contrast, the GRU model's prediction fits well with the empirical action probability marked by the shaded regions (g, bottom). The U-effect in monkey W (f bottom and h bottom) is more pronounced than in monkey V (b bottom and d bottom).

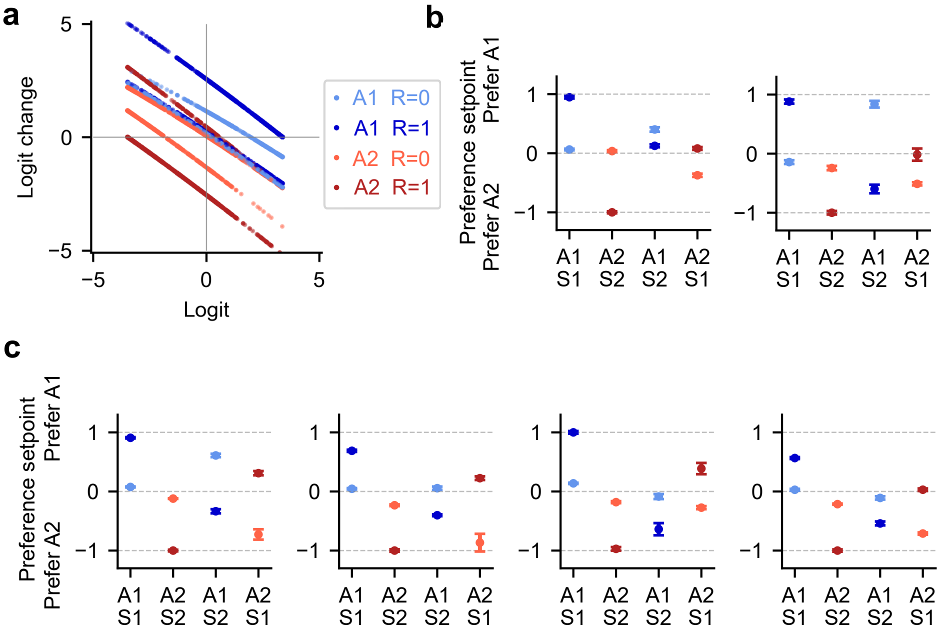


**Fig. S15. Phase portraits and preference setpoint patterns of the one-unit GRU model fitted to animals performing the two-stage task.** (a) The phase portrait of the GRU model fitted to one example rat (Miller et al.). Remarkably, there is no evidence of the strong prior as in the Bayesian inference (latent-state) model. Each color corresponds to two curves, one for each second-stage state ($S_1$ or $S_2$). The x-intercept $L_*$ of each curve corresponds directly to the preference setpoint on the left of (b). (b) The preference setpoint pattern on the left (one example rat, Miller et al.) is the same as in the main text and extracted from the phase portrait of (a). Another rat (Miller et al.) has a slightly different preference setpoint pattern for the rare transitions. The order of rewarded rare transition's and unrewarded rare transition's preference setpoint is opposite to that of the common transitions, suggesting the rat learned the main difference between them. (c) In addition to several mice (Akam et al.) with preference setpoint patterns similar to the left of (b), other mice have slightly different preference setpoint patterns (four examples shown). For all mice, the order of rewarded rare transition's and unrewarded rare transition's preference setpoint is opposite to that of the common transitions, suggesting they learned the main difference between common and rare transitions.

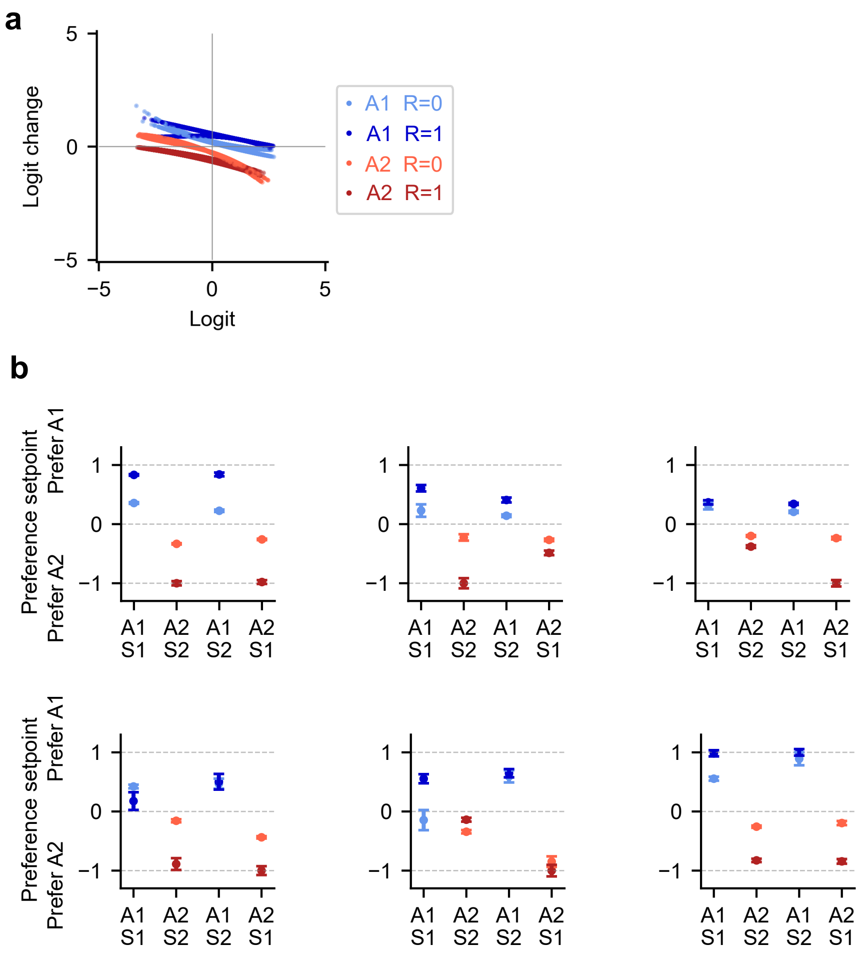


**Fig. S16. Phase portraits and preference setpoint patterns of the one-unit GRU model fitted to mice (Akam et al.) performing the transition-reversal two-stage task.** (a) The phase portrait of the GRU model fitted to an example mouse. Each color corresponds to two (almost overlapping) curves, one for each second-stage state ($S_1$ or $S_2$). The x-intercept $L_*$ of each curve matches the preference setpoint on the top left of (b). (b) The preference setpoint pattern on the top left is the same as in the main text and extracted from the phase portrait of (a). Some other mice have slightly different preference setpoint patterns (five examples shown here).

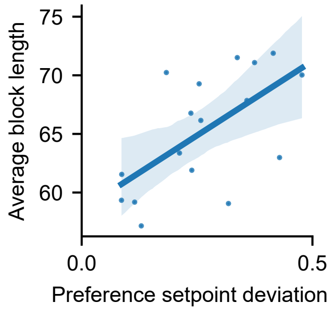


**Fig. S17. The reward-induced indifference effect correlates with task performance in the transition-reversal two-stage task.** A positive correlation ($r(15) = 0.62$, $p = 0.008$, 95% CI= $[0.198, 0.848]$) between an individual's tendency to gravitate towards indifference after a reward ("preference setpoint deviation") and the individual's average block length (reflecting task performance; lower is better). Each point is an animal. The shaded region is the 95% confidence interval of the linear regression.

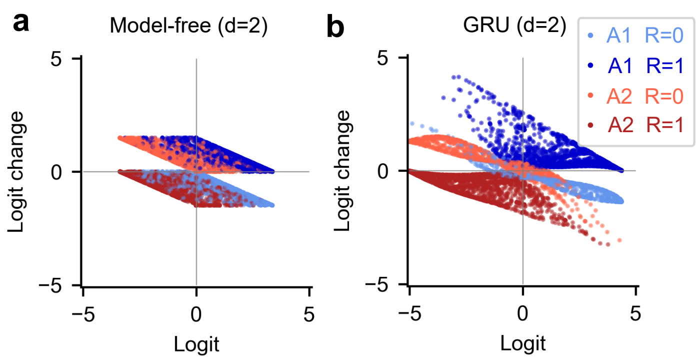


**Fig. S18. Plotting logit-change against logit for the two-dimensional models fitted to one monkey in the reversal learning task**, related to Fig. 4. (a) The two-dimensional model-free RL model without value forgetting. (b) The two-unit GRU model. For rewarded trials, the two-unit GRU model exhibits a pattern of logit-change that generally resembles that of the model-free RL model; for unrewarded trials, however, the two-unit GRU model exhibited a logit-change pattern similar to the one-unit GRU model but very different from the two-dimensional model-free RL model.

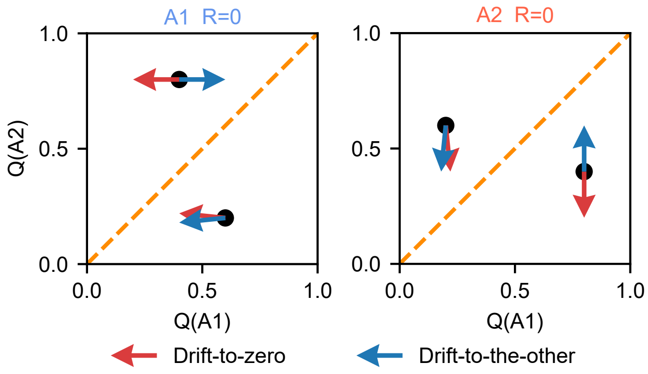


**Fig. S19. Schematic contrasting a drift-to-zero and a drift-to-the-other rule.**

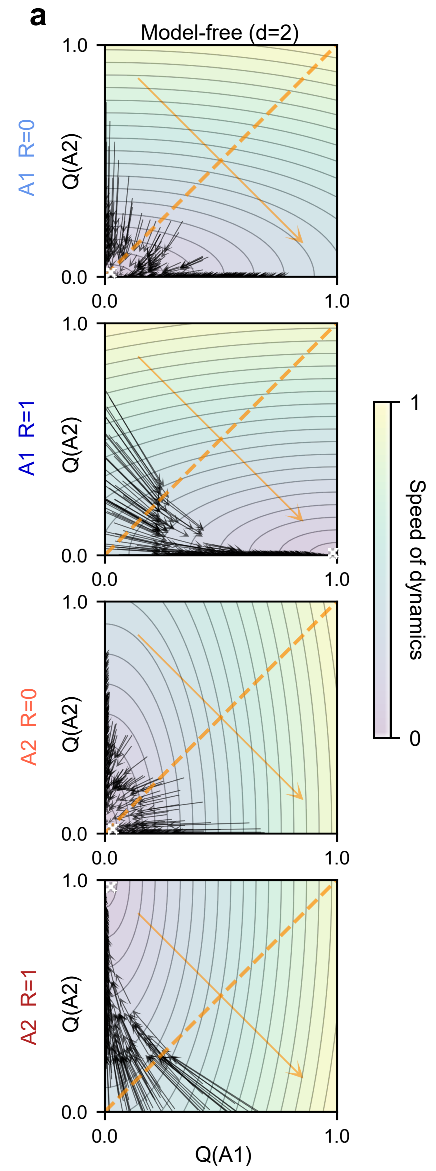


**Fig. S20. Vector fields for two-dimensional models fitted to one monkey in the reversal learning task**, compared to Fig. 4. Two-dimensional vector field analysis for the model-free RL model with value forgetting. Black arrows (flow lines) are the actual state trajectories experienced by the models. Orange dashed lines are the decision boundaries, and the orange arrows orthogonal to decision boundaries are the readout vectors. The crosses mark the point attractors or approximate line attractors. The two-dimensional state space is colored by the speed of the dynamics, defined by the norm of the velocity vector at each position (normalized by the maximal norm).

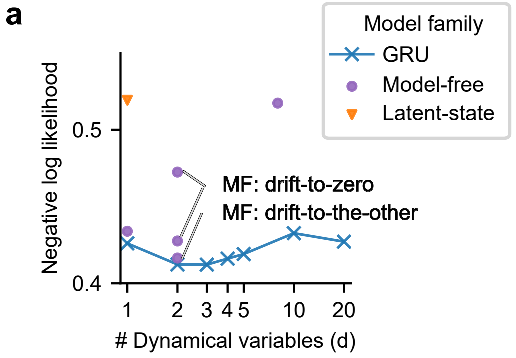


**Fig. S21. For monkeys (Bartolo et al.) performing the reversal learning task, the model-free RL model with the drift-to-other rule has predictive performance close to the two-dimensional GRU and better than the classical drift-to-zero rule.** This new model-free model is constructed from the phase portrait of the two-dimensional GRU. Each model family may include multiple models, leading to two or more identical markers for a given $d$.

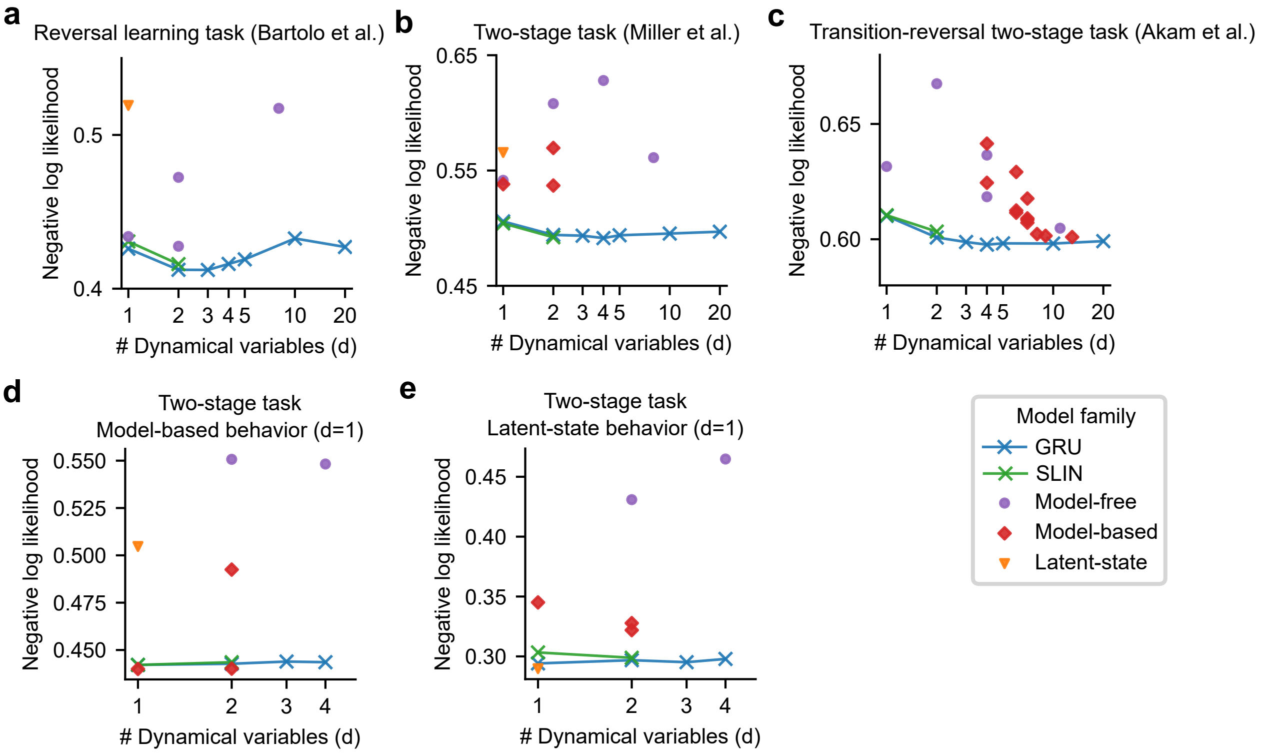


**Fig. S22. Predictive performance of models in three tasks, including switching linear neural networks (SLIN).** SLINs can achieve performance comparable to the GRUs with the same number of dynamical variables for some animals and artificial agents. (a) For monkeys (Bartolo et al.) performing the reversal learning task, the SLIN models with $d = 1$ (or 2) are slightly worse than the GRU models with the same $d$ (at both the group and individual levels). (b) For rats (Miller et al.) performing the two-stage task, the SLIN models with $d = 1$ (or 2) are comparable to the GRU models with the same $d$ (at both the group and individual levels). (c) For mice performing the transition-reversal two-stage task, the SLIN models with $d = 1$ are comparable to the GRU models with $d = 1$ (at both the group and individual levels). At the individual level, the SLIN models with $d = 2$ are comparable to the GRU models with $d = 2$ for some mice but worse for other mice. (d) When the behavior is generated by the model-free RL model with $d = 1$ performing the two-stage task, the SLIN models perform comparably to the best GRU models. (e) When the behavior is generated by the Bayesian inference (latent-state) model with $d = 1$ performing the two-stage task, the SLIN models are inferior to the GRU models with $d = 1$, due to the nonlinear learning rule in the Bayesian inference model. These results indicate that the SLIN models are less flexible than the GRU models and may struggle to capture nonlinear behavioral patterns. However, they can provide a more interpretable characterization of underlying strategies in cases where performance is comparable. Each model family may include multiple models, leading to two or more identical markers for a given $d$.

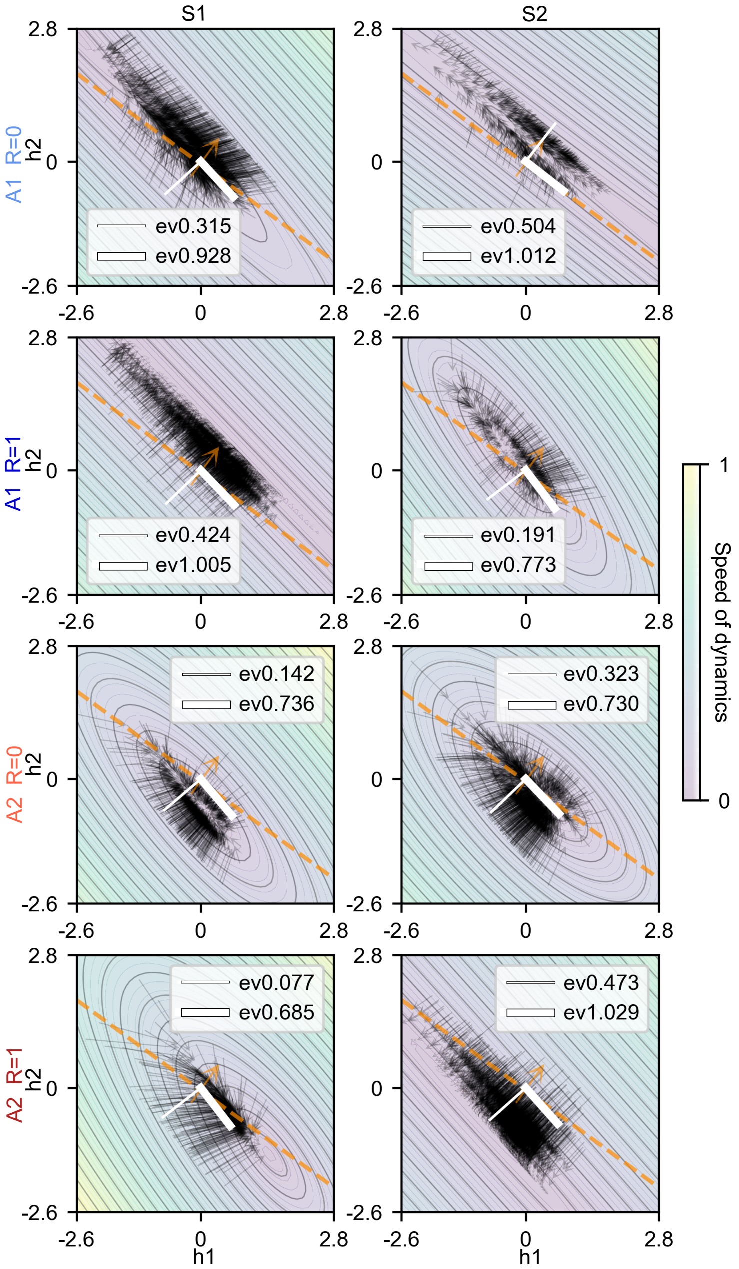


**Fig. S23. Vector fields for two-dimensional symmetric SLIN models fitted to one rat in the two-stage task.** Black arrows (flow lines) are the actual state trajectories experienced by the models. Orange dashed lines are the decision boundaries, and the orange arrows orthogonal to decision boundaries are the readout vectors. Two white lines (thinner for the smaller eigenvalue and thicker for the larger eigenvalue) are the eigenvectors of the recurrent weight matrix. The eigenvectors $v_2$ associated with the smaller $\lambda_2$ are always aligned with the readout vector. The two-dimensional state space is colored by the speed of the dynamics, defined by the norm of the velocity vector at each position (normalized by the maximal norm).

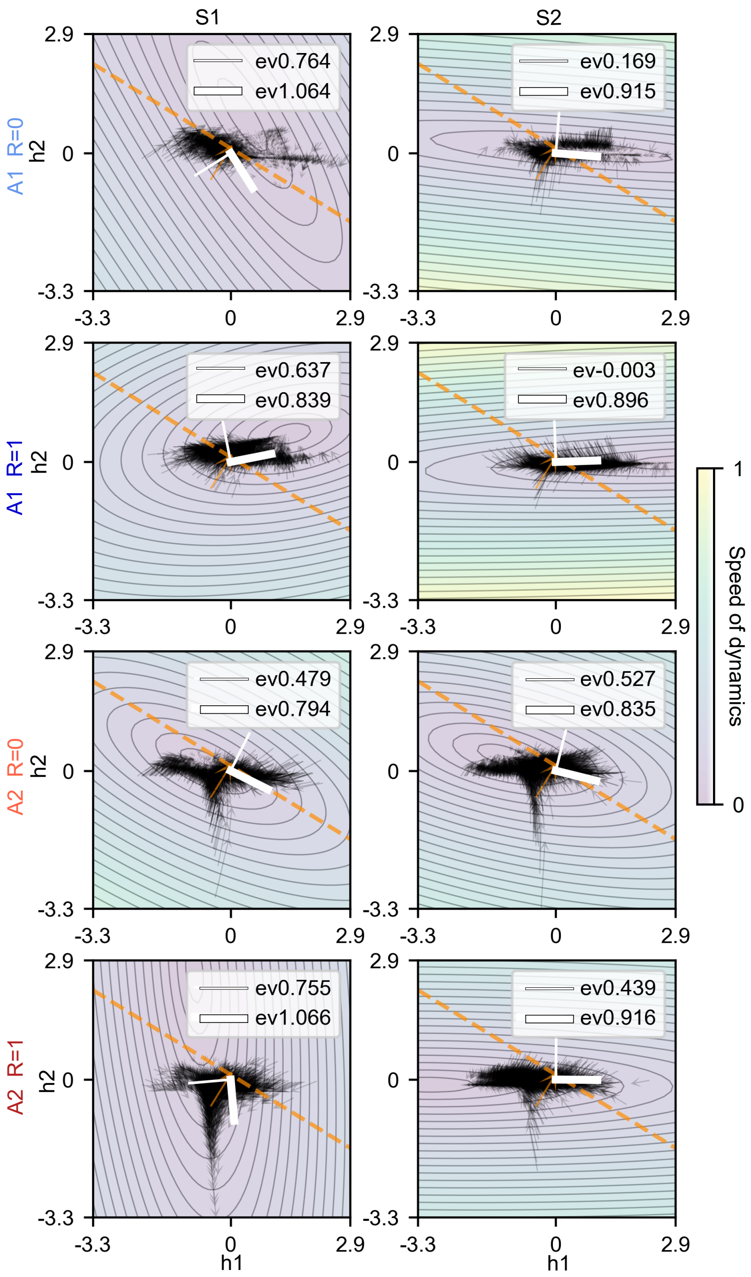


**Fig. S24. Vector fields for two-dimensional symmetric SLIN models fitted to one mouse in the transition-reversal two-stage task.** Black arrows (flow lines) are the actual state trajectories experienced by the models. Orange dashed lines are the decision boundaries, and the orange arrows orthogonal to decision boundaries are the readout vectors. Two white lines (thinner for the smaller eigenvalue and thicker for the larger eigenvalue) are the eigenvectors of the recurrent weight matrix. The two-dimensional state space is colored by the speed of the dynamics, defined by the norm of the velocity vector at each position (normalized by the maximal norm).

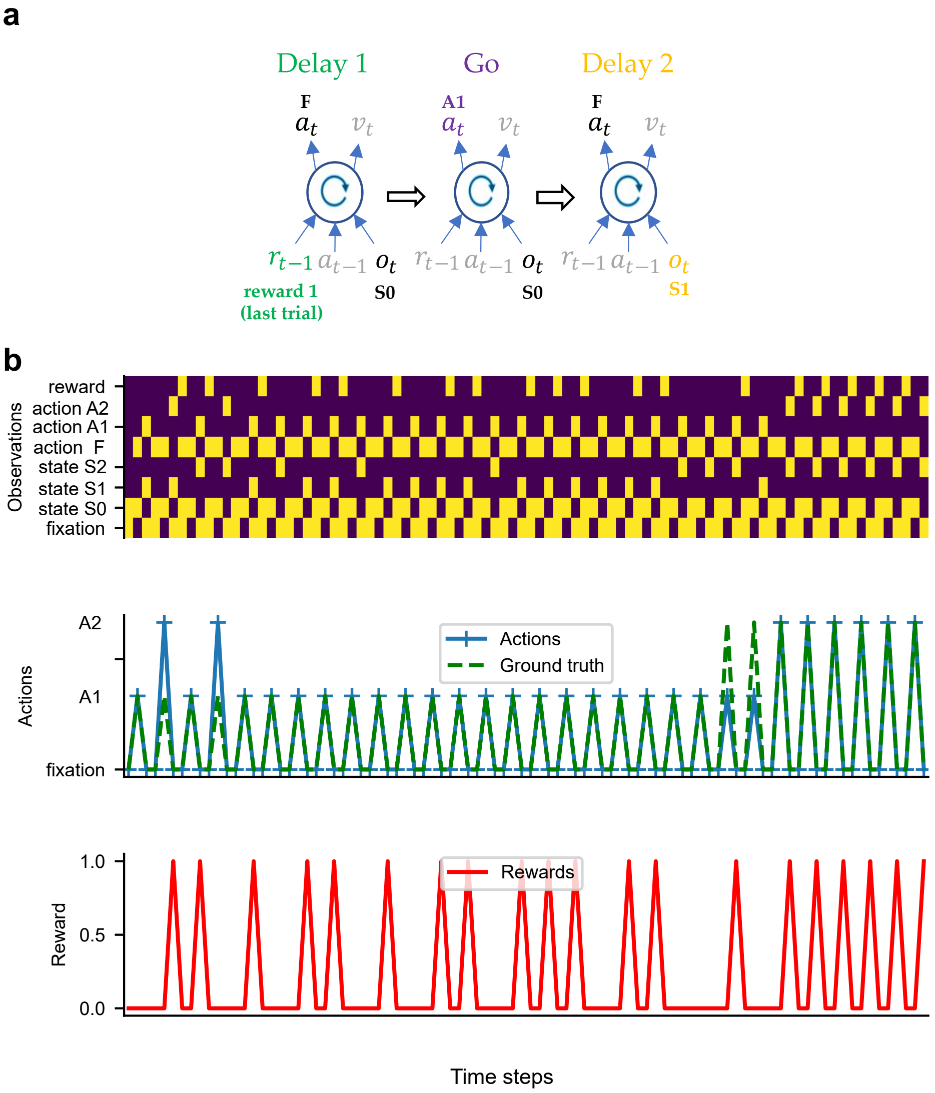


**Fig. S25. Training meta-RL agents on the two-stage task.** (a) Each trial consists of two stages. In the first-stage state $S_0$, the agent selects an action $A_1$ or $A_2$, which leads to a second-stage state $S_1$ or $S_2$ (common transition: $\Pr(S_1|A_1) = \Pr(S_2|A_2) = 0.8$, rare transition: $\Pr(S_2|A_1) = \Pr(S_1|A_2) = 0.2$). Each second-stage state leads to a different probability of a unit reward, with the most valuable state switching stochastically ($\Pr(R = 1|S_1) = 1 - \Pr(R = 1|S_2) = 0.8$ or 0.2, switched with a probability of 0.025 on each trial). There are three periods (discrete time steps) on one trial: Delay 1, Go, and Delay 2. During Delay 1, the agent receives the observation (state $S_0$ and a fixation signal) and the reward (1 or 0) from second-stage states on the last trial. During Go, the agent receives the observation of the state $S_0$ and a Go signal. During Delay 2, the agent receives the observation of state $S_1/S_2$ and a fixation signal. If the agent does not select action $A_1$ or $A_2$ during Go or select action F (Fixate) during Delay periods, a small negative reward (-0.1) is given. (b) An example block trajectory for a well-trained meta-RL agent, showing that the agent received inputs (top), took actions (middle) and received rewards (bottom) at each time step.

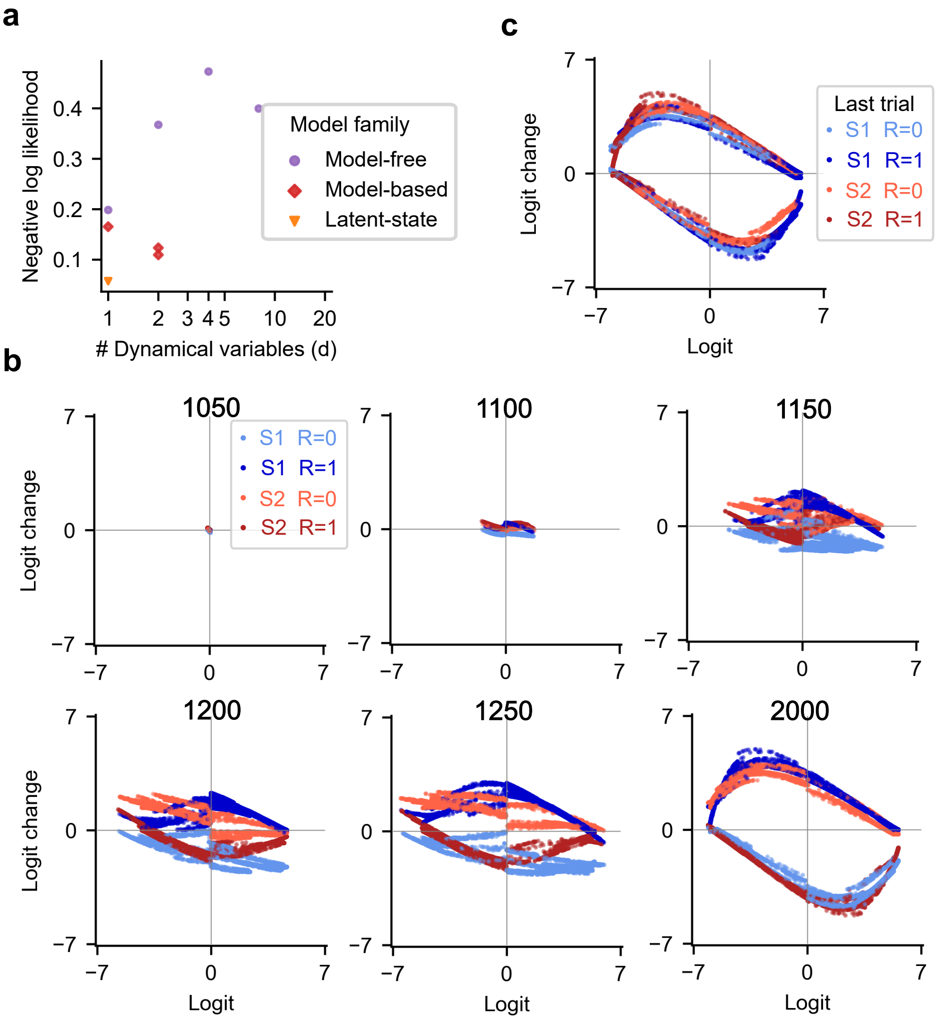


**Fig. S26. Analyzing meta-RL agents on the two-stage task.** (a) Model fitting results for the well-trained meta-RL agent studied in the main text. The Bayesian inference (latent-state) model explained the agent's behavior best. Each model family may include multiple models, leading to two or more identical markers for a given d. (b) The logit analysis of this agent during training (six training time steps shown here; units: 1000 training trials). The last panel is the same as in the main text. This agent learned the Bayesian-inference-like representation directly, with the intermediate representations differing from all cognitive models. (c) For this agent, the logit-change patterns deviate slightly from the Bayesian inference model's prediction of an exact function of logit, lying on several adjacent curves of the logit for a given input condition. These adjacent curves can be distinguished directly by past information, reflecting a history effect (colored by last-trial second-stages and last-trial rewards).

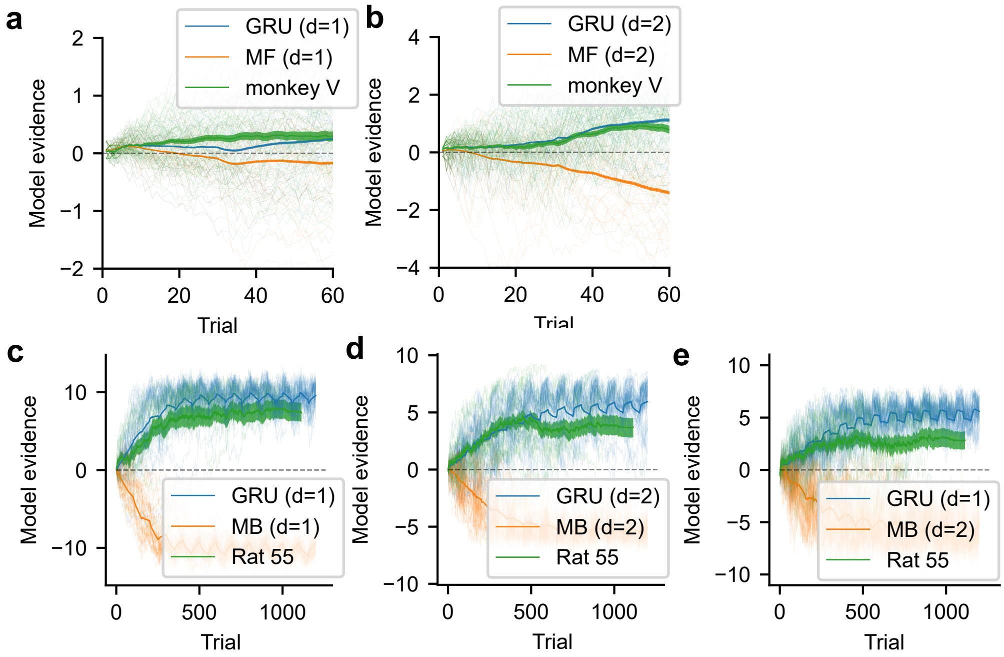


**Fig. S27. Model validation via behavior-feature identifier.** For a given task, we collect the behavioral sequences generated by models with a specific feature (positive class shown in blue) and by models without the feature (negative class shown in orange). We then train an RNN identifier on these sequences to predict their classes. This identifier is then applied on the actual behavioral sequences produced by subjects (shown in green). Each thin line represents a sequence of choices. Each thick line and corresponding shaded region is the average and standard error of model evidence across all sequences from a given model or the subject. (a) GRU (d=1) versus model-free strategy (d=1) fitted to one example monkey's behavior in the reversal learning task. (b) GRU (d=2) versus model-free strategy with value forgetting (d=2) fitted to one example monkey's behavior in the reversal learning task. (c) GRU (d=1) versus model-based strategy (d=1) fitted to one example rat's behavior in the two-stage task. (d) GRU (d=2) versus model-based mixture strategy (d=2) fitted to one example rat's behavior in the two-stage task. (e) GRU (d=1) versus model-based mixture strategy (d=2) fitted to one example rat's behavior in the two-stage task. These identifiers predicted that the actual behavioral sequences belong to the positive classes, thus successfully validating the strategies discovered by GRU models.

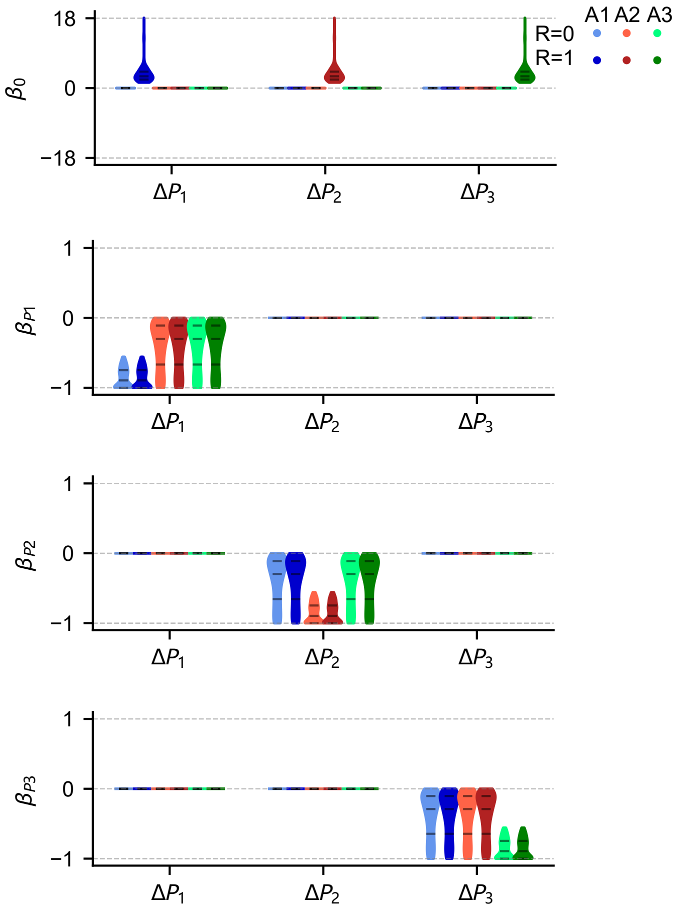


**Fig. S28.** Dynamical regression analysis for the three-dimensional model-free RL model with value forgetting fitted to subjects' behavior in the three-armed reversal learning task. $P_i$ and $\Delta P_i$ represents the preference and the preference change for action $A_i$ $(i = 1, 2, 3)$, respectively. We consider the regression $\Delta P_i \sim \beta_0^{(P_i)} + \beta_{P_1}^{(P_i)} P_1 + \beta_{P_2}^{(P_i)} P_2 + \beta_{P_3}^{(P_i)} P_3$ for each input condition. The linear regression coefficients are first calculated for each subject, and then group-level distributions of coefficients are shown as violins.

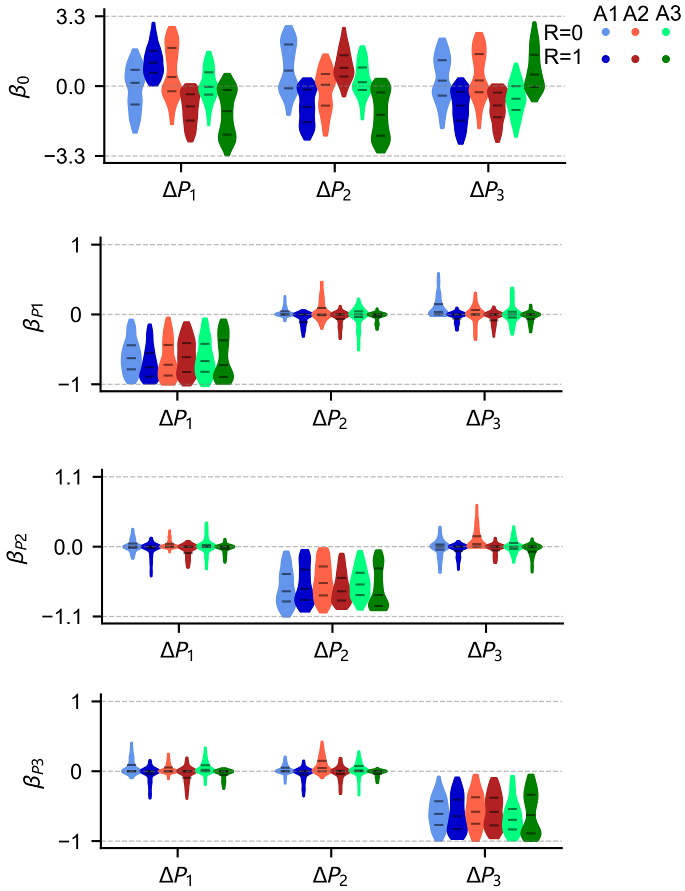


**Fig. S29.** Dynamical regression analysis for the three-dimensional GRU model fitted to subjects' behavior in the three-armed reversal learning task. $P_i$ and $\Delta P_i$ represents the preference and the preference change for action $A_i$ ($i = 1, 2, 3$), respectively. We consider the regression $\Delta P_i \sim \beta_0^{(P_i)} + \beta_{P_1}^{(P_i)} P_1 + \beta_{P_2}^{(P_i)} P_2 + \beta_{P_3}^{(P_i)} P_3$ for each input condition. The linear regression coefficients are first calculated for each subject, and then group-level distributions of coefficients are shown as violins.

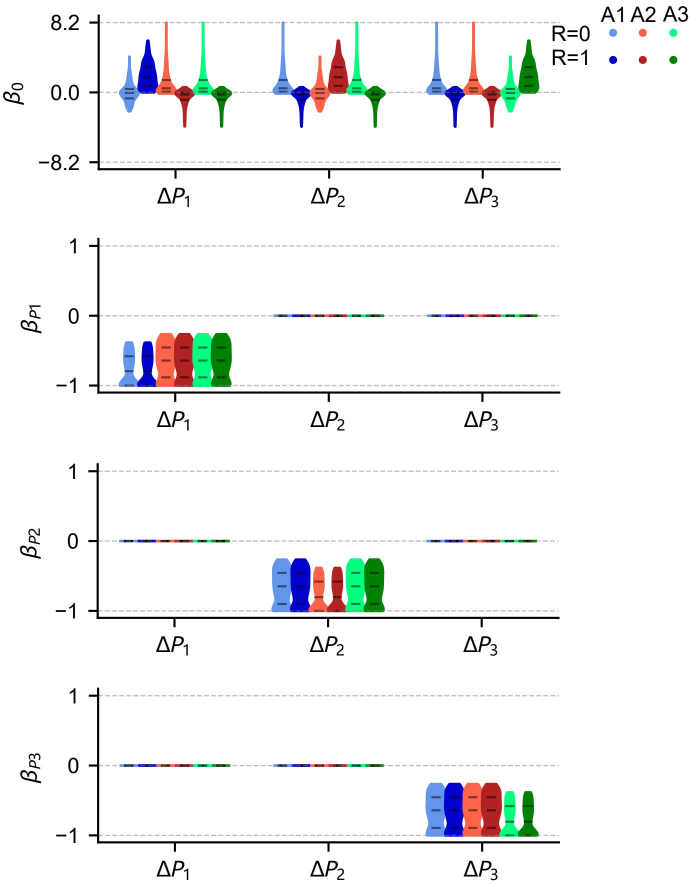


**Fig. S30.** Dynamical regression analysis for the three-dimensional model-free RL model with unchosen value updating and reward utility (inspired by the GRU model) fitted to subjects' behavior in the three-armed reversal learning task. $P_i$ and $\Delta P_i$ represents the preference and the preference change for action $A_i$ ($i = 1, 2, 3$), respectively. We consider the regression $\Delta P_i \sim \beta_0^{(P_i)} + \beta_{P_1}^{(P_i)} P_1 + \beta_{P_2}^{(P_i)} P_2 + \beta_{P_3}^{(P_i)} P_3$ for each input condition. The linear regression coefficients are first calculated for each subject, and then group-level distributions of coefficients are shown as violins.

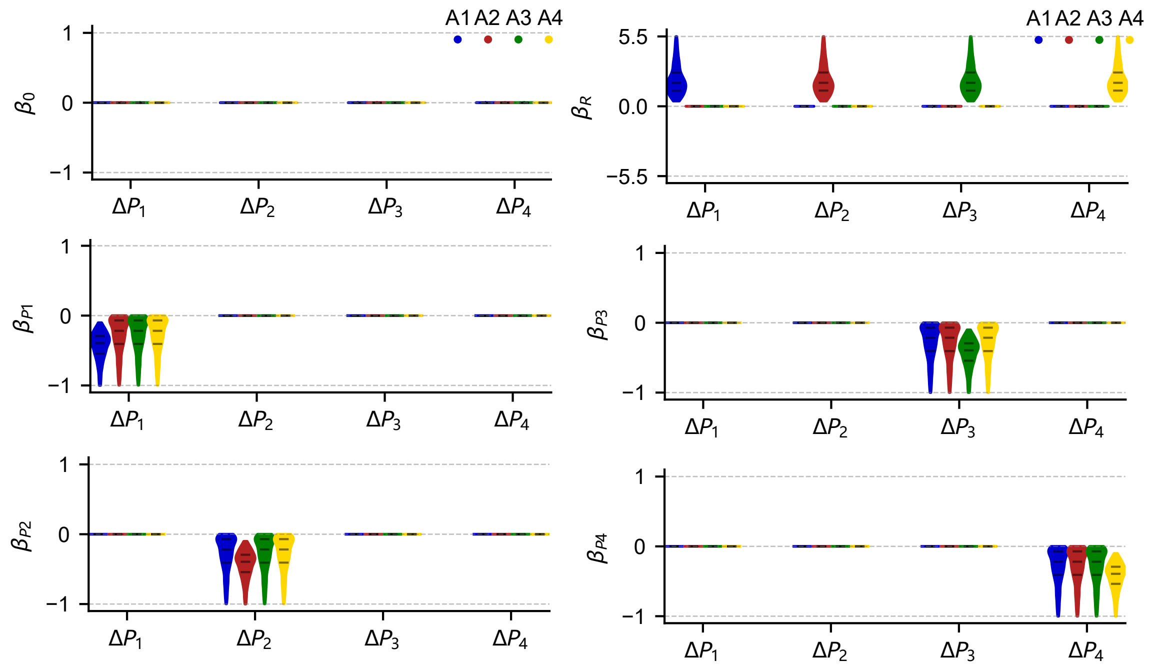


**Fig. S31.** Dynamical regression analysis for the four-dimensional model-free RL model with value forgetting fitted to subjects' behavior in the four-armed drifting bandit task. $P_i$ and $\Delta P_i$ represents the preference and the preference change for action $A_i$ ($i = 1, 2, 3, 4$), respectively. We consider the regression $\Delta P_i \sim \beta_0^{(P_i)} + \beta_R^{(P_i)} r + \beta_{P_1}^{(P_i)} P_1 + \beta_{P_2}^{(P_i)} P_2 + \beta_{P_3}^{(P_i)} P_3 + \beta_{P_4}^{(P_i)} P_4$ for each input condition. The linear regression coefficients are first calculated for each subject, and then group-level distributions of coefficients are shown as violins.

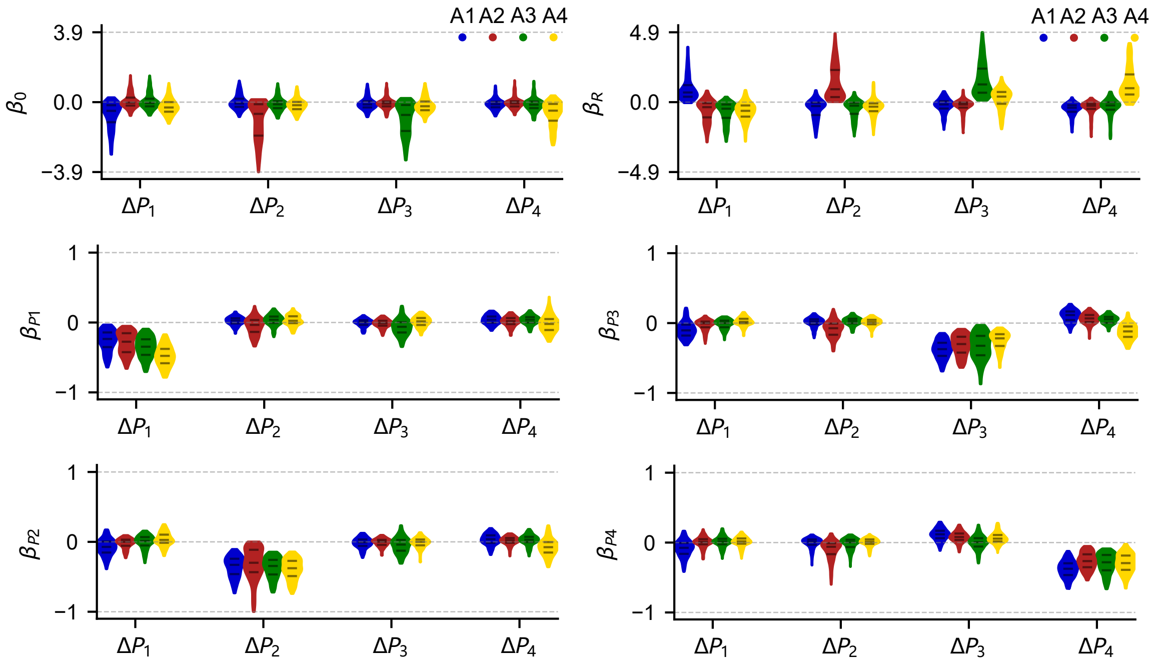


**Fig. S32.** Dynamical regression analysis for the four-dimensional GRU model fitted to subjects' behavior in the four-armed drifting bandit task. $P_i$ and $\Delta P_i$ represents the preference and the preference change for action $A_i$ ($i = 1, 2, 3, 4$), respectively. We consider the regression $\Delta P_i \sim \beta_0^{(P_i)} + \beta_R^{(P_i)} r + \beta_{P_1}^{(P_i)} P_1 + \beta_{P_2}^{(P_i)} P_2 + \beta_{P_3}^{(P_i)} P_3 + \beta_{P_4}^{(P_i)} P_4$ for each input condition. The linear regression coefficients are first calculated for each subject, and then group-level distributions of coefficients are shown as violins.

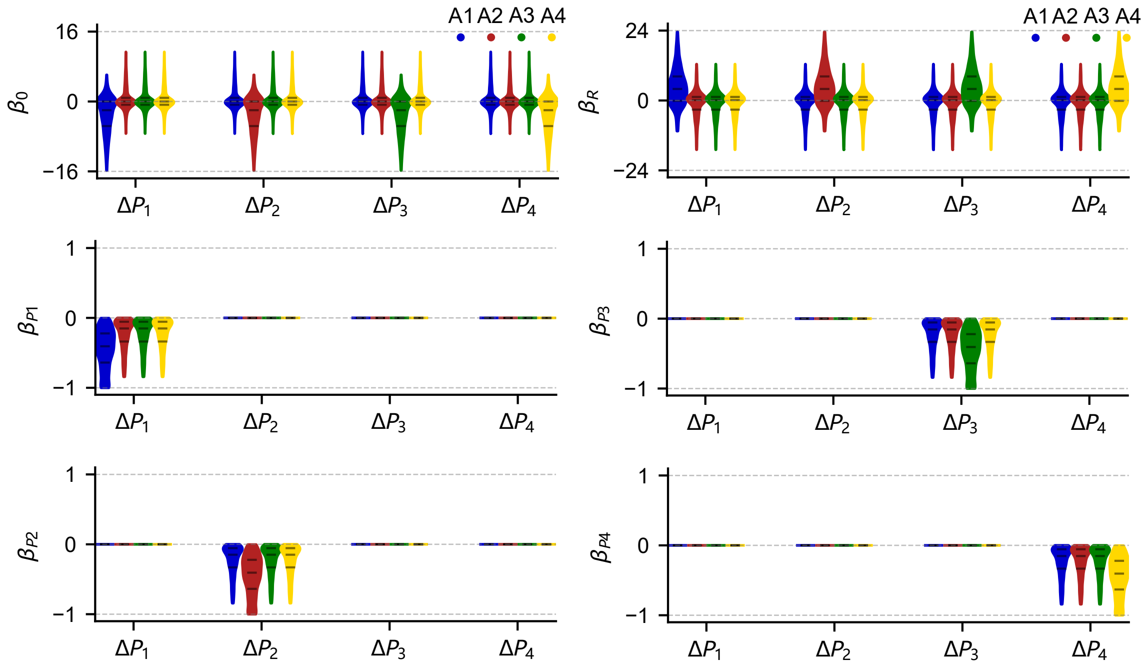


**Fig. S33.** Dynamical regression analysis for the four-dimensional model-free RL model with unchosen value updating and reward reference point (inspired by GRU) fitted to subjects' behavior in the four-armed drifting bandit task. $P_i$ and $\Delta P_i$ represents the preference and the preference change for action $A_i$ $(i = 1, 2, 3, 4)$, respectively. We consider the regression $\Delta P_i \sim \beta_0^{(P_i)} + \beta_R^{(P_i)} r + \beta_{P_1}^{(P_i)} P_1 + \beta_{P_2}^{(P_i)} P_2 + \beta_{P_3}^{(P_i)} P_3 + \beta_{P_4}^{(P_i)} P_4$ for each input condition. The linear regression coefficients are first calculated for each subject, and then group-level distributions of coefficients are shown as violins.

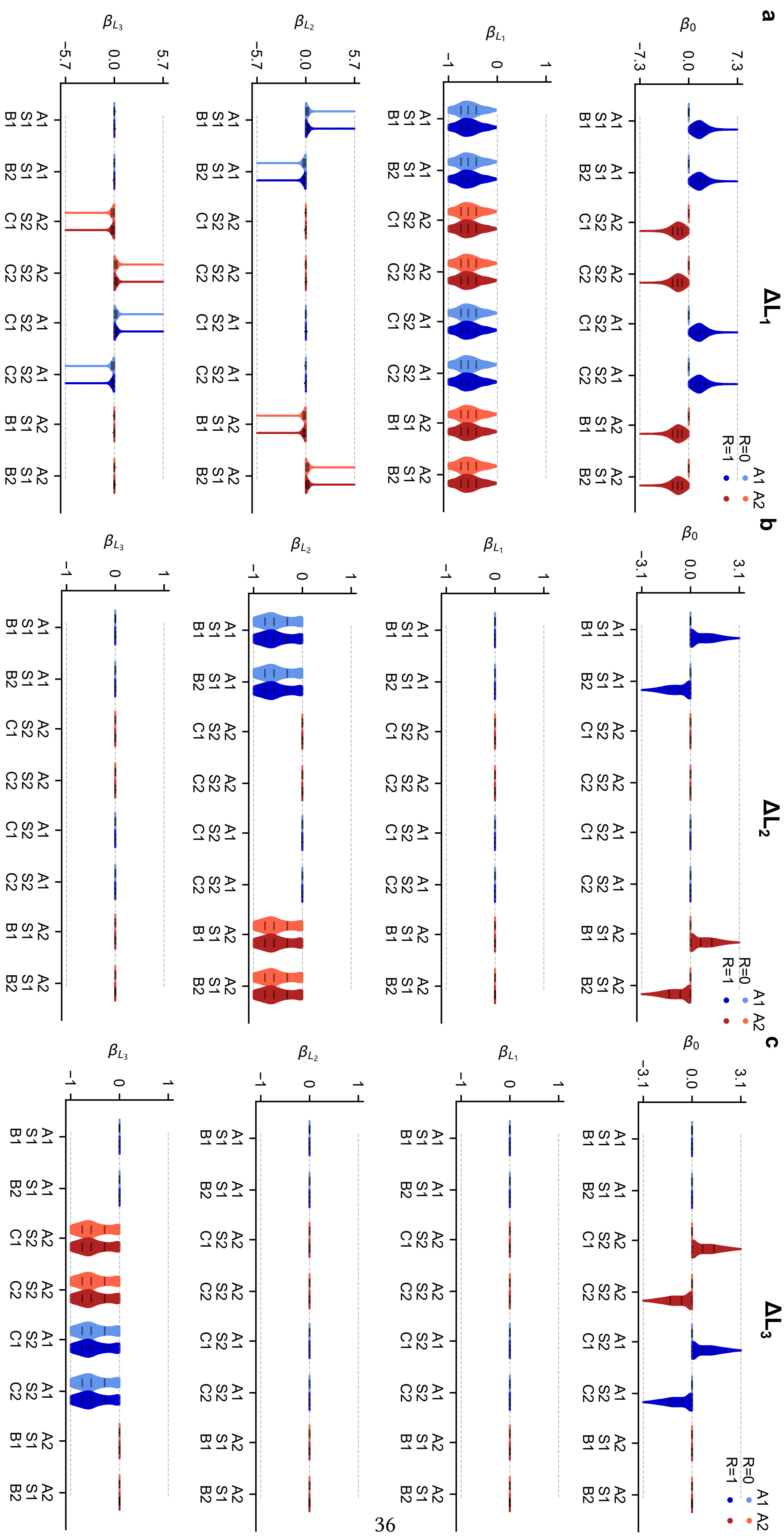
a
b
c
ΔL1
ΔL2
ΔL3
R=0
R=1
A1
A2

**Fig. S34.** Dynamical regression analysis for the three-dimensional model-free RL model fitted to subjects' behavior in the original two-stage task. $L_1$, $L_2$, $L_3$ are the logits for $A_1/A_2$ at the first-stage state, logits for $B_1/B_2$ at the second-stage state $S_1$ and logits for $C_1/C_2$ at the second-stage state $S_2$, respectively. we considered $\Delta L_i \sim \beta_0^{(L_i)} + \sum_{j=1}^{3} \beta_{L_j}^{(L_i)} L_j$ for each input condition. The linear regression coefficients are first calculated for each subject, and then group-level distributions of coefficients are shown as violins.

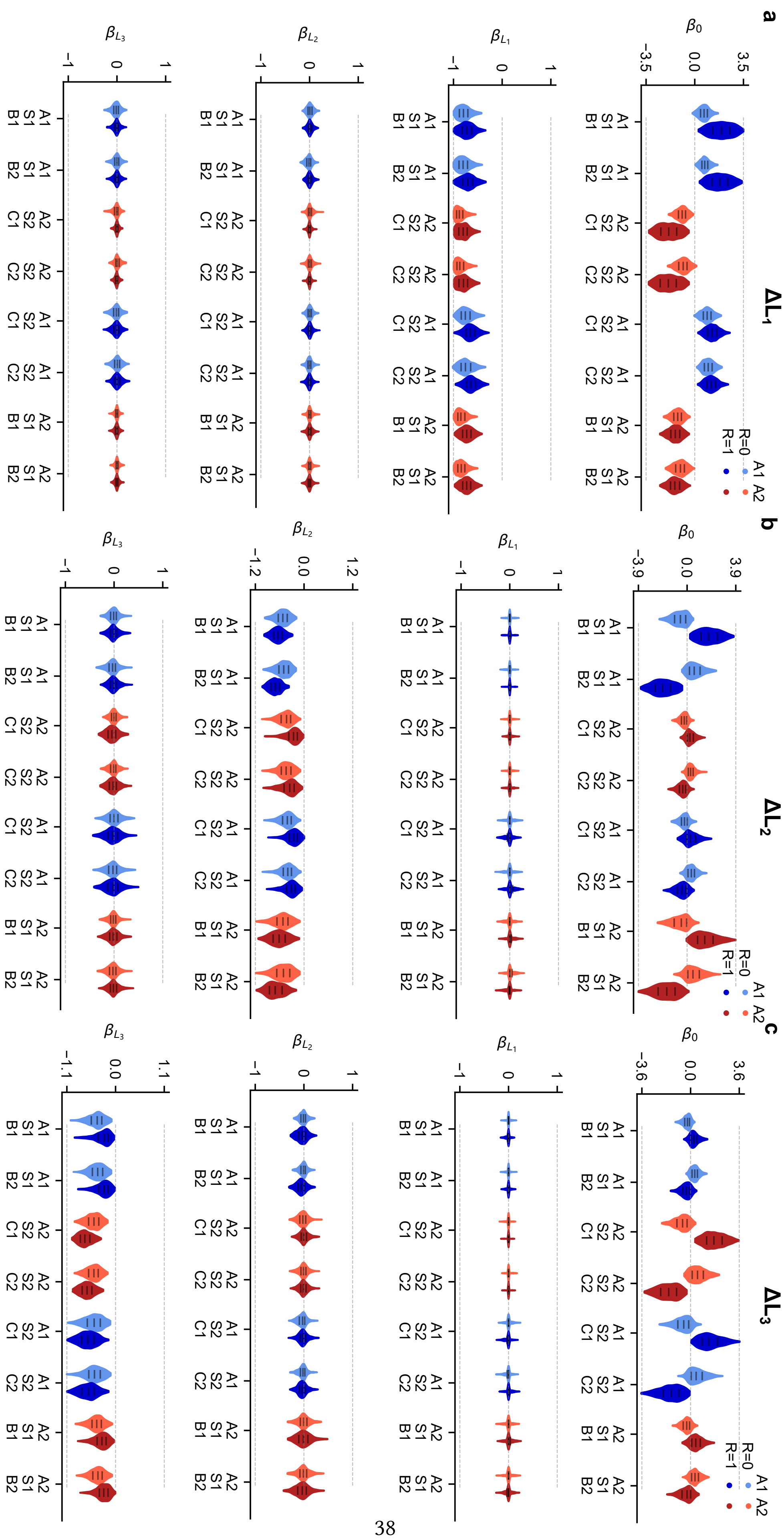
a
b
c
ΔL1
ΔL2
ΔL3
A1 A2
R=0
R=1

**Fig. S35.** Dynamical regression analysis for the three-dimensional GRU model fitted to subjects' behavior in the original two-stage task. $L_1$, $L_2$, $L_3$ are the logits for $A_1/A_2$ at the first-stage state, logits for $B_1/B_2$ at the second-stage state $S_1$ and logits for $C_1/C_2$ at the second-stage state $S_2$, respectively. we considered $\Delta L_i \sim \beta_0^{(L_i)} + \sum_{j=1}^{3} \beta_{L_j}^{(L_i)} L_j$ for each input condition. The linear regression coefficients are first calculated for each subject, and then group-level distributions of coefficients are shown as violins.

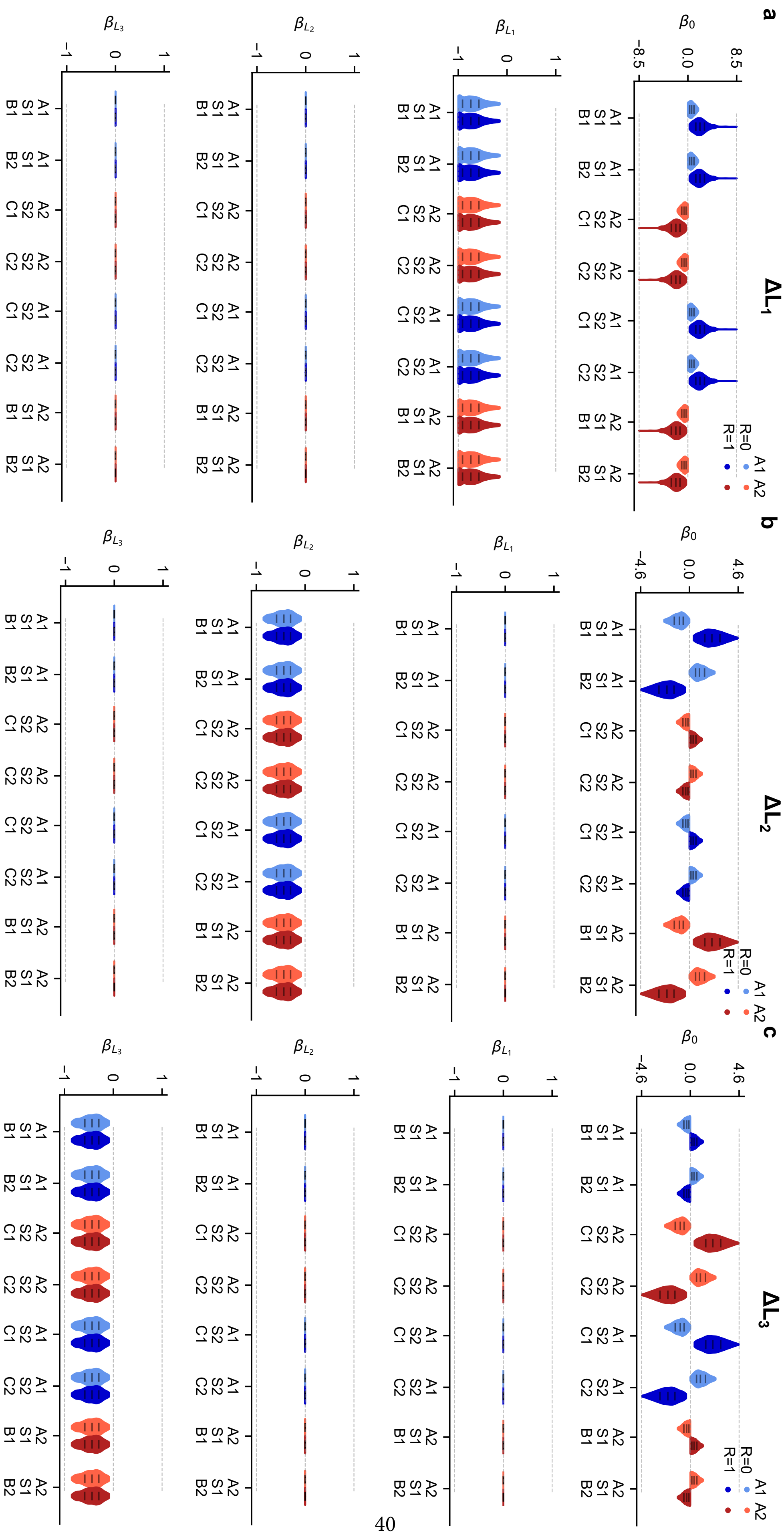
a
b
c
ΔL1
ΔL2
ΔL3
R=0
R=1
A1 A2
β0
βL1
βL2
βL3
A1 S1 B1
A1 S1 B2
A2 S2 C1
A2 S2 C2
A1 S2 C1
A1 S2 C2
A2 S1 B1
A2 S1 B2

**Fig. S36.** Dynamical regression analysis for the three-dimensional model-free RL model with reward utility (inspired by the GRU model) fitted to subjects' behavior in the original two-stage task. $L_1$, $L_2$, $L_3$ are the logits for $A_1/A_2$ at the first-stage state, logits for $B_1/B_2$ at the second-stage state $S_1$ and logits for $C_1/C_2$ at the second-stage state $S_2$, respectively. we considered $\Delta L_i \sim \beta_0^{(L_i)} + \sum_{j=1}^{3} \beta_{L_j}^{(L_i)} L_j$ for each input condition. The linear regression coefficients are first calculated for each subject, and then group-level distributions of coefficients are shown as violins.

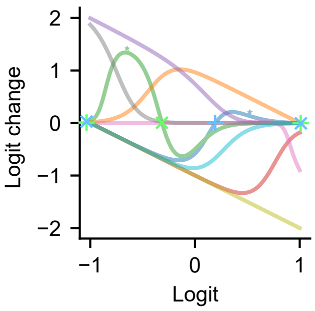


**Fig. S37. Possible phase portraits captured by one-unit GRU models.** In the GRU model, the effect of a constant input can be absorbed into the input biases (see Methods). We thus can determine its logit-change functions from the parameters (weights and biases) while ignoring the constant input. We randomly sampled a few sets of parameters (i.i.d. from a uniform distribution between -10 and 10) to generate phase-portrait curves (here the logits are normalized by the maximum). Empirically, we observed that these phase portraits have at most three fixed points where the logit change vanishes (e.g., green and blue curves and crosses; $\times$ crosses for stable fixed points, and $+$ crosses for unstable fixed points), indicating that the complexity of dynamics that can be captured is limited by the expressive power of one-unit GRUs.

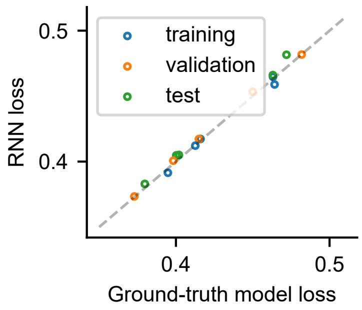


**Fig. S38. Evaluation of the interspersed split protocol used in human datasets.** To evaluate whether there exists any test data leak in this new protocol, we simulated artificial choice data using cognitive models in the two-stage task. We then train GRU models (with ten neurons) on the sequences generated by one cognitive model. Because the RNNs have the capacity to memorize trained sequences, if the test data is leaked into the training data, RNNs will demonstrate better losses on the test data than the ground-truth cognitive model. Empirically, we found that RNNs' test losses never go below the ground-truth model losses, ruling out the possibility of the data leak.

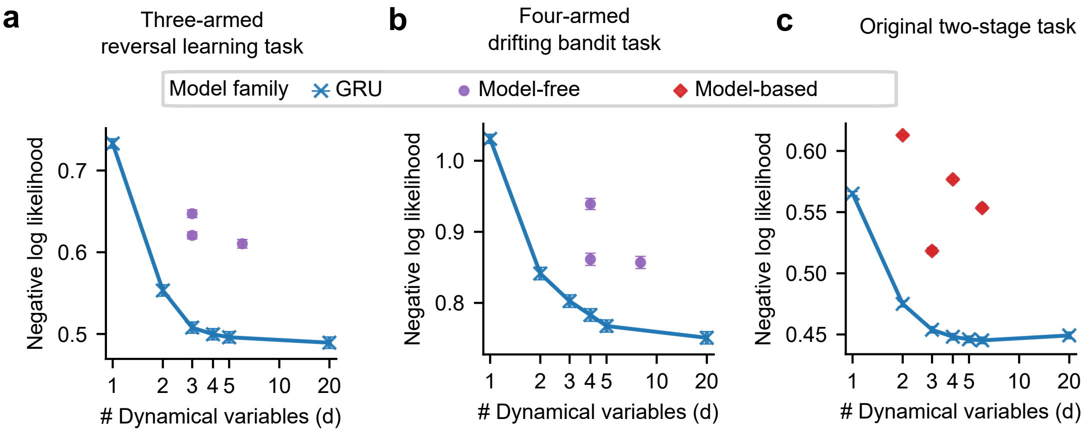


**Fig. S39. Application of the cross-subject split protocol to human datasets.** Predictive performances (trial-averaged negative log-likelihood; lower is better) in three tasks (one per panel). Performances are displayed as a function of the number of dynamical variables $d$, plotted on a log scale along the x-axis. The reported performance for each student RNN model and cognitive model is the average over all subjects on the unseen subjects' test trials using the cross-subject split protocol. Error bars show the SEM across individuals. Each dynamical variable in the GRU model family corresponds to one network unit; in cognitive models, dynamical variables can correspond to action values, state values, multi-trial choice perseveration, learned state-transition probabilities, among others. Each model family may include multiple model variants, leading to two or more identical markers for a given $d$. **(a)** Three-armed reversal learning task. **(b)** Four-armed drifting bandit task. **(c)** Original two-stage task.

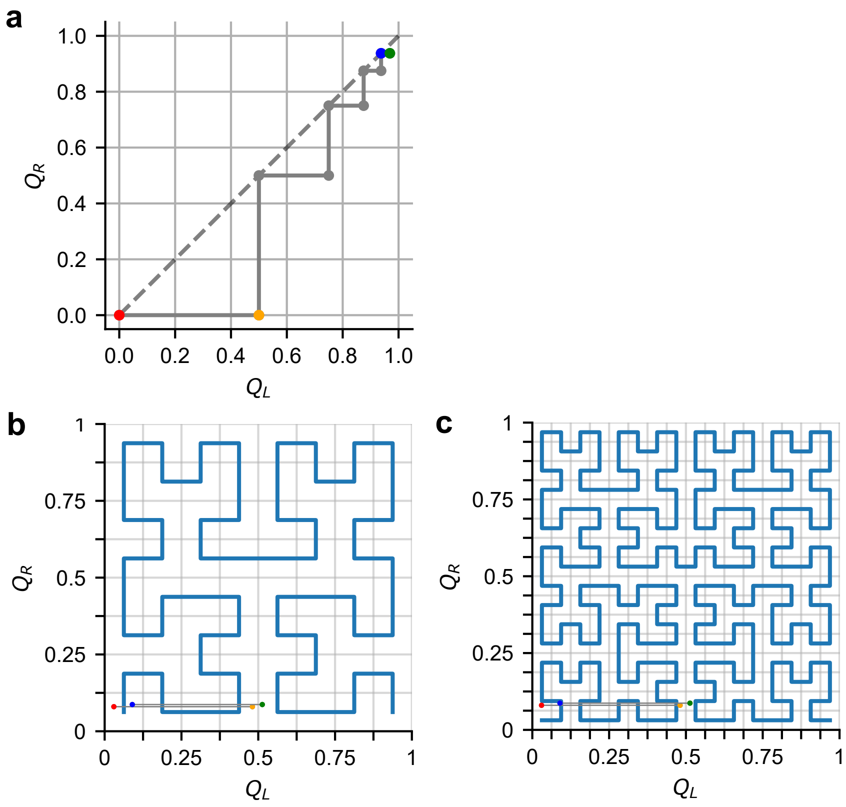


**Fig. S40.** Illustration of two-dimensional state space of the model-free RL model, defined by the action values $Q_L$ and $Q_R$. Upon receiving [$A_L$ R=1], the red-point state transits to the orange-point state, while the blue-point state transits to the green-point state. (a) In the one-dimensional RNN, the readout from the hidden state to the output layer (before softmax) is linear. (b-c) In the one-dimensional RNN, the readout from the hidden state to the output layer (before softmax) can take an arbitrarily nonlinear form. Blue curves represent the (b) third and (c) fourth iterations of the Hilbert curves, representing the one-dimensional hidden state $h$ of the RNN.

#### References


[1] Manuel Molano-Mazón, Yuxiu Shao, Daniel Duque, Guangyu Robert Yang, Srdjan Ostojic, and Jaime de la Rocha. Recurrent networks endowed with structural priors explain suboptimal animal behavior. *Current Biology*, 2023.

[2] Pamela Lyon and Franz Kuchling. Valuing what happens: a biogenic approach to valence and (potentially) affect. *Philosophical Transactions of the Royal Society B*, 376(1820):20190752, 2021.

[3] Thomas Akam, Ines Rodrigues-Vaz, Ivo Marcelo, Xiangyu Zhang, Michael Pereira, Rodrigo Freire Oliveira, Peter Dayan, and Rui M Costa. The anterior cingulate cortex predicts future states to mediate model-based action selection. *Neuron*, 109(1):149–163, 2021.

[4] Jane X Wang, Zeb Kurth-Nelson, Dharshan Kumaran, Dhruva Tirumala, Hubert Soyer, Joel Z Leibo, Demis Hassabis, and Matthew Botvinick. Prefrontal cortex as a meta-reinforcement learning system. *Nature neuroscience*, 21(6):860–868, 2018.

[5] Safa Alver and Doina Precup. What is going on inside recurrent meta reinforcement learning agents? *arXiv preprint arXiv:2104.14644*, 2021.

[6] Vladimir Mikulik, Grégoire Delétang, Tom McGrath, Tim Genewein, Miljan Martic, Shane Legg, and Pedro Ortega. Meta-trained agents implement bayes-optimal agents. *Advances in neural information processing systems*, 33:18691–18703, 2020.

[7] Jay Hennig, Sandra A. Romero Pinto, Takahiro Yamaguchi, Scott W. Linderman, Naoshige Uchida, and Samuel J. Gershman. Emergence of belief-like representations through reinforcement learning. *bioRxiv*, 2023. doi: 10.1101/2023.04.04.535512. URL https://www.biorxiv.org/content/early/2023/04/07/2023.04.04.535512.

[8] Hans Sagan. *Space-filling curves*. Springer Science & Business Media, 2012.

[9] Steven H Strogatz. *Nonlinear dynamics and chaos: with applications to physics, biology, chemistry, and engineering*. CRC press, 2018.
